# Flipping the odds of drug development success through human genomics

**DOI:** 10.1101/170142

**Authors:** Aroon D. Hingorani, Valerie Kuan, Chris Finan, Felix A. Kruger, Anna Gaulton, Sandesh Chopade, Reecha Sofat, Raymond J. MacAllister, John P. Overington, Harry Hemingway, Spiros Denaxas, David Prieto, Juan Pablo Casas

**Author notes:** equal contribution. Correspondence to: Aroon Hingorani.

## Abstract

Drug development depends on accurately identifying molecular targets that both play a causal role in a disease and are amenable to pharmacological action by small molecule drugs or bio-therapeutics, such as monoclonal antibodies.

Errors in drug target specification contribute to the extremely high rates of drug development failure.

Integrating knowledge of genes that encode druggable targets with those that influence susceptibility to common disease has the potential to radically improve the probability of drug development success.

## Part 1: System flaws in drug development

> ‘***The greatest obstacle to discovery is not ignorance* – *it is the illusion of knowledge’***
>
> — - **Attributed to Daniel J. Boorstin (Historian, 1914-2004).**

### Background

The patent and drug regulatory systems encourage innovation by rewarding risky but potentially transformative research and development (R&D). However, since 96% of drug development programmes currently fail^1 2^, the imbalance between risk and reward in the pharmaceutical sector has led to a range of undesirable consequences.

Chief among these is the inflationary pressure on drug prices. This is imposed by the need to recoup the incurred cost of historical failures through any development successes, so as to continue to provide shareholders with a return on their investment^3^. This cost is borne by healthcare systems and transferred to citizens via health insurance premiums or taxation.

All too frequently, high-profile failures of anticipated ‘blockbuster’ or ‘niche-buster’^4^ drugs lead pharmaceutical companies to restructure and refocus in-house R&D, leading to job losses, site closures, off-shoring, or mergers and acquisitions, aimed at containing cost and supporting the company share price in the short to mid-term^5 6 7 8^. Small and medium sized companies (SMEs) in the biotech sector, alongside increased public funding of academic translational research^9^, absorb some of the early stage R&D risk. However, the interest of these organisations may be less in the ultimate therapeutic success of a new drug and more in its value as an asset-with-prospects. Value is often added by incremental (rather than definitive) preclinical or early clinical phase proof-of-concept studies, before the compound, know-how and patent for a disease indication is then licensed to the next developer in the chain, and so on. Under this model, no single organisation has an end-to-end capability or responsibility for taking a potential treatment from concept to licence.

With high risk and infrequent reward, R&D can become misdirected from the innovative to the derivative ^10^. This is because both the patent and regulatory systems are vulnerable to some element of gaming. New compounds with identical mechanisms of action (so called ‘me-too treatments’), and minor changes in formulation (e.g. the separation of the pharmacologically active stereoisomer from an already effective racemic mixture, slow-release delivery vehicles for existing drugs, and new combinations of old drugs) can occasion a new license and, in effect, the same level of patent protection as a drug with a truly novel mechanism of action. Sometimes, patients reap real benefit from the improved compound or formulation. More often, the process is simply a means for companies to extend patent life (ever-greening) ^11^.

However, healthcare providers are now raising the therapeutic bar, such that even newly licensed drugs cannot be guaranteed to capture a market share sufficient to recoup R&D costs, unless they demonstrate a genuine cost-effective advance over existing therapies^12 13^.

In response, governments, who are conflicted in their need to ensure cost-efficient healthcare on the one hand, but to support the pharmaceutical sector as a major employer and taxpayer on the other, have explored schemes to reduce barriers to market access. Examples include the breakthrough designation scheme in the US^14^, the priority medicines scheme (PRIME) in Europe^15^, and the Early Access to Medicines scheme in the UK^16^. However, the success of such initiatives is reliant on truly innovative and transformative products emerging efficiently from pharmaceutical R&D pipelines, which has not been the experience of the last few decades.

As a consequence, the economic sustainability of the current model of drug development has been questioned and calls made for some form of disruptive solution to improve both scientific and market efficiency, and to fuel innovation^17 18 19^.

### Reasons for the high drug development failure rate

To understand how drug development efficiency could be improved, it is necessary to understand the reasons for failure. **Box 1** summarises the process of drug development.

#### Box 1. The process of drug development^20^

Developing a drug with a new mechanism of action requires fulfilling a series of tasks in sequence:

1. Selecting a disease for which there is a deficit in existing therapies;
2. Identifying a pathogenic mechanism and potential drug target (almost all of which are proteins);
3. Screening for and optimising a compound (sometimes a small molecule or, increasingly, a monoclonal antibody or peptide) that specifically modulates the function of the target protein, is free of toxicity and has the desired pharmacokinetic properties;
4. Demonstrating target engagement by the compound (through the use of biomarkers or surrogate measures of the disease process); and,
5. Demonstrating efficacy against the disease end-point in tandem with an adequate safety profile.

Operationally, this is achieved in two stages: preclinical and then clinical. Preclinical studies utilise isolated cells, organoid cultures, tissue preparations *ex vivo*, and (if available) animal models of human disease. They test the hypothesis that the selected target plays a controlling role in the disease of interest (proof of concept) and that the compound has an adequate safety profile. If preclinical studies are encouraging, a critical decision is made to progress to clinical evaluation. This is initially through healthy volunteer studies for pharmacokinetics, dose finding and tolerability (Phase 1); and then exposure of a small number of patients often evaluating surrogate measures of disease (Phase 2). If these studies appear promising, a larger randomised (Phase 3) outcome trial will follow, typically 10 or more years after programme initiation, following several hundred million pounds of investment.

During the lengthy development process, there is relentless attrition of programmes and products. Even for compounds reaching clinical phase, only around 10% of entrants emerge as licensed drugs.^1 2 21^ The key productivity-limiting obstacle turns out to be ‘late-stage failure’ during phase 2 or phase 3 randomised trials^22^. This has major consequences, particularly for smaller pharmaceutical companies with a thin therapeutic pipeline and limited financial resources to absorb such failures.

But why is late-stage failure a recurrent problem? Two decades ago, unfavourable pharmacokinetics was the most frequent single cause of clinical phase attrition^23^. By a decade later, this problem had largely been resolved such that two thirds of late-stage failures of first-in-class compounds can now be attributed to a different problem: lack of efficacy in the intended disease, despite adequate engagement of the target protein and apparently favourable signals from preclinical and early phase clinical studies.^24 25 26 27 28^. *Thus*, *most late*-*stage failures now occur because the target turns out not to play the causal role in the disease that was hypothesised at the outset*. Late-stage failure for lack of efficacy therefore exposes a critical problem in drug development: matching the correct drug targets to each disease. The established system of drug development has been poor at this crucial task because of two key system flaws.

### First system flaw: preclinical studies are unreliable predictors of development success

Preclinical studies in cell culture systems, tissues, isolated organs and animal models that are widely used for drug target identification (and validation) have a range of acknowledged limitations^29^. Cells provide an incomplete picture of responses in tissues, which are composed of a wide range of interacting cell types. In turn, responses in whole organs *ex vivo* may not reflect the response of the whole animal. Experiments in animals may be poorly representative of responses in humans because of species differences in pathophysiology, while some animal disease models may be an artifice of the human disorder^30 31 32^. Concerns are also now being raised that most (perhaps >90%)^33^ of the nominally positive preclinical research studies undertaken in academia (perhaps in industry too), and which sometimes seed a drug development programme, are often not only poorly representative of human pathophysiology but are also frequently irreproducible. Investigating the causes of irreproducibility is becoming an area of funded research^34^. Reasons for irreproducibility encompass data selection to flatter or overestimate any real effect, and flaws in experimental design, including the failure to routinely randomise experimental interventions, and to blind the assessment of outcome. A pervasive cause of irreproducibility occurs from errors of statistical inference arising from common misconceptions about P values, including confusion between significance and hypothesis testing^35 36^, which contributes to high rates of false discovery^37^. **Box 2** expands on the reasons for the high false discovery rate in biomedical research.

#### Box 2. False discovery rate (*FDR*) in biomedical research

A frequent misconception in biomedical research is that the false discovery rate (*FDR*) and the Type 1 (false positive) error rate (*α*) are equivalent ^37, 38^. The reason this is not the case is illustrated by a hypothetical example. Imagine a field of study in which experiments are undertaken with robust design: all interventions are allocated at random and, in each experiment, the estimated treatment effect has informed the sample size such that the experimental false positive error rate (*α*) is 0.05 and the Type 2 (false negative) error rate (*β*), is 0.2. The power, (1 – *β*), which can be conceptualised as the detection rate for a real effect, is therefore 0.8. We introduce a third parameter (*γ*), the proportion of true relationships out of all those tested in the field. In the current illustration, we assume *γ* = 0.1. **Table 1a** illustrates that, despite the robust experimental design, these parameters dictate that 36% (not 5%) of nominally positive experimental outcomes are false discoveries. In general, *FDR* is related to *α*, *β* and *γ* as follows:

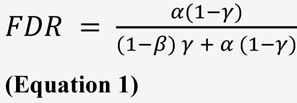

**Table 1b and Table 2** demonstrate how *FDR* varies at different values of *α*, *β* and *γ.* Reducing *α* has the effect of reducing *FDR.* Increasing *β* (equivalent to reducing power, e.g. from 0.8 to 0.2, which is close to the mean power recently found in a survey of preclinical studies in the field of neuroscience)^39^ increases *FDR* (from 36% to 69% in this example, so that false discoveries would then outnumber true discoveries by about 2:1). *FDR* increases as the proportion of true relationships (*γ*) decreases. In addition, it is not widely appreciated that real effects, even when present can be overestimated by small studies, because a positive finding must be extreme for it to exceed the usual experimental significance threshold (a similar notion to small study bias in clinical trials, and the winner’s curse^40^).

**Table 1a.** The difference between the type 1 error (false-positive) rate (*α*) and the false-discovery rate (*FDR*). 1000 different hypotheses in a field are tested by experiments designed with a detection rate (power; 1 – *β*) = 0.8, with *α* = 0.05. With 100 real effects to discover (*γ* = 0.1), the false discovery rate is 45/125 = 36%.

|  | True relationship | No relationship | Hypotheses tested | *TDR* | *FDR* |
| --- | --- | --- | --- | --- | --- |
| Observed relationship | 80 | 45 | 125 | 0.64 | 0.36 |
| No observed relationship | 20 | 855 | 875 |  |  |
| Total | 100 | 900 | 1000 |  |  |

**Table 1b.** The relationship between *α*,*β*, and *γ*, the true discovery rate (*TDR*) and the false discovery rate (*FDR*).

| Outcome | Causal pairings | Non-causal pairings | Hypotheses tested | *TDR* | *FDR* |
| --- | --- | --- | --- | --- | --- |
| Declared positive | $\gamma(1 - \beta)$ | $\alpha(1 - \gamma)$ | $[\gamma(1 - \beta)] + [\alpha(1 - \gamma)]$ | $\frac{\gamma(1 - \beta)}{\gamma(1 - \beta) + \alpha(1 - \gamma)}$ | $\frac{\alpha(1 - \gamma)}{(1 - \beta)\gamma + \alpha(1 - \gamma)}$ |
| Declared negative | $\gamma\beta$ | $(1 - \alpha)(1 - \gamma)$ | $[\gamma\beta] + [(1 - \alpha)(1 - \gamma)]$ | | |
| | $\gamma$ | $1 - \gamma$ | 1 | | |

**Table 2:**
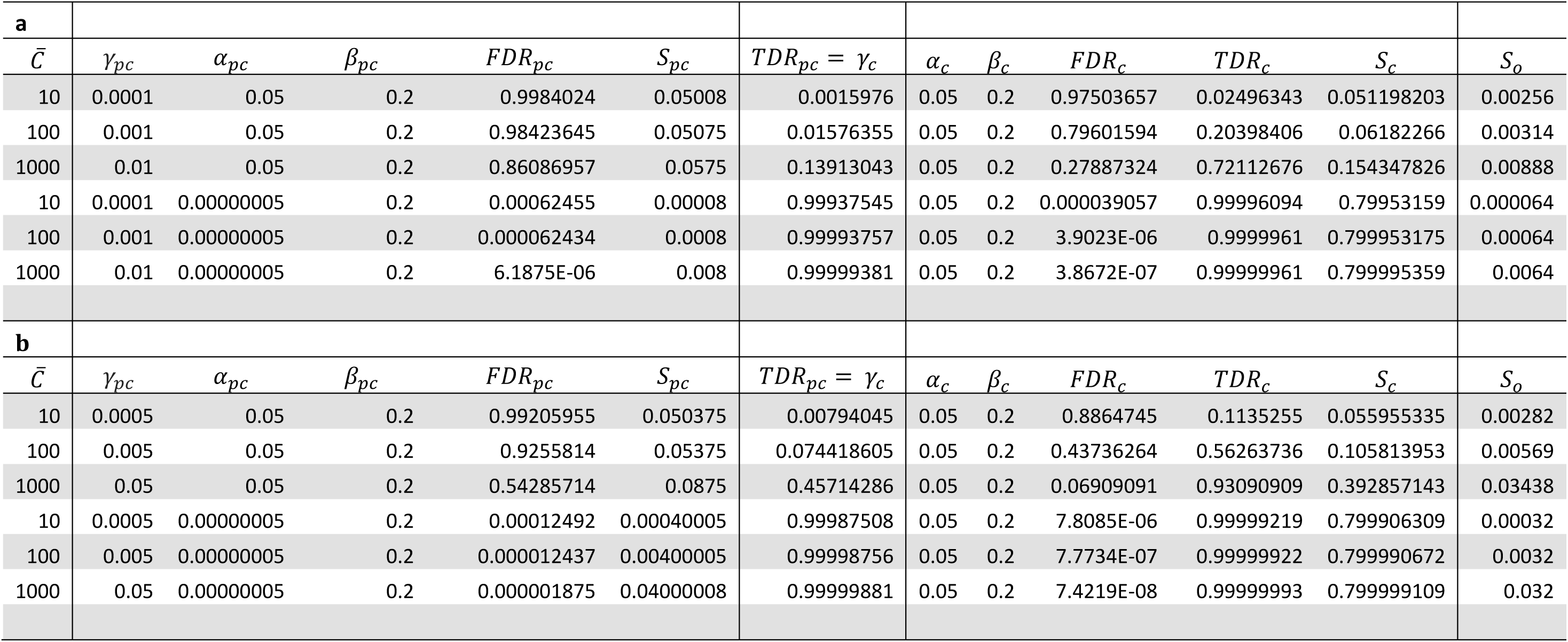
*A priori* estimates of preclinical (*pc*), clinical (*c*) and overall (*o*) drug development success contrasting orthodox (non-genomic) with genomic approaches. *TDR*, *FDR*, *S_pc_*,*S_C_* and *S_o_* are presented at different values of *α* (Type 1 error rate) *β* (Type 2 error rate) and *γ* (proportion causal and druggable targets). *γ_pc_* = (*^C^*/*N_g_*) (*^N^_T_*/*N_G_*) when the sample space is defined by **a)** *N_G_* × *N_D_*, and **b)** when the sample space is restricted to the druggable genome (*N_G_* × *N_T_*). See text for details.

| **a** |  |  |  |  |  |  |  |  |  |  |  |  |
| --- | --- | --- | --- | --- | --- | --- | --- | --- | --- | --- | --- | --- |
| $\bar{C}$ | $\gamma_{pc}$ | $\alpha_{pc}$ | $\beta_{pc}$ | $FDR_{pc}$ | $S_{pc}$ | $TDR_{pc} = \gamma_c$ | $\alpha_c$ | $\beta_c$ | $FDR_c$ | $TDR_c$ | $S_c$ | $S_o$ |
| 10 | 0.0001 | 0.05 | 0.2 | 0.9984024 | 0.05008 | 0.0015976 | 0.05 | 0.2 | 0.97503657 | 0.02496343 | 0.051198203 | 0.00256 |
| 100 | 0.001 | 0.05 | 0.2 | 0.98423645 | 0.05075 | 0.01576355 | 0.05 | 0.2 | 0.79601594 | 0.20398406 | 0.06182266 | 0.00314 |
| 1000 | 0.01 | 0.05 | 0.2 | 0.86086957 | 0.0575 | 0.13913043 | 0.05 | 0.2 | 0.27887324 | 0.72112676 | 0.154347826 | 0.00888 |
| 10 | 0.0001 | 0.00000005 | 0.2 | 0.00062455 | 0.00008 | 0.99937545 | 0.05 | 0.2 | 0.000039057 | 0.99996094 | 0.79953159 | 0.000064 |
| 100 | 0.001 | 0.00000005 | 0.2 | 0.000062434 | 0.0008 | 0.99993757 | 0.05 | 0.2 | 3.9023E-06 | 0.9999961 | 0.799953175 | 0.00064 |
| 1000 | 0.01 | 0.00000005 | 0.2 | 6.1875E-06 | 0.008 | 0.99999381 | 0.05 | 0.2 | 3.8672E-07 | 0.99999961 | 0.799995359 | 0.0064 |
| **b** |  |  |  |  |  |  |  |  |  |  |  |  |
| $\bar{C}$ | $\gamma_{pc}$ | $\alpha_{pc}$ | $\beta_{pc}$ | $FDR_{pc}$ | $S_{pc}$ | $TDR_{pc} = \gamma_c$ | $\alpha_c$ | $\beta_c$ | $FDR_c$ | $TDR_c$ | $S_c$ | $S_o$ |
| 10 | 0.0005 | 0.05 | 0.2 | 0.99205955 | 0.050375 | 0.00794045 | 0.05 | 0.2 | 0.8864745 | 0.1135255 | 0.055955335 | 0.00282 |
| 100 | 0.005 | 0.05 | 0.2 | 0.9255814 | 0.05375 | 0.074418605 | 0.05 | 0.2 | 0.43736264 | 0.56263736 | 0.105813953 | 0.00569 |
| 1000 | 0.05 | 0.05 | 0.2 | 0.54285714 | 0.0875 | 0.45714286 | 0.05 | 0.2 | 0.06909091 | 0.93090909 | 0.392857143 | 0.03438 |
| 10 | 0.0005 | 0.00000005 | 0.2 | 0.00012492 | 0.00040005 | 0.99987508 | 0.05 | 0.2 | 7.8085E-06 | 0.99999219 | 0.799906309 | 0.00032 |
| 100 | 0.005 | 0.00000005 | 0.2 | 0.000012437 | 0.00400005 | 0.99998756 | 0.05 | 0.2 | 7.7734E-07 | 0.99999922 | 0.799990672 | 0.0032 |
| 1000 | 0.05 | 0.00000005 | 0.2 | 0.000001875 | 0.04000008 | 0.99999881 | 0.05 | 0.2 | 7.4219E-08 | 0.99999993 | 0.799999109 | 0.032 |

Many previous discussions of the extent of the *FDR* problem have been somewhat abstract in nature. But is it possible to estimate real-world *FDR*, and, if so, to compute the impact on drug development success rates?

By setting some simplifying assumptions and approximating certain parameters, we now estimate *FDR* for preclinical studies that usually provide a start point for drug development.

Understanding disease aetiology can frequently be distilled to understanding which of the proteins encoded in the genome plays a controlling or causal role in each disease process. Drug targets are also almost exclusively proteins. We therefore introduce the following:

**Assumption 1:** Each gene encodes a unique protein with a single function
**Assumption 2:** A given protein can influence the risk of more than one disease
**Assumption 3:** The probability of a protein influencing the pathogenesis of one disease is independent of the probability that it influences any other

We recognise that these assumptions, as well as others we will introduce in due course, represent very substantial oversimplifications, and many exceptions can be identified from current drugs and diseases. However, they can also help to estimate certain ‘base-case’ probabilities. Later in this article we dissect these assumptions, as well as others we introduce later, and explore the impact of any modifications on the base-case probabilities.

The key parameters needed for the estimation of *FDR* in biomedical research are the number of human diseases of interest; the number of protein coding genes; and the average number of proteins that are likely to play a causal role in any given disease.

Taking the complexities and inaccuracies of disease definition into account (see **Box 3** and **Table 3** for details), we assume, as a start point, that the number of complex (multifactorial) diseases is close to 10,000, and that the number of human protein coding genes^41^ is around 20,000 (**Figure 1**). **Box 4** provides a historical overview of the route to establishing this estimate.

**Table 3:** The number of terms within widely used disease classification systems and ontologies as of 24 February 2016.

| **Coding Scheme** | **Type** | **Number of terms** | **Data source** |
| --- | --- | --- | --- |
| **ICD-10** | Disease classification | 12,445 | http://apps.who.int/classifications/apps/icd/ClassificationDownload/DLArea/Download.aspx |
| **Human Disease Ontology** | Ontology | 9,196 | https://github.com/DiseaseOntology/HumanDiseaseOntology/tree/master/src/ontology |
| **Human Phenotype Ontology** | Ontology | 11,683 | http://human-phenotype-ontology.github.io/downloads.html |
| **Experimental Factor Ontology** | Ontology | 17,263 | https://sourceforge.net/p/efo/code/HEAD/tree/trunk/src/efoinobo/efo.obo |
| **Expanded Diagnostic Cluster** | Disease groups | 282 | The Johns Hopkins ACG® System Version 11.0 Technical Reference Guide |
| **Clinical Classification Software** | Disease groups | 259 | http://www.ahrq.gov/research/data/hcup/icd10usrgd.html |
| **PheWAS Catalog** | Disease groups | 1,645 | https://phewas.mc.vanderbilt.edu/ |
| **SNOMED CT** | Clinical Terminology | 422,382 | https://www.nlm.nih.gov/research/umls/licensedcontent/snomedctfiles.html |
| **READ CTV3** | Clinical Terminology | 329,147 | https://isd.hscic.gov.uk/trud3/user/guest/group/0/pack/9 |

**Figure 1.**
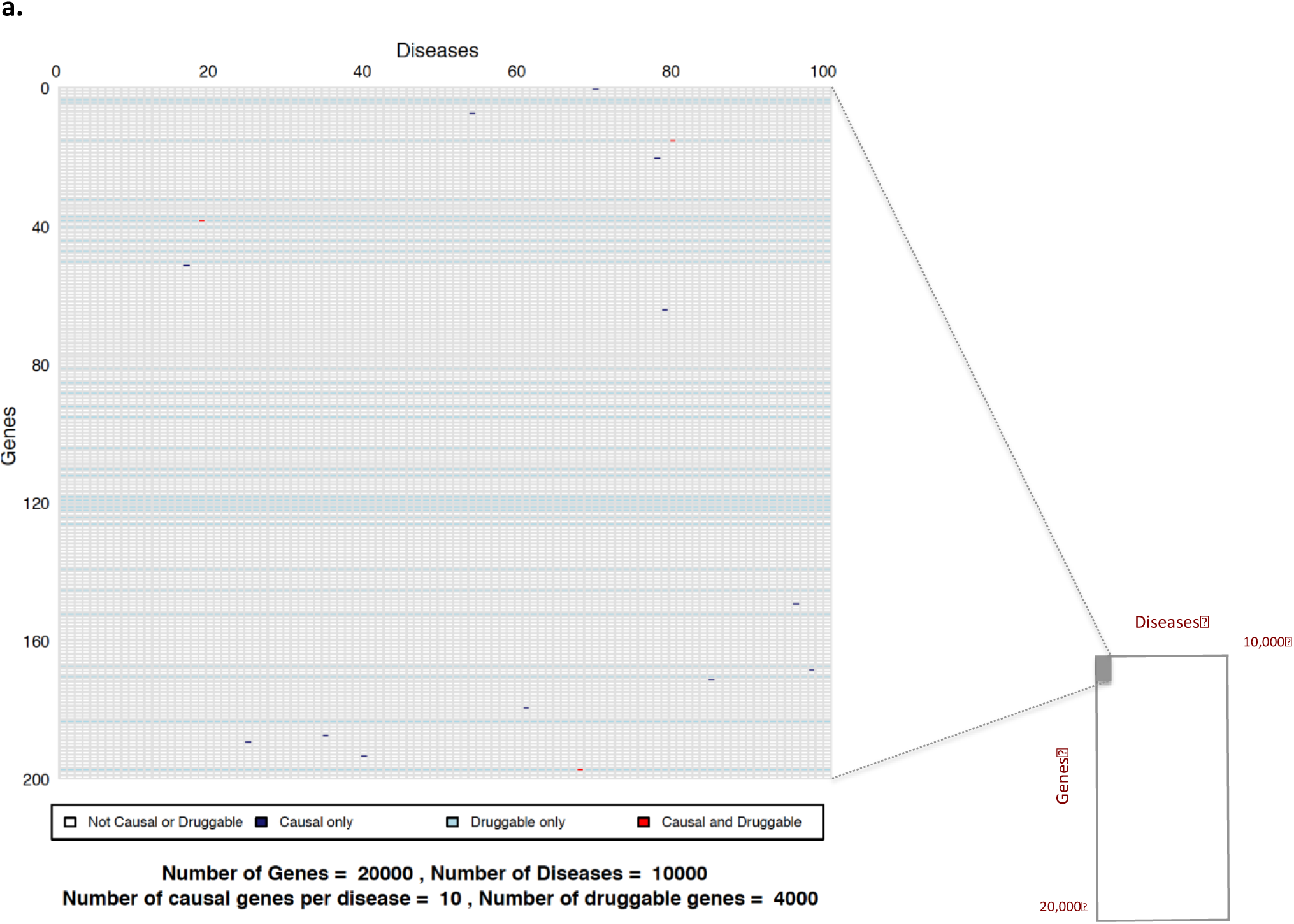
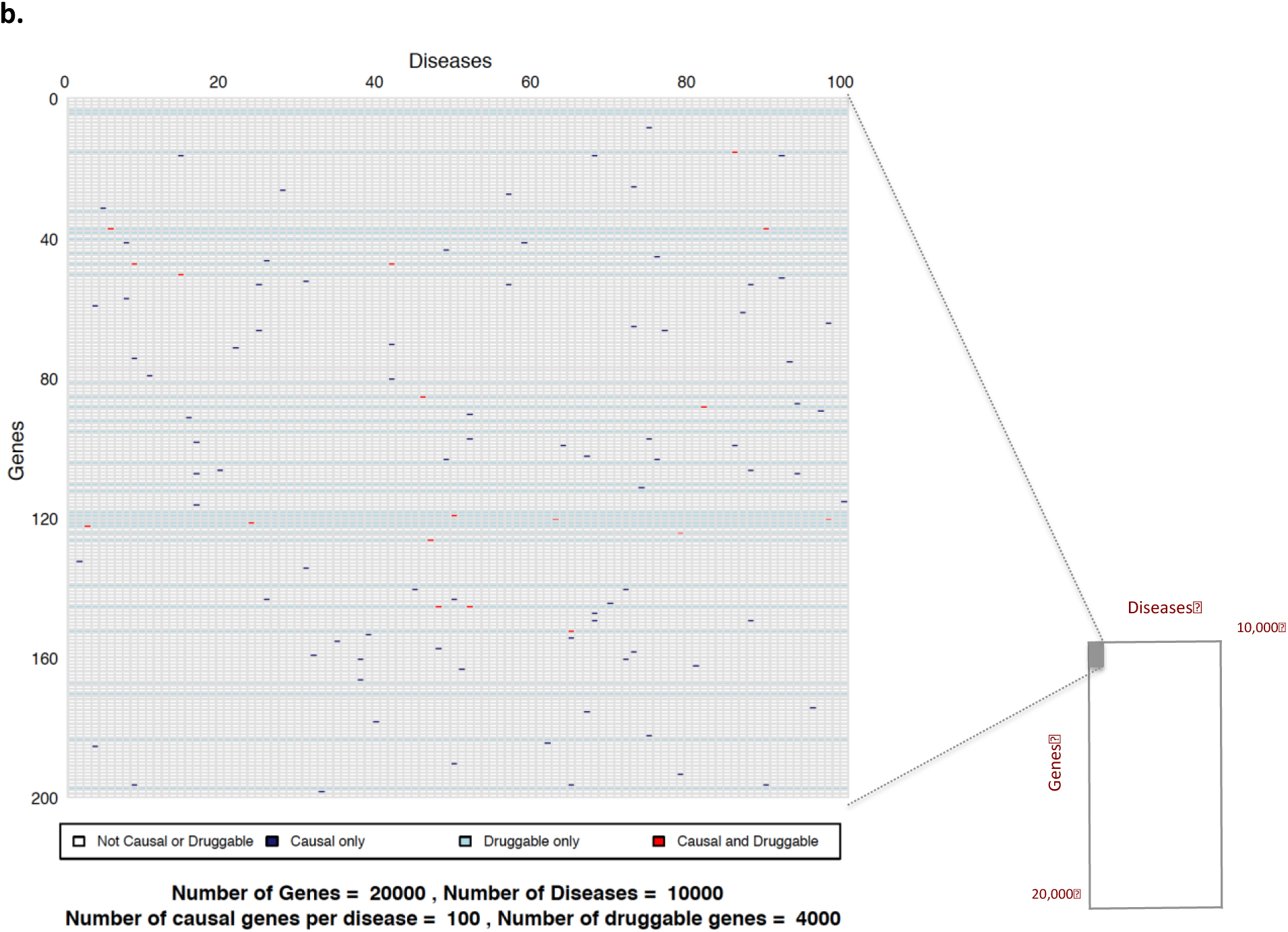
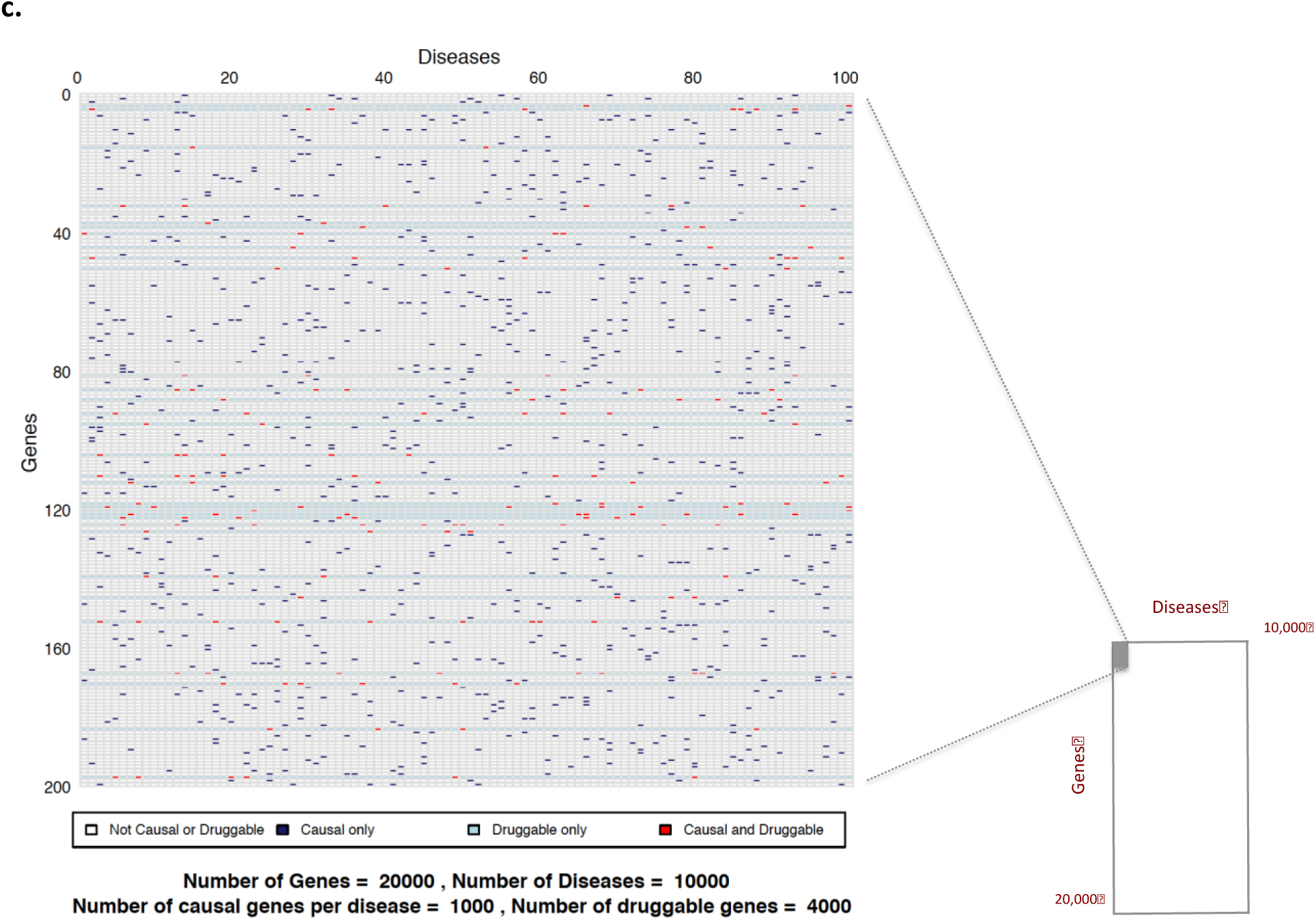
Sample space defined by 10,000 human diseases (columns) and 20,000 proteincoding genes (rows). Expanded region comprising 1/10,000*th* of the whole sample space is enlarged: **a** (based on 10 causative genes per disease); **b** (based on 100 causative genes per disease); and **c** (based on 1000 causative genes per disease). Each cell represents a unique gene-disease pairing. Dark blue cells indicate causal gene-disease pairings, light blue cells druggable gene-disease pairings, with red cells indicating causal and druggable gene disease pairings.

**Figure 2a.**
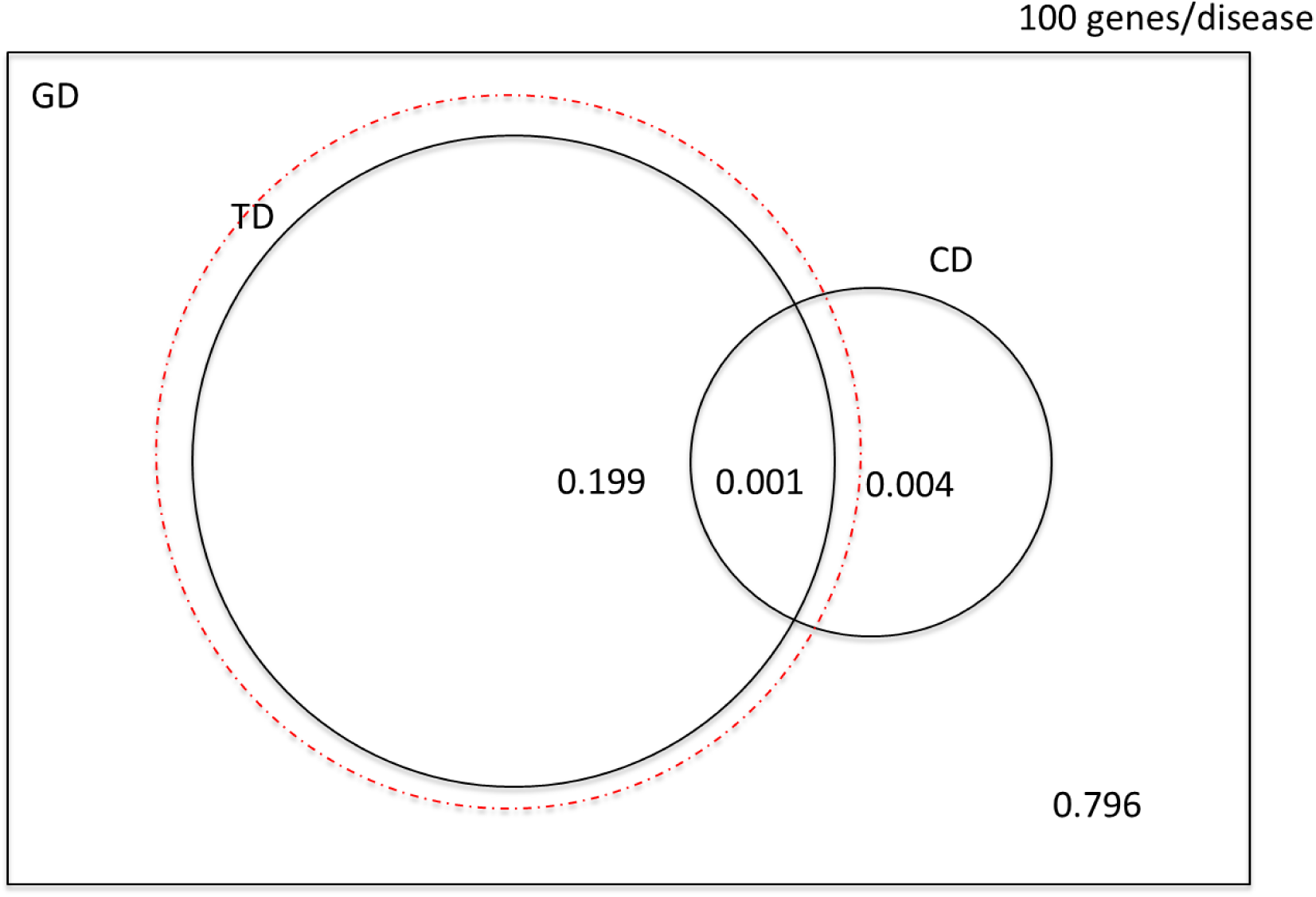
Venn diagram illustrating the probabilities of selecting a causal, druggable gene-disease pair (*CD* ∩ *TD*), a druggable gene disease pair (*TD*) and a causal, gene disease pair (*CD*) from a sample space of 200 *x* 10^6^ gene disease pairings, 100 causal genes per disease and 4000 druggable genes from the 20,000 in the genome. The dashed red circle encloses a probability space restricted to druggable genes. (Not to scale).

**Figure 2b.**
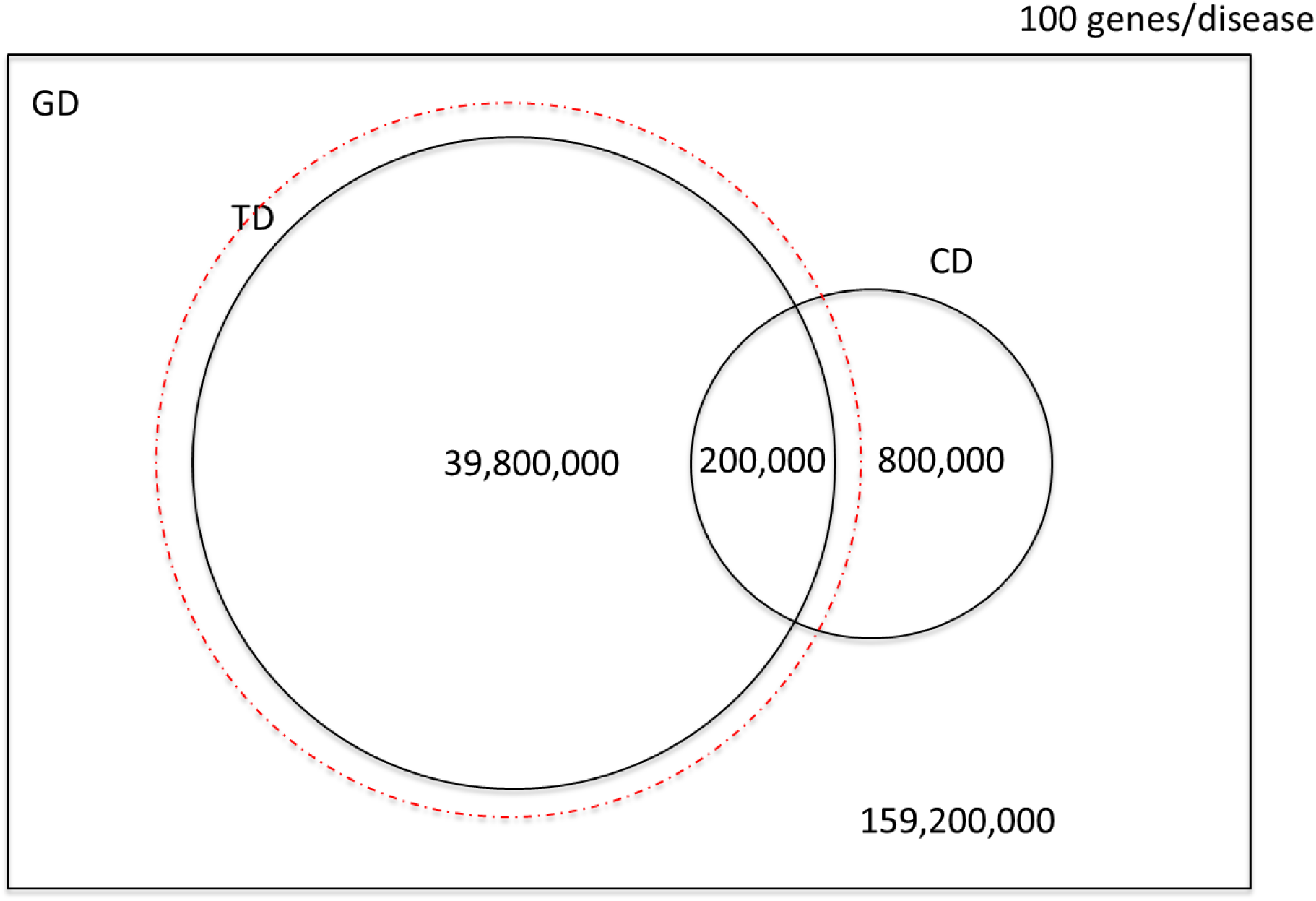
Venn diagram illustrating the number of causal, druggable gene-disease pairs (*CD* ∩ *TD*), druggable gene disease pairs (*TD*) and causal, gene disease pairs (CD) from a sample space of 200 *x* 10^6^ gene disease pairings, 100 causal genes per disease and 4000 druggable genes from the 20,000 in the genome. The dashed red circle encloses a probability space restricted to druggable genes. (Not to scale).

#### Box 3. Estimating the number of human disease entities

Estimating the exact number of human diseases is a surprisingly challenging task. Clinical priorities have led to definitions of disease that rely on characteristic clusters of symptoms and signs supported to a varying degree by biophysical, laboratory, radiological or histological tests that detect abnormalities of structure or function. Defining disease on the basis of manifestations rather than cause means that diagnoses may be remote from the molecular mechanisms leading to disease, many of which remain unknown. In this paper, we set aside rare monogenic conditions, focusing instead on common (multifactorial) human diseases of potential therapeutic interest that have both a genetic and environmental contribution. A list of medical coding schemes covering such diseases, from clinical terminologies to disease classification systems, is shown in **Table 3**. Standard vocabularies of medical terms such as SNOMED CT (Systematised Nomenclature of Medicine - Clinical Terms) which includes Read Clinical Terms Version 3 (CTV3), which are used in electronic health records, capture clinically relevant data related to individuals and their care. The difficulty with using these vocabularies to enumerate diseases is that multiple codes can refer to a single disease, both because of duplicate terms (largely rectified in SNOMED CT) and the hierarchical nature of these vocabularies. In addition, disease diagnoses comprise only a proportion of the descriptive terms, with many covering symptoms, procedures, treatments, drugs and healthcare administration. The International Classification of Diseases (ICD) is widely regarded as the authoritative classification system for causes of death and illnesses. Its use in recent revisions has been broadened to medical records indexing and reimbursement.

Approximately 4,000 of over 12,000 classes in the tenth revision, ICD-10, refer to health administration and external causes of morbidity and mortality and their consequences. Of the more than 8,000 remaining classes, (fewer than 500 of which are specific for rare diseases)^42 43^, overlaps occur within the hierarchical coding structure, such that a particular disease may be described by several codes. The same is true of disease and phenotype ontologies. Categorisation schemes such as the Clinical Classification Software developed by the US Agency for Healthcare Research and Quality (AHRQ), the Expanded Diagnostic Clusters (EDC) developed at Johns Hopkins University and the PheWAS Catalog designed at Vanderbilt University, collapse ICD codes into a smaller number of clinically meaningful categories that can be useful for presenting descriptive statistics.

#### Box 4. Estimating the number of protein coding genes in the human genome

As summarised by Pertea and Salzberg^44^, estimates of the number of human protein-coding genes have been revised progressively downward since the early 1960s. Very early estimates, predating the first draft of the human genome by around 40 years, were based on extrapolation from emerging information on the amino acid sequences of proteins^45^, or theoretical considerations^46^. When the human genome project was at its planning stage, the number of human genes was projected to stand at 50-100,000 (National Institutes of Health/Department of Energy report on the Human Genome Project). However, when the initial results emerged, the estimate was revised to around 25-30,000 genes^47^. With more exhaustive sequencing of the genome and its transcripts, more detailed annotation of sequence, comparative analysis of proteomic and sequence data, and the construction of a tissue based map of the human proteome^48^, the consensus estimate of the number of protein coding genes has fallen yet again^49^. Summary statistics on the human genome are now regularly updated by the GENCODE project. The resource has catalogued a consensus value for the number of human genes since 2009, at which time 22,250 protein-coding genes were listed. In the latest data freeze (March 2016, Version 25), the number of genes listed is 19,950.

To estimate the average number of protein-coding genes that play a causal role in any given disease, we draw on experience from previous genome wide association studies (GWAS; **see Box 5**). This is the only routinely used study design that estimates the influence of every gene (and protein) on a disease systematically. The ability to detect disease-causing genes differs from one GWAS to the next, depending both on the underlying genetic effect in the disease of interest and the available sample size. We therefore confine our consideration to those GWAS and meta-analysis of GWAS (meta-GWAS) with the very largest sample sizes. Examples of such meta-GWAS include inflammatory bowel disease (60,000 individuals studied; 99 loci identified)^50^, type 2 diabetes (150,000 individuals; 150 loci)^51^, and coronary heart disease (200,000 individuals; 46 loci)^52^. Thus, each of these meta-GWAS has identified in the order of 100 susceptibility loci per disease. The number of disease-associated loci may not equate precisely to the number of causal genes per disease, and it may also be anticipated that yet larger sample sizes will yield yet more loci, because much of the heritability of common disorders remains unexplained^53^. There is also a school of thought that all genes (and proteins) play some role in all diseases – the infinitesimal^54^ or omnigenic^55^ model – which we discuss in more detail later. However, with these caveats, let us assume, initially, that there are 100 causal genes per disease on average.

We now define the following:

{*G*} is the set of protein – coding genes
{*D*} is the set of common human diseases
{*GD*} is the set of all possible gene – disease pairs
{*C*} is the set of causal genes for a given disease
{*CD*} is the set of all causal gene – disease pairs

*N_G_* = Total number of protein – coding genes = 20,000
*N_D_* = Total number of complex human diseases = 10,000
*N_GD_* = Total number of possible gene - disease pairs = 10,000 × 20,000 = 200 × 10^6^
*C* = the number of causal genes in a given disease
*C̄* = the average number of causal genes per disease = 100
*N_CD_* = Total number of causal gene – disease pairs = 100 × 10,000 = 1 × 10^6^

Based on assumptions 1-3, the probability (*P_C_*) that any gene- (or, equivalently, any protein)-disease pairing selected at random from the set of all possible gene-disease pairs {*GD*} also belongs to the set of causal gene-disease pairs {*CD*} is given by:

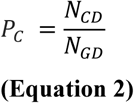

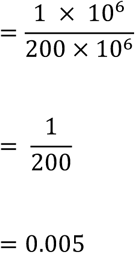

This can also be written as:

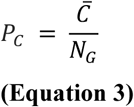

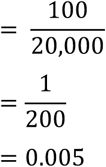

*P_C_* =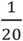 if *C* =1000, but *P_C_* falls to 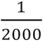 if *C* = 10.

As follows from **Equation 3**, *P_C_* is independent of the number of diseases under consideration, as long as *C* is constant. As an illustration, focusing on 5000 diseases (rather than 10,000) would shrink the sample space by half to 5000 × 20,000 (= 100 × 10^6^) gene (protein)-disease-pairings, but would also reduce the number of causal gene (protein)-disease pairs in the sample space by the same proportion, from 1 × 10^6^ to 500,000.

Importantly, *P_C_* can also be interpreted as the proportion of true hypotheses for tests of causality amongst all possible gene-disease pairings, and can hence also be represented as *γ_C_* (**see Box 2**). In this case, *γ_C_* refers to the probability of a true causal gene-disease pairing occurring within the sample space {*GD*}. Therefore:

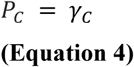

Let us now consider preclinical experiments designed such that *α* = 0.05, and a detection rate (power) for causal pairings (1 – *β*) = 0.8.

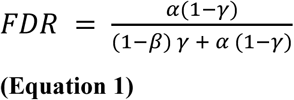

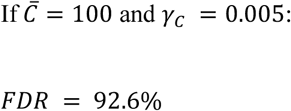

This *FDR* value for biomedical research is very close to that estimated previously by Ioannidis^33^.

However, scientists, it might be argued, do not select protein-disease pairings at random: they work on particular diseases and proteins that have been seemingly confidently paired on the basis of previous research. Scientists are also not generally interested in identifying a protein that is causal for *any* disease, but rather in identifying proteins contributing to the pathogenesis of a particular disease of interest, a point to which we return in a later section. But if, as Ioannidis and others have argued, there is strong empirical evidence from many research fields of extremely high rates of false discovery, leading to pervasive unreliability of the evidence base, then seemingly informed hypotheses may turn out to be spurious^56^. In Bayesian terms, the prior probability of correctly pairing a gene (or protein) with a disease may be close to that of the background probability of a success in a *random pick* from the sample space. The proportion of false discoveries in the medical literature could be inflated further because of the greater likelihood of positive than negative findings being submitted and accepted for publication^57^.

For now, in summary, preclinical research is poorly predictive of drug development success partly because of the poor external validity of cell, tissue and animal models, partly because of flaws in experimental design and significance chasing and publication bias, but perhaps mainly because of the pervasive *FDR* problem. This occurs because:

a. Preclinical studies are often too small to detect true positive associations because the actual power (1 – *β*) is lower than that pre-specified at the study design stage because of over-optimistic estimates of effect sizes: when real associations are detected, the effect sizes will be overestimated.
b. The usual experimental false positive rate (*α*) of 0.05 leads to an excess of false discoveries because;
c. Causally-relevant gene (or protein)-disease pairings (true disease hypotheses) in most areas of research are greatly outnumbered by the number of non-causal ones, that is the value of *γ_C_* tends to be small, often far below 0.1.

It is easy to envisage how these conditions could lead to drug development programmes being initiated on the basis of misleading preclinical research, progressing into the clinical phase of development only to stumble expensively at phase 2 or 3.

Expensive late-stage failure would appear to be an consequence of the high *FDR* in preclinical target validation studies. But is it avoidable?

Lessons can be learnt from the field of common disease genetics, which overcame the high *FDR* problem in the era of candidate gene association studies. Resolution was achieved through a complete re-examination of the way in which research in that field was conducted. As a consequence, genetic association studies now yield some of the most reproducible findings in any field of biomedicine, detecting loci throughout the genome influencing a wide range of diseases and biomarkers^58^. The steps taken to rescue common disease genetics from the epidemic of false discoveries in the ‘candidate gene era’ are summarized in **Box 5**^59^.

#### Box 5. Resolution of the high false discovery rate problem in the field of common disease genetics

Three major factors contributed to the resolution of the high *FDR* problem in the field of common disease genetics in the candidate gene era. These were:

a. The development of fixed content genotyping arrays that, to a first approximation, could interrogate all genes in a genome, not just a subset of them, triggering the move from candidate gene to whole-genome (genome-wide) association studies (GWAS);
b. Recognition that a much more stringent a-value threshold would be needed in such studies to minimize false discoveries, as can be observed from **Table 2**, where changing *α* from 0.05 to 5 × 10^−8^ (the now widely used genome wide Type I error rate) reverses *TDR* and *FDR*
c. Understanding that larger sample sizes than had been usual up to that time would be needed to retain power in the context of the much stricter α-value threshold. As a consequence, clinicians and scientists began to assemble large collections of patients with diseases of interest (and controls) and, by necessity, to work together in consortia to achieve datasets of the necessary size, pooling information from individual studies in a statistically robust way using meta-analysis, a technique which, by then, had already become well-established in the clinical trial setting. A GWAS incorporating data from over 200,000 individuals by meta-analysis would now be viewed as unexceptional. The findings from GWAS are curated by a number of repositories ^60 61^ including the NHGRI-EBI GWAS catalog at https://www.ebi.ac.uk/gwas/.

Yet, while the problem of high false discovery rates has led to a root and branch change in the field of complex disease genetics, a similar transformation is yet to take place in preclinical laboratory science that precedes most drug development. The *α*-value of 0.05 remains almost universal in preclinical studies. The power (1 – *β*) continues to be lower than asserted because of the overestimation of effect sizes and consequent under-estimation of necessary sample sizes. Moreover, the prior probability of a hypothesis being true, (*γ*), may not be much greater than for a randomly selected hypothesis, given that many of the research findings purported to support the tested hypothesis may themselves be false discoveries.

### Second system flaw: the definitive target validation experiment is delayed to the end of drug development pipeline

The phase 3 randomised controlled trial (RCT) is often regarded simply as a test of the efficacy and safety of a new compound for a particular disease indication. However, when the compound evaluated is the first in its class, the RCT is also the first human test of the causal relevance of a previously untested drug target in a particular disease. This exposes the second major system flaw in the development of drugs with a novel mechanism of action: the most important target identification and validation experiment is the concluding not the initiating step. Risk therefore accumulates rather than diminishes as a drug development programme progresses towards the RCT, accounting for the high actual and opportunity cost of late-stage failure. A theoretical solution to this problem would be to obtain large-scale randomised human evidence on a target and disease state earlier in a drug development programme, without recourse to developing a medicinal compound to obtain the necessary evidence. Though this might seem unattainable at first glance, human genomics again provides a solution. Population genetic association studies can be viewed as ‘natural randomized trials’ without drugs ^62 63 64 65^. This is because germ line genetic variants such as single nucleotide polymorphisms (SNPs), which associate with differences in expression or activity of an encoded protein, assort at random according to Mendel’s Law, in an analogous way to drug treatment allocation in a randomised clinical trial.

In comparisons of genetic associations in populations with drug treatment effects in clinical trials, using a set of biomarkers and disease outcomes common to both study types, SNPs in a gene encoding a potential drug target have been observed to anticipate the mechanism-based effect of pharmacological action on the same protein. The approach is sometimes referred to as Mendelian randomisation for drug target validation (see **Appendix 1**, Ref 1), since it was inspired by, and represents a special case of the Mendelian randomisation paradigm, developed initially to help determine the causal relevance of environmental exposures or disease related biomarkers^66^. Mendelian randomisation for drug target validation is disease agnostic, though it may be unsuited to aspects of cancer drug development, where somatic rather than germ line mutations perturb the targets of interest, or to the development of anti-infective drugs, in cases where the therapeutic drug target is in the pathogen rather than the human host.

Importantly, genotyping arrays containing many thousands of SNPs across the genome, including those in genes encoding potential drug targets, provide the opportunity to interrogate systematically the influence of genetically mediated target perturbation on hundreds (eventually perhaps thousands) of biomarkers and disease outcomes in parallel, in a manner analogous to high-throughput compound screening (HTS) against a target. In this way, a genome-wide extension of the Mendelian randomisation paradigm could be used for drug target identification.

### Genomic studies for disease-specific target identification

There are sound reasons for thinking that genomic studies to specify drug targets for a human disease is likely to be a more reliable approach than the standard hypothesis-driven, non-genomic preclinical research in cells, tissues and animal models described previously. This is because:

a. The evidence obtained in GWAS comes from intact humans, the species of interest, not isolated cells, tissues studied *ex vivo*, or animal models
b. GWAS are some of the most statistically robust study designs in any field of biomedicine by virtue of their low false discovery rates, large sample sizes and the routine replication of positive findings
c. Genetic associations are protected from certain biases that affect other human observational study designs by virtue of the natural randomisation of genetic variants, which mimics treatment allocation in an RCT.
d. With appropriate coverage of the set of genes encoding human drug targets, and an adequate sample size, GWAS can be conducted for most (if not all) human drug targets simultaneously

Indeed, the same arguments apply to studies in which whole exome or whole genome sequencing (rather than genotyping) is used as the primary means of acquiring information on naturally occurring genetic variation and its association with disease.

Evidence is already emerging that such genetic association studies can help systematically match the correct drug targets to the correct disease. This comes partly from the like-with-like comparisons of the effects of licensed drugs on biomarkers and disease outcomes in clinical trials with the association of variants in the gene encoding corresponding drug target in population studies, examples of which, now span several diseases (**Appendix 1**). It also comes from the apparently sporadic ‘rediscovery’ by GWAS of drug targets already exploited for the treatment of the corresponding disease, as well as rediscoveries of the known mechanism-based adverse effects of several drug classes. We provide examples of this in **Table 4** and a linked paper^67^.

**Table 4.**
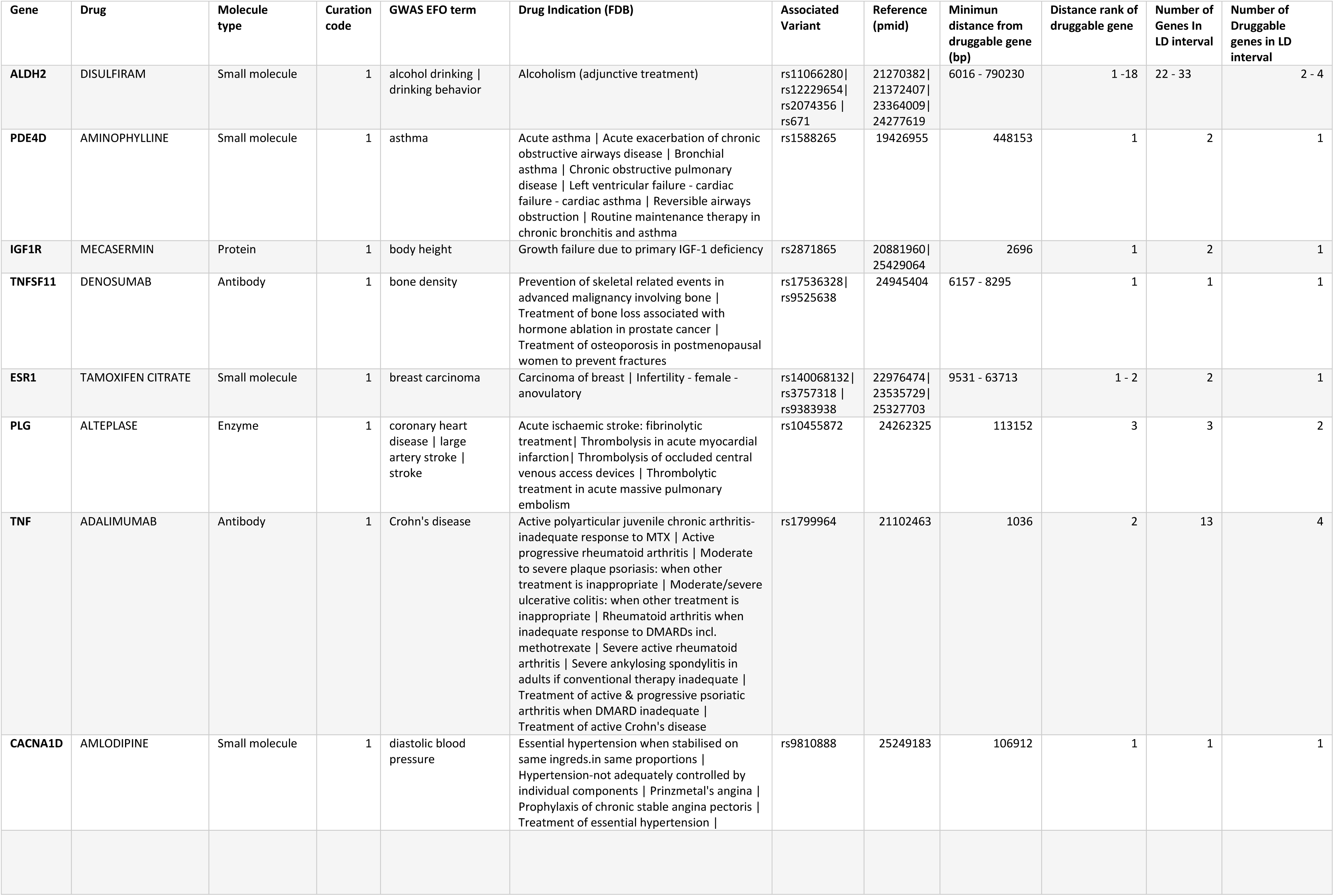
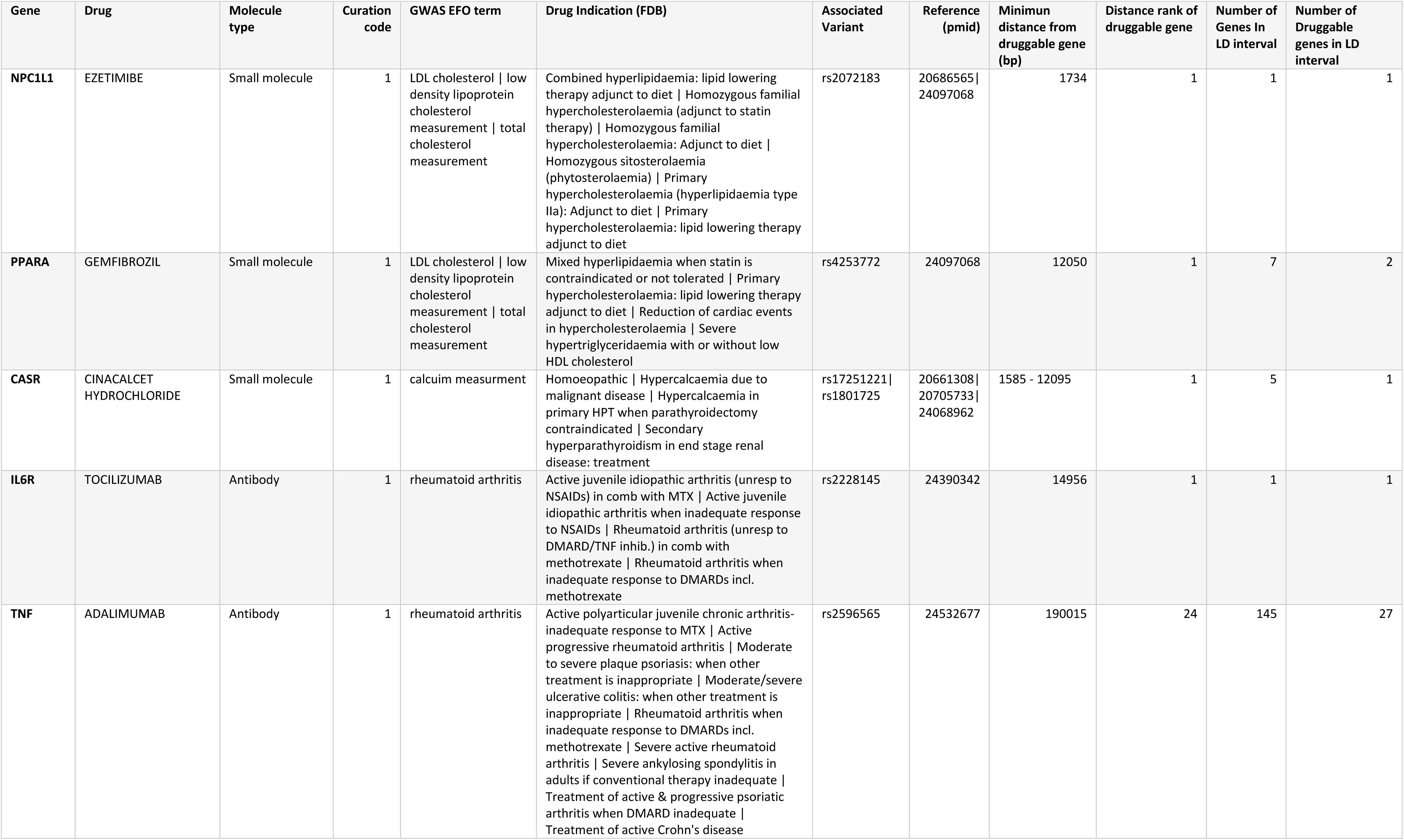
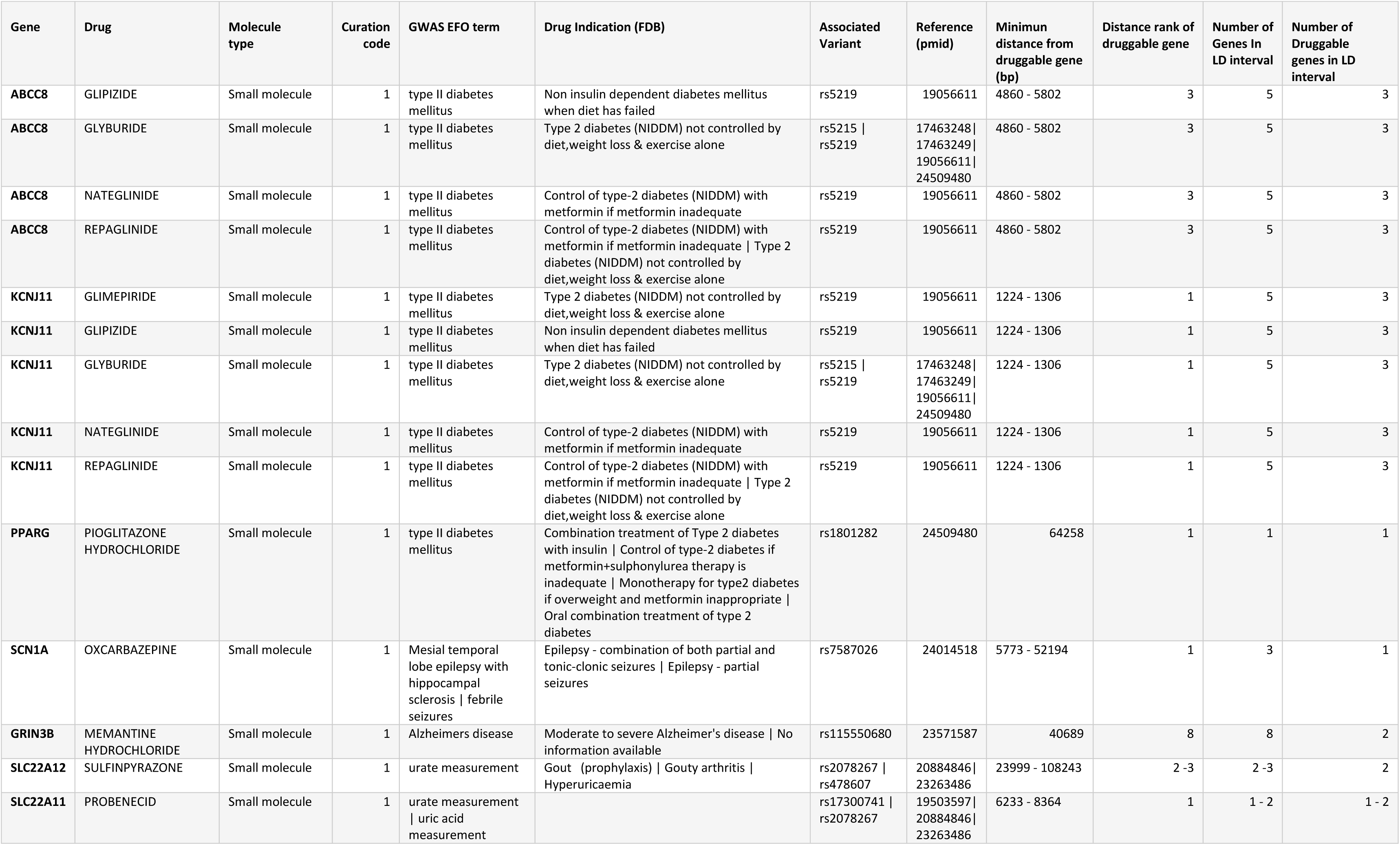
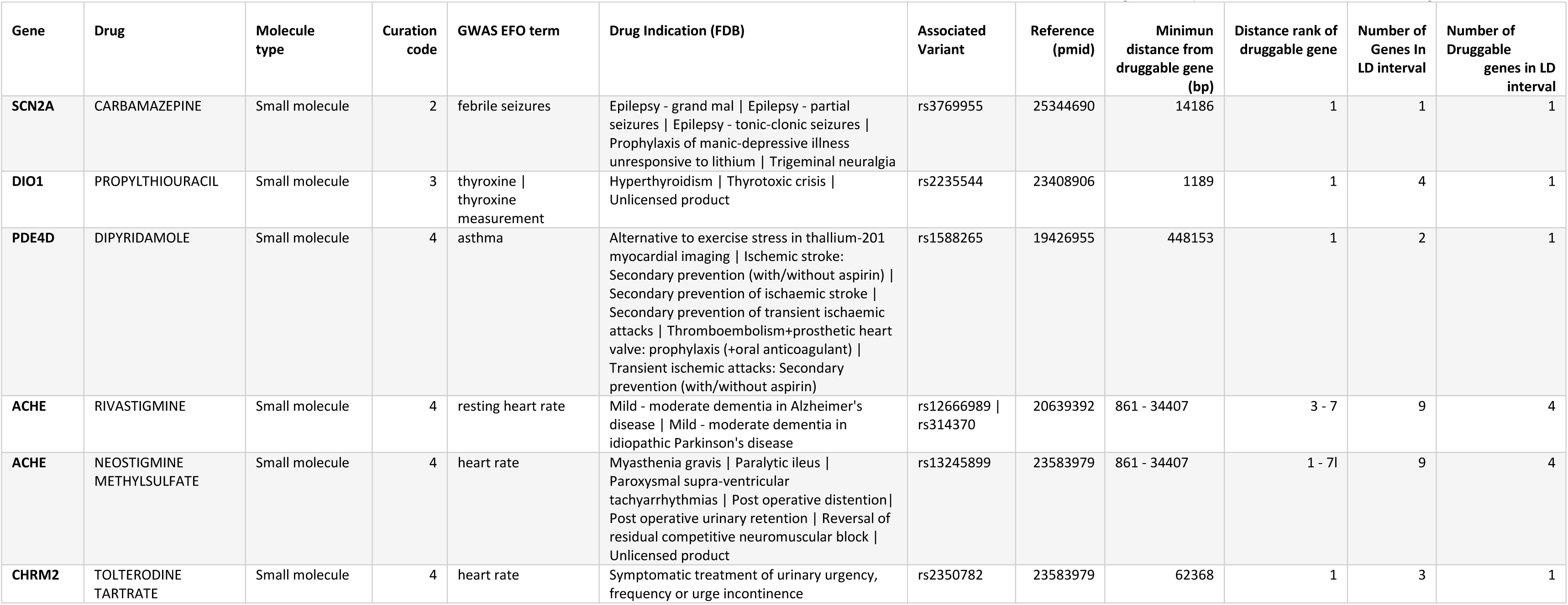
Illustrative examples of mapping SNPs curated in the GWAS catalogue to genomic linkage dis-equilibrium (LD) intervals containing targets of licensed and clinically used drugs (adapted with modification from Finan et al. http://biorxiv.org/content/early/2016/07/26/066027). The gene encoding the drug target is listed using Human Genome Nomenclature Catalogue designation. Drug names and indications are from First Data bank. GWAS SNPs are listed according to Refseq number and physical distances are in base pairs (bp). Curation code refers to the correspondence between the treatment indication and GWAS disease or trait association (see Text). Examples are shown of treatment indication rediscoveries which refer to a drug target indication-genetic association match (Curation code 1= precise match, code 2=disease area match). For many of these the drug target gene is the sole occupant of the LD interval defined by the GWAS SNP. Examples come from a variety of disease areas and, for some diseases (e.g. type 2 diabetes and rheumatoid arthritis), multiple target rediscoveries are noted. Examples of rediscoveries of mechanism of action (curation code 3) and mechanism-based side effects are also seen (curation code 4)

But are such rediscoveries fortunate coincidences or predictable occurrences that can be harnessed for the purposes of drug development?

To address this question, we formalise some further assumptions. Again, we discuss their validity in a later section.

**Assumption 4:** Drug treatments for human disease target proteins encoded in the germ line^a^
**Assumption 5:** DNA sequence variants in and around a gene encoding a drug target, that alter expression or activity of the encoded protein (*cis*-acting variants) are ubiquitous in the genome
**Assumption 6:** The association of *cis*-acting variants with biomarkers and disease end-points in a population genetic study accurately predict the effects of pharmacological modification of the encoded target in a clinical trial
**Assumption 7:** Genotyping arrays used in GWAS provide comprehensive, appropriately powered coverage of the genome, and associations discovered at any one gene are independent of those detected at any other

Among those diseases that have at least one licensed drug treatment, the total number of targets will vary. For example, nine drug classes (corresponding to nine different drug targets) contain compounds currently licensed for the treatment of type 2 diabetes (insulin, metformin, sulphonylureas, meglitinides, glitazones, DPP IV inhibitors, GLP-1 receptor agonists, SGLT-2 inhibitors and acarbose), but only two therapeutic classes (cholinesterase inhibitors and NMDA-receptor antagonists) contain compounds licensed for treatment of dementia. We can safely assume, from the efficacy of these drugs, that their targets (along with others, yet to be identified) play a causal role in those diseases.

Consider a hypothetical disease (*d*_1_) for which there are *n*_1_ independent genes encoding targets of drugs that have already been licensed on the basis of proven efficacy in the condition. We denote these as genes *g*_1_, *g*_2_…*g_n_*. Let us assume that a GWAS in disease *d*_1_ utilises a genotyping array with adequate coverage of all *n*_1_ licensed drug target genes, and that there is a probability ((1 – *β*_1_), (1 – *β*_2_)…(1-*β*_*n*_1__) of detecting the genetic association at each of these loci. Thus (1 – *β_i_*) is the power (or the detection rate) for a real effect of gene *g_i_* in disease *d*_1_.

We consider testing for a genetic association at the locus encoding each drug target in each hypothetical GWAS of *d*_1_ to be an independent trial (**Assumption 7**), where success equates to detection of an association at the locus and failure to overlooking the association. Consider a situation in which there are 3 licensed drug targets in disease *d*_1_ that are available for rediscovery, and that power to detect true associations is the same at all 3 target loci (i.e. (1 – *β*_1_) = (1 – *β*_2_) = (1 – *β*_3_) = (1 – *β*). The probability of missing such a target, is the false negative rate *β* A GWAS in *d*_1_ might detect 0,1,2 or all 3 of the known drug targets, and the probability that each of these situations occurs is given by the binomial distribution:

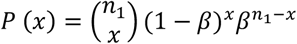

*P* (*x*) = the probability of detecting × licensed drug targets
*n*_1_ = the number of licensed drug targets in disease *d*_1_
*n*_1_ – *x* = the number of undetected licensed drug targets
*β* = Type II (false negative) error rate at each genetic locus

If *n*_1_ = 3, and *β* = 0.2, the probability (*P*) that a GWAS in disease *d*_1_:

- Detects none of the three licensed drug target genes, *P*(*x* = 0) = *β*^3^ = 0.008
- Detects only one of the three licensed drug target genes but misses the remaining two, *P*(*x* = 1) = 3*β*^2^ (1 –*β*) = 0.096
- Detects only two of the three licensed drug target genes but misses the other, *P*(*x* = 2) = 3*β* (1 – *β*) ^2^ = 0.384
- Detects all three licensed drug target genes, *P*(*x* = 3) = (1 – *β*)^3^ = 0.512
- Detects at least one of the three licensed drug target genes, *P*(*x* > 0) =1 –*β*^3^ = 1 – 0.008 = 0.992

In general, the expected (average) number of licensed drug target rediscoveries (*E_d_*) detected in a GWAS of a disease *d* with *n_d_* licensed drug targets will be:

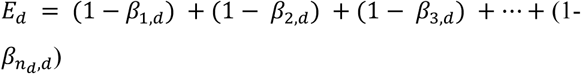

If power at all loci is (1 – *β*):

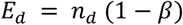

The variance (*V_d_*) is given by:

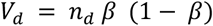

For example, for a GWAS conducted in disease *d* with (1 – *β*) = 0.8 at all three loci encoding the targets of licensed drugs:

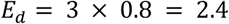

The variance (*V_d_*) = 3 × 0.8 × 0.2 = 0.48

The standard deviation (*SD_d_*) = 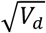 = 0.7

In the worked example, we would therefore expect 2.4 (*SD* = 0.7) of the 3 possible licensed drug targets to be rediscovered, on average.

Suppose we do one GWAS for each of *K* different diseases (*d*_1_, *d*_2_…*d_K_*) where, for each disease, the number of licensed targets available for rediscovery is (*n*_1_, *n*_2_,…*n_K_*). If we assume that the power to detect an association at gene *i* encoding the target of licensed drug is the same for all drug targets in *all* GWAS *j*, regardless of disease (i.e. (1 – *β*_*i*,*j*_) = (1 – *β*) for all *i* and *j*), then the expected number of true drug target-indication rediscoveries (*E_T_*) across the *K* GWAS would be the sum of the expected rediscoveries in each GWAS. Therefore:

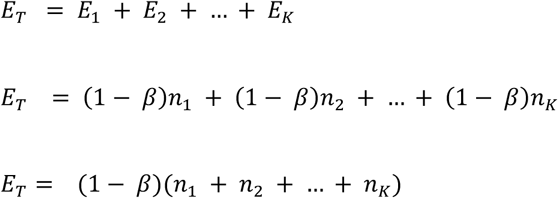

Thus,

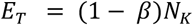

Where *N_K_* = (*n*_1_ + *n*_2_ +…+ *n_K_*)= the total number of licensed drug targets for *K* diseases

Dividing and multiplying the above equation by *K*, we obtain:

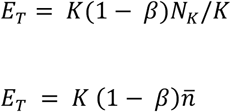

Where;

*n* = *N_K_*/*K*= the average number of targets of licensed drugs per disease

The standard deviation (*SD_T_*) is given by:

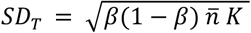

Suppose a GWAS was done for each of 200 different diseases, each with power (1 – *β*) = 0.8 to detect each true licensed target, and *n* = 3 (i.e. an average of 3 targets per disease and *N_K_* = *nK* = 600 potentially re-discoverable target-disease combinations in total).

The total number of licensed drug target rediscoveries from the combined dataset would be expected to be:

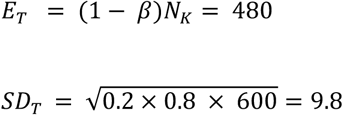

Values of *E_T_* for a range of plausible values of *β* and *n*, given *K* = 200 are provided in **Table S1**

It seems reasonable to ask if the number of licensed drug target rediscoveries already made by GWAS is close to that expected from these arguments. However, the answer is not straightforward. It requires enumerating the number of GWAS that have already been done for conditions that correspond to either a treatment indication or a mechanism based adverse effect for at least one licensed drug target, and counting the total number of licensed drug targets represented across all these conditions (since some diseases may be connected with multiple licensed drug targets). These efforts are hampered by different disease terminologies being used when cataloguing GWAS, drug indications and adverse effects. There is also a requirement to make strong assumptions about the average power of eligible GWAS to detect a true association at a gene encoding a licensed drug target.

However, the question can be inverted: given the observed number of rediscoveries, what was the average power of GWAS to rediscover loci encoding licensed drug targets for the same indication or through a known mechanism-based adverse effect? We previously reported that GWAS to 2015 had encompassed 315 unique MeSH disease terms and led to the ‘rediscovery’ of 74 of the 670 or so known licensed drug targets, either through treatment indication, or mechanism-based adverse effect associations^67^.

To estimate average power, we use:

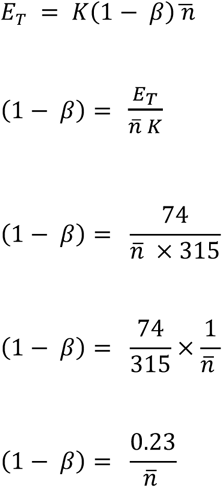

If *n* = 1,(1 – *β*) = 0.23

If *n* < 1,(1 – *β*) > 0.23 (as would be the case if some GWAS concerned diseases with no licensed drug target available for rediscovery)

If *n* > 1,(1 – *β*) < 0.23

Despite the modest estimated average power, the discovery by GWAS of around 70 of the 600 or so known licensed targets (**see Box 6**), suggests the approach shows promise as a means of identifying target-disease indication pairings more systematically in the future, particularly if power were to be enhanced. We return to this point in a later section.

### Estimating the yield of all druggable targets by GWAS

In the previous section, we discussed the rediscovery of known licensed drug targets by GWAS. In this section, we discuss the potential for GWAS to specify new drug targets for common diseases prospectively.

To estimate the total number of drug target - disease indication discoveries that might be possible in adequately powered GWAS with comprehensive coverage of the genome, we return to the concept of a sample space demarcated by 20,000 human genes and 10,000 common diseases.

Since only a portion of the genome encodes proteins that are readily accessible to small molecule drugs, monoclonal antibodies or peptides that currently comprise the major chemical categories of medicines, we now define the following:

{T} = the set of genes encoding druggable targets (the druggable genome – See **Box 6** for definition)

*N_T_* = Total number of genes encoding druggable targets = 4000 (**see Box 6**)

#### Box 6. The druggable genome

In 2002, at a time when the human genome was thought to contain ~30,000 protein coding genes, Hopkins and Groom estimated that 120 targets had already been exploited by licensed drugs but that ~3000 genes in total encoded proteins potentially accessible to small molecule agents, coining the term ‘the druggable genome ^68^. Subsequent estimates of the druggable genome have included between 2000 and 10,000 genes depending on the data set used and assumptions made^69 70^. Our recent work in developing a genotyping array with marker coverage of genes encoding actual or potential drug targets, led to a revised estimate that approximately 4000 human genes (or about one fifth of the protein-coding genome; see **Box 4**) encode druggable proteins ^67^. We use this estimate in the calculations that follow. Notably more than half of the known small molecule drug targets belong to four key gene families: class I G-protein coupled receptors (GPCRs), nuclear receptors, and ligand- or voltage gated ion channels, while targets for monoclonal antibodies or peptide therapeutics are cell membrane-bound or secreted and circulating proteins^71^. Rask-Anderson et al^72^ note around 555 targets are already exploited by currently licensed drugs (around 12% of the druggable genome) with a further 475 unique targets being the subject of investigation in clinical trials. More recently, Santos et al. estimated that FDA approved drugs for human diseases target 667 proteins encoded by the human genome^71^. Therefore, in combination, about a quarter of the druggable genome (one-twentieth of the whole genome), has already been drugged by licensed therapies or those in clinical phase development. Note again that antimicrobial treatments that interfere with targets in a pathogen rather than human host, and cancer treatment targets encoded by an abnormal cancer cell genome, distinct from the germ line, are excluded from these estimates.

With *N_G_* = 20,000, and *C* = 100, we showed the probability *P_C_* of selecting a causal protein-disease pairing from the sample space at random (**Equation 3**) is given by:

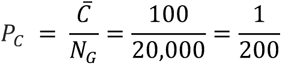

The probability (*P_T_*) of selecting a druggable gene (protein)-disease pairing at random from the sample space is independent of the number of diseases, and is given by:

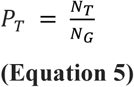

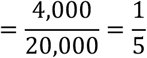

To estimate the probability *P_CT_* of selecting a disease-causing, druggable protein-disease pairing at random from the sample space we introduce a further assumption.

**Assumption 8:** The probability that a protein affects disease pathogenesis and the probability the protein can be targeted by a drug is independent.

Therefore,

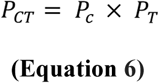

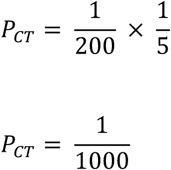

(**see Figure 2**).

Corresponding probabilities and counts for scenarios in which *C* = 100, and *C* = 1000 are shown in **Figure S1 and S2 and Table S2**. Note that these probabilities are independent of *N_D_*, the number of common diseases.

Following the arguments presented previously (see **Equation 4**), *P_CT_* can also be interpreted as *γ_CT_*, the true proportion of causal, druggable gene-disease pairs from the sample set of all gene-disease pairings.

These probabilities are of general interest, but the probability of more direct interest is that of identifying a druggable, disease-causing gene having already specified the disease of therapeutic interest.

Since we assume the probability of a protein influencing the pathogenesis of one disease is independent of the probability that it influences any other (**Assumption 3**), the values for *P_C_*, *P_T_* and *P_CT_* are the same for each individual disease as they are for the complete sample set.

We can therefore write, for any given disease, with *C* causal genes:

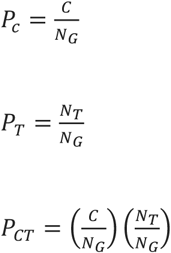

These estimates can now be used to re-assort all genes in the genome for a given disease from a therapeutic perspective (**Figure 3**).

**Figure 3.**
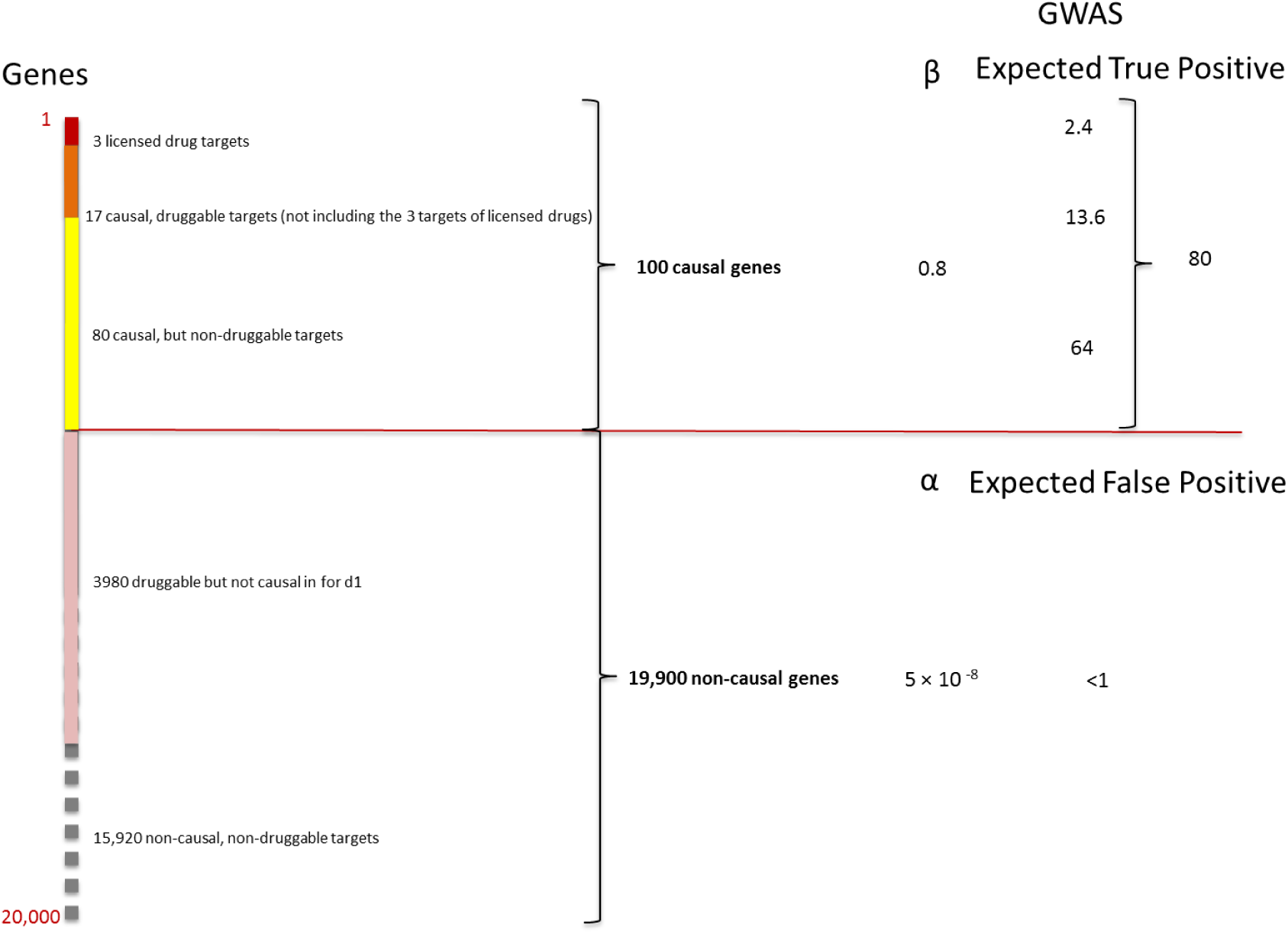
Re-assorted ‘therapeutic genome’ of a hypothetical disease (*d*_1_). The 20,000 protein coding genes are organised into 100 causal and 19,900 non-causal genes. Causal genes are further subdivided into 20 that are also druggable and 80 that are not. Of the 20 causal, druggable genes, 3 are the targets of licensed drugs for the treatment of *d*_1_. Of the non-causal genes, 3980 are druggable but not causal for *d*_1_. The right hand panel indicates the expected number of true and false positive genes (including druggable genes) expected in a GWAS of *d*_1_ undertaken with a sample size that provides power, 1 – *β* = 0.8 and type 1 error rate of *α* = 5 × 10^−8^ at all loci.

For example, in the hypothetical disease (*d*_1_), where *C* = 100, the expected number of causal and druggable genes is given by:

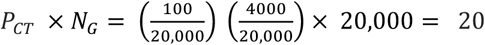

Eighty of the 100 causal genes would therefore be categorized as non-druggable. Of the remaining 19,900 non-causal genes, one fifth (3980) would be expected to be druggable but not causal in disease *d*_1_ (though of course they might be causal and of therapeutic interest in a different disease). The remaining 15,920 genes would be classified as neither causal for *d*_1_, nor druggable.

Assuming a GWAS in *d*_1_ interrogates each of the causal protein-coding genes with power (1 – *β*) = 0.8, the expected number of causal, druggable targets (*E*_*CT*,*d*1_ identified by such a GWAS is given by:

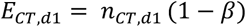

(where *n*_*CT*,*d*1_ is the true number of causal, druggable targets in *d*_1_)

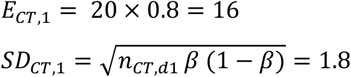

The probability of a GWAS detecting *x* = 0,1,2,3,4,…all 20 of the available causal, druggable targets is again given by the binomial distribution:

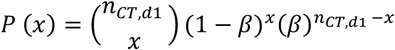

where:

*P* (*x*) is the probability of detecting *x* causal, druggable targets *n*_*CT*,*d*1_ is the number of causal, draggable targets in disease *d*_1_ (20 in this example)

*n*_*CT*,*d*1_ – *x* is the number of causal, druggable targets not detected in the GWAS

(1 – *β*) is the power of the GWAS to detect a true association at a genetic locus (set at 0.8 in this analysis and assumed to be homogeneous for all loci)

In summary, with *C* = 100, *P_C_* = 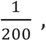 *P_T_* = 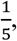 i.e. 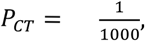 a GWAS with power 1 – *β* = 0.8 at all loci would be expected to discover 16 (*SD* 1.8) of the 20 available, causal, druggable targets, on average. Moreover, it would be extremely unlikely that a GWAS with (1 – *β* = 0.8) at all loci, would discover fewer than 10 druggable targets.

The exceedingly stringent type I error rate (*α*) incorporated in such studies (e.g. 5 × 10^−8^) also makes the probability of even one false target discovery being present among the declared associations very low indeed (**Figure 3**). These calculations suggest that adequately powered GWAS (designed with appropriate consideration of the distribution of genetic effect sizes, sample size and comprehensive coverage of sequence variation in protein coding genes) should provide a highly accurate and reliable way of specifying drug targets for human diseases, addressing the high *FDR* problem that underpins inefficiency in drug development.

## Part 2: Probability of drug development success

> ‘***The Industry must rethink its process culture. Success in the pharmaceutical industry depends on the random occurrence of a few ‘black swan’ products.’***
>
> — -Bernard Munos. Lessons from 60 years of pharmaceutical innovation. *Nature Rev. Drug Discov.* 2009 8, 959–968

If our assessment is accurate, the use of genomic information to support drug target identification should offer an opportunity to improve drug development success rates by bringing statistically robust, large-scale, randomised evidence from humans much earlier (even to the very start) of a drug development programme. But is it possible to quantify what the improvement in drug development efficiency might be?

Recent analyses have considered the influence of genomic evidence on drug development success rates but mainly from a retrospective viewpoint based on observed frequencies: e.g. ‘what are the observed rates of progression from one developmental phase to the next’ and, ‘to what extent have successful vs. unsuccessful drug development programmes had prior genetic support for the target?’ ^27 73^.

Instead, we consider:

a. The *a priori* probability of accurate target identification comparing orthodox (non-genomic) with genomic approaches.
b. The number of orthodox (non-genomic) drug development programmes that need to be pursued in parallel to ensure 90% probability of at least one licensing success
c. The probability of repurposing success
d. Preclinical target identification as a ‘predictive test’ for drug development success, comparing orthodox (non-genomic) with genomic approaches

We then go on to use observed rates of preclinical and clinical development success to estimate the proportion of true target-disease relationships that are studied in contemporary drug development. Finally, we gauge the impact of the target selection step on ultimate success rate, which is necessary in orthodox (non-genomic) but not genomic preclinical development

### *A priori* probability of accurate target identification

Around 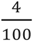 preclinical drug development programmes yield licensed drugs^1 2^. However, this estimate is based on the success rates of compounds rather than targets. The success in early development of a first-in-class molecule for a given disease indication is often followed by a flurry of development programmes, distributed across several companies, based on the same target and disease indication. The consequence is that multiple drugs may emerge, all in the same class (e.g. there are 7 different HMG coA reductase inhibitors (statins) licensed for lowering LDL-cholesterol for coronary heart disease prevention, and >12 different angiotensin converting enzyme inhibitors for the treatment of hypertension, heart failure and related conditions. Using the ChEMBL database, we estimate a median of 2 (mean of 4) licensed drugs per efficacy target (**Figure 4**). Therefore, the overall developmental success rate for targets could be around half that of compounds i.e.

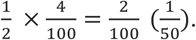

**Figure 4.**
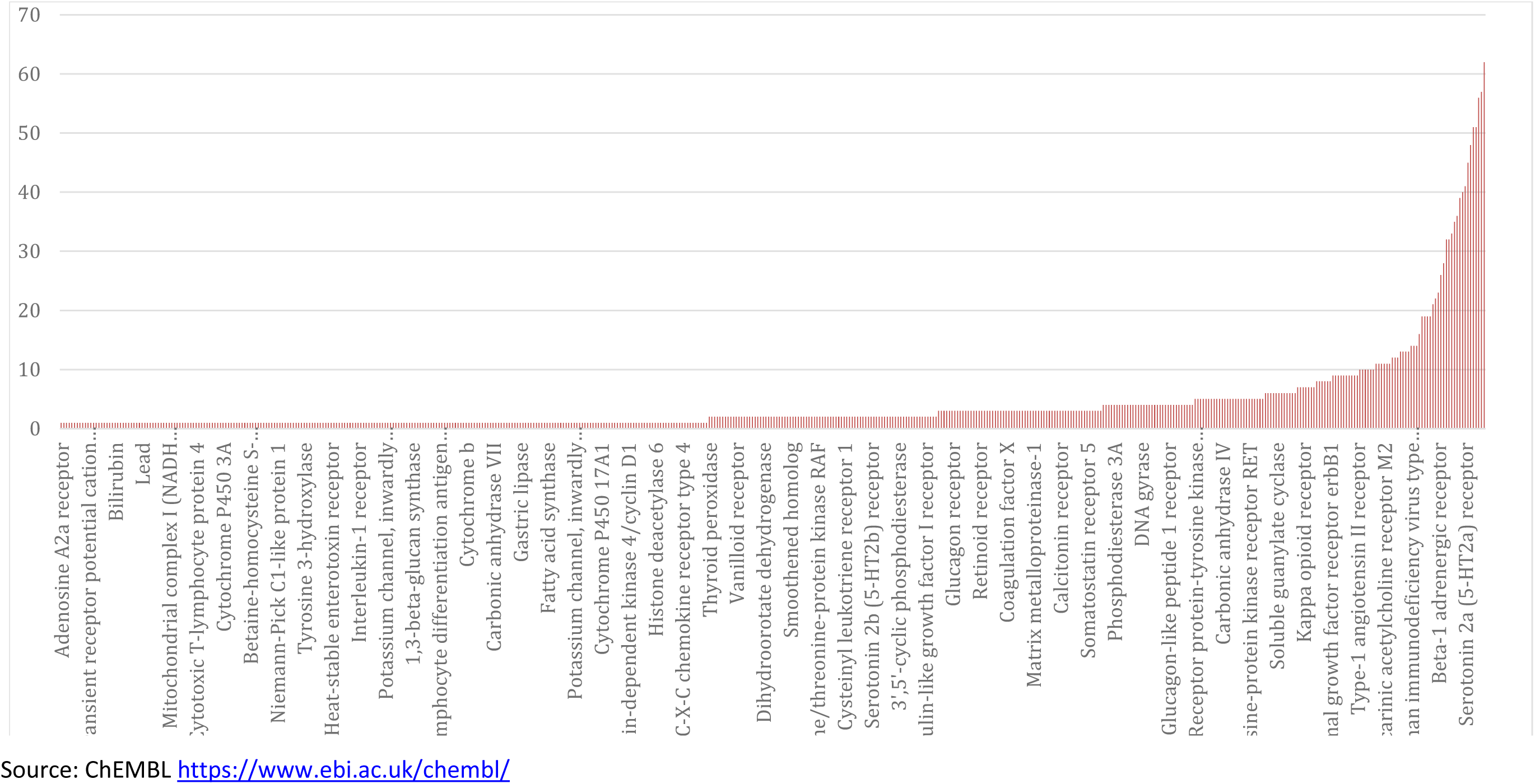
Distribution of number of licensed drug compounds per target

Drug development success depends on correctly identifying a causal, druggable target-disease indication pairing, and then demonstrating the validity of the target in preclinical studies, and the efficacy of target modification in clinical trials.

We showed previously (see **Equation 6**) that the a prior probability (*P_CT_*) of selecting a disease-causing, druggable protein-disease pairing at random is:

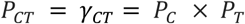

From **Equations 3 and 5**;

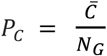 in the general case, or 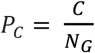 in the case of a specific disease, where *C* = average number of causal genes per disease, and *C* =the number of causal genes in the disease of interest.

Thus, for a given disease:

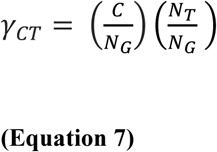

Based on **Equation 7**, *γ_CT_* could be increased, in theory, by increasing *C*, increasing *N_T_*, or by reducing *N_G_*.

**Table S2 and S3** illustrate the influence of different estimates of *C* on the probability on *P_C_* = *γ_C_* and *P_CT_* = *γ_CT_*.

*C*, however, is not amenable to manipulation, being largely determined by evolutionary forces;
*N_G_*, is also fixed;
*N_T_*, however, could be increased by developing technologies that allow a broader range of gene products to be targeted therapeutically.

It can be argued that the development of therapeutic monoclonal antibodies has already increased *N_T_* by permitting targeting of proteins that were not previously amenable to a small molecule therapeutic strategy^74^. (The development of therapeutic antisense RNA and related technologies is likely to further extend future therapies into the RNA target space).

However, there are also ways of reducing the number of genes under consideration in a given disease, so as to increase *γ_CT_*.

Consider focusing solely on the druggable genome in a given disease. We can then write:

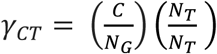

Therefore;

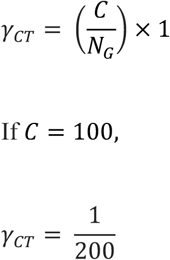

Thus, among the set of druggable genes, all causal genes are automatically both causal and druggable. Therefore, if *C* = 100, the simple expedient of focusing target identification in a specific disease on the 4000 or so druggable genes, rather than the genome as a whole, increases *γ_CT_* by a factor of five from 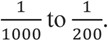

Alternatively, we could remove genes from consideration that we perceive to have a low probability of playing a causal role in the disease of interest, instead focusing on a subset of the genome *N*_*C*′_, where *N*_*C*′_ = the set of likely to be causal genes in the disease of interest.

We could then write:

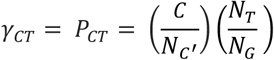

If it were possible to enrich the sample space by progressively eliminating all non-causal while retaining all causal genes, then:

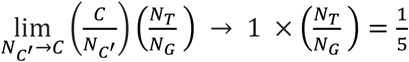

Thus, in the limiting case, among an exclusively causal set of genes, the probability of being causal and druggable is simply the probability of being druggable (**see Box 6 and Assumption 8**).

Eliminating non-causal while retaining causal genes is the crux of the target identification problem. For reasons we outlined previously, an adequately powered GWAS in a disease of interest, with a stringent *α* has the capability to exclude the non-causal while identifying the set of causal genes for any disease, of which 1/5^th^ on average 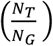 is
expected to be druggable under **Assumption 8**.

In summary, the probability of selecting a causal, druggable target for a disease of interest based on a random pick from the whole genome is 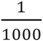 (assuming *C* = 100), but 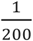 based on a random pick from the druggable genome. We note that these probabilities from a random pick are not vastly different to the *observed* rates of drug development success: 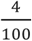 for compounds (perhaps closer to 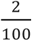 for novel targets). In a later section, we show that these estimates are also similar in order to values for *γ_CT_* (the proportion of causal and druggable target-disease pairs available for discovery) calculated *a posteriori* from reported preclinical and clinical development success rates ^2^.

Taken together, the calculations suggest that the current, mainly non-genomic preclinical approach to target identification only weakly enriches the sample space for causal target-disease pairings that are then taken forward into clinical development.

### Number of parallel development programmes required, to ensure 90% probability of at least one licensing success

A common industry strategy to address low developmental success rates has been to pursue multiple drug development programmes in parallel, recognizing that the majority will fail, but that even a single success could ensure profitability because of revenues generated through the patent system. For example, 1120 unique pipeline drug programmes for Alzheimer’s disease were initiated across the industry in the period 1995 – 2014^75^. But, with the estimated current developmental success rate of around 2% for targets, on average, how many programmes would need to be pursued in parallel to have a 90% chance of at least one success? This can be calculated as follows. Let:

*P_s_* = within – programme success rate.

Assuming all programmes are independent, the probability of all *N* programmes failing is:

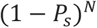

A 90% probability of at least 1 success equates to a 10% probability of no success in any programme (i.e. a 10% probability of all programmes failing)

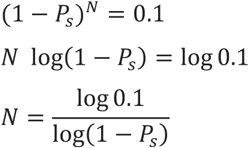

Let us assume:

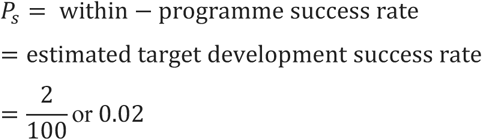

Therefore,

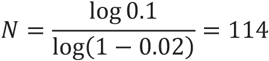

Thus, when *P_s_* = 0.02, industry needs to pursue 114 independent programmes in parallel, on average, to have a 90% probability of at least one developmental success; 34 programmes would need to be pursued to have an 50% (evens) chance of at least one success. Values of *N* for a range of hypothetical values of *P_s_* are shown in **Table S4**.

### Probability of repurposing success

Another approach to address poor drug development success rates is to try to identify new disease indications for drugs that failed to show efficacy for the original indication, but which have proved safe in man; or to expand indications for a drug already effective in one disease to another condition. However, repurposing or indication expansion relies on the assumption that different diseases share at least some common drug targets. How likely is this to be the case?

Again, this can be tackled from a probabilistic perspective using two of the previous simplifying assumptions:

**Assumption 3:** The probability of a protein influencing the pathogenesis of one disease is independent of the probability that it influences any other
**Assumption 8:** The probability that a gene (protein) affects disease pathogenesis and the probability that a gene encodes a druggable protein is independent

Repurposing or indication expansion can be considered from three perspectives:

- How many diseases are likely to be influenced by the perturbation of a single therapeutic target?
- How many diseases need to be considered for at least one pair to share a common therapeutic target, under the assumption of independence?
- How many diseases need to be studied to find at least one that will be affected by pharmacological perturbation of a particular target of interest?

*Diseases influenced by perturbation of a single protein:* We showed previously that the probability (*P_C_*) of identifying a causal gene-disease pairing *CD* from the sample space comprising all genes and diseases, *GD*, assuming *C* = 100 and *N_D_* = 10,000, is:

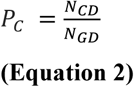

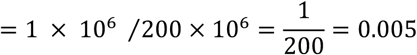

Under **Assumption 3**, the expected number diseases (*E_D_*) affected by any given gene is given by:

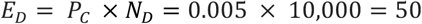

With standard deviation equal to:

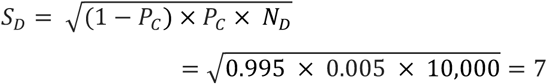

*E_D_* declines the fewer diseases under consideration, or if *C* < 100. (**Table S2**). Since the estimate of *E_D_* should be precisely the same for a gene encoding a druggable as a non-druggable target, under **Assumption 8**, it can be inferred that even the most specific of therapies is likely to influence a range of conditions; leading either to mechanism-based adverse effects, efficacy in more than one condition, or some combination of the two. In fact, under the assumptions above, perturbation of most therapeutic targets will affect between 36 and 64 diseases and only 1 in 1000 targets would affect 28 or fewer conditions.

*Shared therapeutic targets:* The second question is akin to a well-known statistical problem of how many people need to be assembled for at least one pair to share the same birthday.

Consider two diseases. Again, we assume *C* = 100. The first disease in the pair could have any 100 of the 20,000 genes in the genome as its causal set. The probability of the second disease sharing a number *x* of the 100 genes already involved in the first disease is given by the hypergeometric distribution:

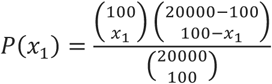

So, the probability that they do not share any gene is:

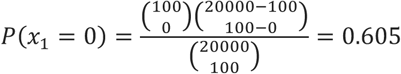

If we study a third disease, the probability of that disease not sharing any of the 200 genes involved in the previous two diseases would be:

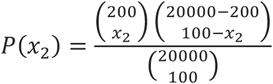

So, the probability of the third disease not sharing a single gene with the other two (*x*_2_= 0) is:

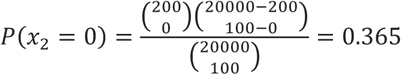

So the total probability of the three diseases not sharing any of the genes is:

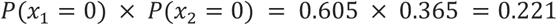

With four diseases, the probability of none of them sharing a gene is < 5%, and for eight diseases it is less than 1 in a million: it is almost certain that at least two diseases from this pool of eight, will share at least one common susceptibility gene.

*Number of diseases that need to be studied to identify at least one that is affected by perturbation of a given target:* The answer to the third question follows the same reasoning as that used previously to estimate the number of drug development programmes that need to be pursued in parallel to have at least a 90% or greater chance of at least one development success. With 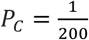 (i.e. focusing on the druggable genome), 459 diseases would need to be studied to have ≥ 90% chance of identifying at least one condition that is causally affected by perturbation of a particular target of interest. If *C* = 1000, the number of diseases that need to be studied is 45.

Despite these considerations, the ultimate challenge for repurposing remains the same as that for *de novo* drug development: knowing precisely which targets are important in which diseases and therefore which targets are shared among a set of diseases of interest. This, we believe, can only be tackled systematically by the genomic approach we have described in previous sections.

### Preclinical target identification as a ‘predictive test’ for drug development success

We next reduce drug development to a two-stage process: a preclinical component whose function is to predict target-disease pairings destined for clinical phase success (stage 1), and a clinical component (stage 2) whose function is to evaluate target-disease pairings brought forward from stage 1. Success in stage 2 is thus dependent on the predictive performance of stage 1.

Since clinical drug development failure, a consequence of incorrect target specification, currently accounts for around two in every three late-stage failures ^22 28^, we introduce one further simplifying assumption.

**Assumption 9:** Inaccurate target selection is the exclusive reason for clinical phase (stage 2) drug development failure.

Key variables in the following section are indexed by the lower-case suffix *pc* to denote preclinical and the lower case suffix *c* to denote clinical stage development.

Possible outcomes from pre-clinical and clinical phase development are summarized in the embedded tables below.

| Stage 1:<br>Preclinical<br>development<br>(*pc*) | True<br>relationship | No true<br>relationship | All |
| --- | --- | --- | --- |
| Declared<br>success | $TP_{pc} = \gamma_{pc}(1 - \beta_{pc})$ | $FP_{pc} = \alpha_{pc}(1 - \gamma_{pc})$ | $S_{pc}$ |
| Declared<br>failure | $FN_{pc} = \gamma_{pc}\beta_{pc}$ | $TN_{pc} = (1 - \alpha_{pc})(1 - \gamma_{pc})$ | $1 - S_{pc}$ |
| | $\gamma_{pc}$ | $1 - \gamma_{pc}$ | 1 |

| Stage 2:<br>Clinical<br>Development<br>(*c*) | True relationship | No true<br>relationship | All |
| --- | --- | --- | --- |
| Declared<br>success | $TP_c = \gamma_c(1 - \beta_c)$ | $FP_c = \alpha_c(1 - \gamma_c)$ | $S_c$ |
| Declared<br>failure | $FN_c = \gamma_c\beta_c$ | $TN_c = (1 - \alpha_c)(1 - \gamma_c)$ | $1 - S_c$ |
| | $\gamma_c = TDR_{pc}$ | $1 - \gamma_c$ | 1 |

$\gamma$ = proportion of true target-disease relationships $TP$ = true positive rate $FP$ = false positive rate $TN$ = true negative rate $FN$ = false negative rate $S$ = declared success rate $1 - S$ = declared failure rate

Declared preclinical successes (*S_pc_*) comprise both true and false positive findings. Therefore:

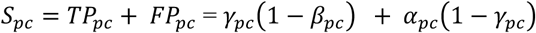

The proportion of true positive findings among reported preclinical successes equates to the preclinical true discovery rate (*TDR_pc_*), where:

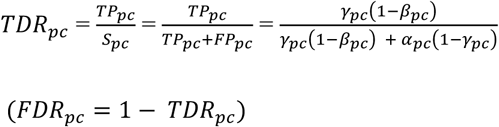

If a *clinical* phase drug development programme follows every declared *preclinical* success, the proportion of true target disease relationships in clinical phase development is equivalent to the *preclinical* true discovery rate, so we can write:

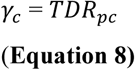

Similarly, for clinical phase (stage 2) development:

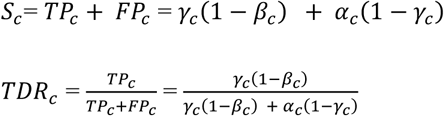

Since 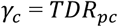(**Equation 8**)

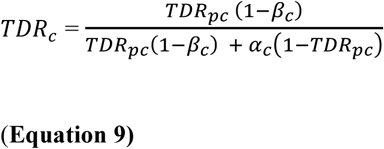

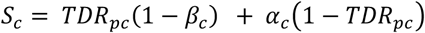

These equations underline the close mathematical relationship between preclinical and clinical discovery and success rates, which can be formalised as follows:

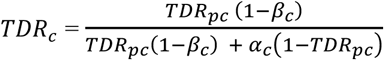

Dividing the numerator and denominator by *TDR_PC_* (1 – *β_c_*) and then rearranging:

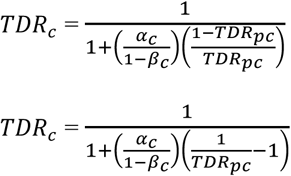

Since,

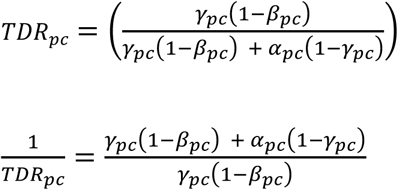

Consequently,

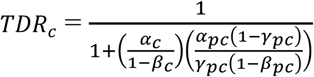

Rearranging,

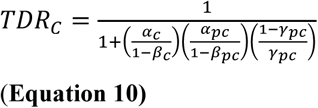

**Equation 10** illustrates that the clinical phase discovery rate can be resolved mathematically into terms that encompass clinical phase power and false positive rate (the term 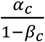), preclinical phase power and false positive rate (the term 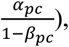 and the true relationships available for discovery (the term 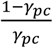). In this sense, **Equation 10** can be conceived as a mathematical summary of the probabilities and parameters determining drug development success.

Consider orthodox non-genomic preclinical (stage 1) drug development programmes with base case parameters defined by the sample space, *N_G_* × *N_D_* where:

*N_G_* = Total number of protein - coding genes = 20,000
*N_D_* = Total number of complex human diseases = 10,000
*C* = Average number of causal genes per disease = 100
*N_T_* = Total number of genes encoding druggable targets = 4000

From **Equation 7**, we can infer that;

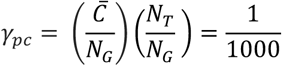

Setting *α_pc_* and *β_pc_* to 0.05 and 0.2 respectively, as is as standard for (non-genomic) preclinical experiments, and assuming it were somehow possible to evaluate every protein in every disease in such studies, then *TDR_pc_* = 0.016 and *FDR_pc_* = 0.984. *TDR_pc_* increases to 0.14 and the *FDR_pc_* falls to 0.86 if *C* = 1000 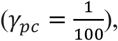 but the corresponding values are 0.002 and 0.998 if *C* = 10 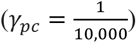 (**Table 2**).

In striking contrast, with the same sample space but a genomic approach to target identification, where (1 – *β*) = 0.8, *α* = 5 × 10 ^−8^ and all 20,000 targets encoded by the genome are, by definition, interrogated simultaneously, *TDR_pc_* = 0.999, and *FDR_pc_* = 0.001. This is a reversal of *TDR_pc_* and *FDR_pc_* values when compared to the orthodox (non-genomic) preclinical approach. The performance of genomic studies for target identification, based on these values of *α* and 1 – *β*, is little affected by 100-fold differences in *C* and *γ_pc_* (**Table 2)**.

As we showed previously, if sampling were restricted to the a sample space demarcated by the druggable genome, *N_T_* × *N_D_*, where;

*N_D_* = Total number of complex human diseases = 10,000
*N_T_* = Total number of genes encoding druggable targets = 4000
*C* = Average number of causal genes per disease = 100
*N_TD_* = Total number of possible druggable gene – disease pairs = 4,000 × 20,000 = 40×10^6^

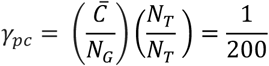

Focusing orthodox (non-genomic) preclinical studies on this restricted sample space (with conventional values for *α* and 1 – *β*) marginally increases the *TDR_pc_*(from 0.016 to 0.08) and reduces *FDR_pc_* but also only marginally (from 0.998 to 0.920). Applying the genomic approach in the same sample space, where (1 – *β*) = 0.8, and *α* = 5 × 10 ^−8^, and all 4,000 draggable targets encoded by the genome are interrogated simultaneously, the already high *TDR_pc_* increases to 0.9999, and the already low *FDR_pc_* would fall further to 0.0001. (**Table 2**).

It might be argued that *TDR_pc_* and *S_pc_* in conventional (non-genomic) preclinical pipelines could also be enhanced by simply setting a more stringent false positive rate in experiments involving cells, tissues and animal models. This is correct, but the change would have practical consequences. Very substantial increases in sample size would be required to maintain power. This might be perceived as being at odds with efforts to reduce the number of animals used in medical research, for example. However, in the long run, larger, more definitive large-scale animal experiments conducted early in the exploration of a hypothesis might actually make an important contribution to the goal of reducing the number of animals sacrificed, by minimizing wasted research. However, attending to the type 1 error rate issue alone fails to address the problem of the questionable validity of many animal models of human disease. It is also predicated on being able to evaluate every protein in every disease, a task we know to be beyond the capability of orthodox (non-genomic) preclinical studies based on cells, tissues and animal models. We return this issue in a later section.

Turning now to clinical (stage 2) development, *α_C_* and 1 – *β_c_* are typically set to 0.05 and 0.8 respectively, so it is also possible to examine the influence of variation in *γ_pc_*, *α_pc_* and *β_pc_* on preclinical (*S_pc_*), clinical (*S_C_*) and overall success (*S_o_* = *S_pc_* × *S_C_*), using **Equations 9** and **10**. The results are summarised in **Table 2**.

For orthodox (non-genomic) preclinical development, with sampling from the whole genome (where *C* = 100,1 – *β_pc_* = 0.8, *α_pc_* = 0.05, 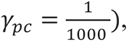 *S_pc_* = 0.05 (*TDR_pc_* = 0.016; *FDR_pc_* = 0.984) and *S_C_* = 0.06 (*TDR_C_* =0.2; *FDR_C_* = 0.8) giving an overall declared drug development success rate *S_o_* = *S_pc_* × *S_C_* = 0.003 (**Table 2**).

With the same parameters (*C* = 100, 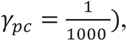 but with the genomic approach replacing orthodox non-genomic preclinical programmes, *S_pc_* = 0.0008 (*TDR_pc_* = 0.99994; *FDR_pc_* = 0.00006), *S_C_* = 0.79995 (*TDR_C_* = 0.999996; *FDR_C_* = 0.000004), and *S_o_* = 0.00064.

It may at first seem surprising that *S_pc_* (and *S_o_*) is actually lower for genomic than orthodox (non-genomic) stage 1 development, because of a higher stage 1 ‘failure’ rate. However, a ‘failure’ in a GWAS simply refers to a null association with the disease of interest of a specific gene (from all 20,000 evaluated), which is very different from the expensive failure of a lengthy orthodox preclinical development programme focusing on a single target at a time. The high ‘failure rate’ (i.e. high rate of null associations) in GWAS reflects the much more stringent *α_pc_* in this type of study design, which results in a much lower *FDR_pc_* and much higher *TDR_pc_.* Since *TDR_pc_*= *γ_C_*, the GWAS design ensures fewer false relationships are carried forward into clinical development when compared to the non-genomic approach. Consequently, *TDR_C_* is much increased with the genomic (compared to non-genomic) preclinical target identification. In summary, the calculations indicate that a genomic approach to preclinical target validation has the potential to reverse the probability of drug development success when compared to the established (non-genomic) approach.

### Estimating the proportion of true target-disease relationships currently studied based on observed development success rates

The preceding estimates of *γ_pc_* and the corresponding estimates of *S*, *FDR* and *TDR* are based on naïve pairings of genes (or proteins) and diseases (selection at random), using the sample spaces defined by common human diseases and either the whole genome or the druggable genome. But how closely do these estimates reflect current drug development?

Since observed values for *S_pc_* and *S_C_* have been reported ^2 28^, it should be possible to make *a posteriori* estimates of *γ_C_* and *γ_pc_* and other relevant metrics, and compare them to the *a priori* estimates based on a random pick of target-disease pairings in the sample space.

Both *γ_C_* and *γ_pc_* can be estimated from observed preclinical and clinical success rates as follows:

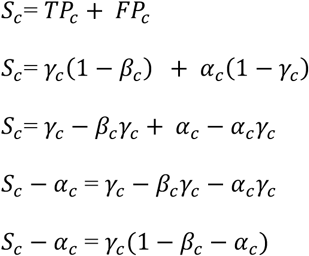

Therefore,

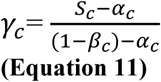

We previously established (**Equation 8**) that

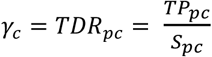

Since*TP_pc_* = *γ_pc_*(l – *β_pc_*

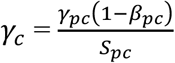

Rearranging, we have

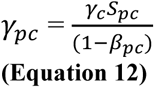

The reported clinical success rate ^2 28^, *S_c_* = 0.1

Assuming *α_c_* = 0.05, *β_c_* = 0.2 (commonly used false positive and negative rates for clinical trials) and using **Equation 11**:

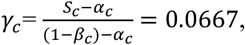

Since,

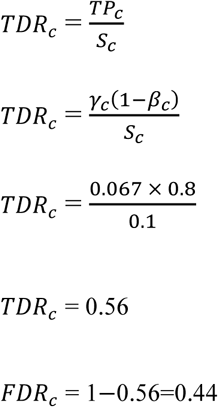

This calculation suggests that nearly one in two declared clinical trial successes may be a false discovery.

Since *γ_c_* = *TDR_pc_* and *TDR_pc_* = 1 – *FDR_pc_*

*TDR_pc_* = 0.0667

*FDR_pc_* = 1 – 0.0667 = 0.9333

These *a posteriori* estimates for *TDR_pc_* and *FDR_pc_* are of a similar order to the *a priori* estimates documented earlier.

Now,

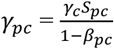

The reported preclinical success rate ^2^, *S_pc_* = 0.4

Using the value *γ_C_* = 0.0667, and setting power for preclinical studies at (l – *β_pc_*) = 0.8, we have:

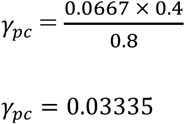

In estimating *α_pc_*, we use the following:

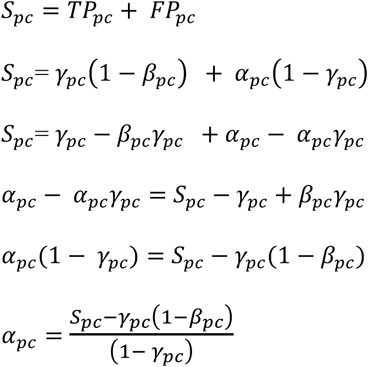

Note: the term *S_pc_ – *γ*_pc_*(1 – *β_pc_*) = *S_pc_* – *TP_pc_* = *FP_pc_*

Therefore 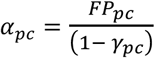 (see embedded table)

With *S_pc_* = 0.4; *γ_pc_* = 0.03335; and 1 – *β_pc_* = 0.8; *α_pc_* = 0.386

Values of *γ_pc_* and *α_pc_* for a range of values for 1 – *β_pc_* from 0.2 to 0.8, and a fixed value of *γ_C_* = 0.067, are illustrated in **Figure 5**. For values of 1 – *β_pc_*, in this range, values for *γ_pc_* lie in the range 0.033 to 0.133, representing 6.5-fold to 26.5-fold enrichment of true relationships over a random pick from a sample space demarcated by all diseases and the druggable genome 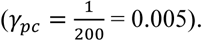 Although these enrichment rates for established preclinical drug development appear substantial, the very low values of *γ_pc_* mean that they are insufficient to prevent a large proportion of false target-disease relationships being pursued during clinical phase development, which accounts for the low rates of clinical success, and the possibility that a large proportion of declared clinical successes are actually false discoveries.

**Figure 5.**
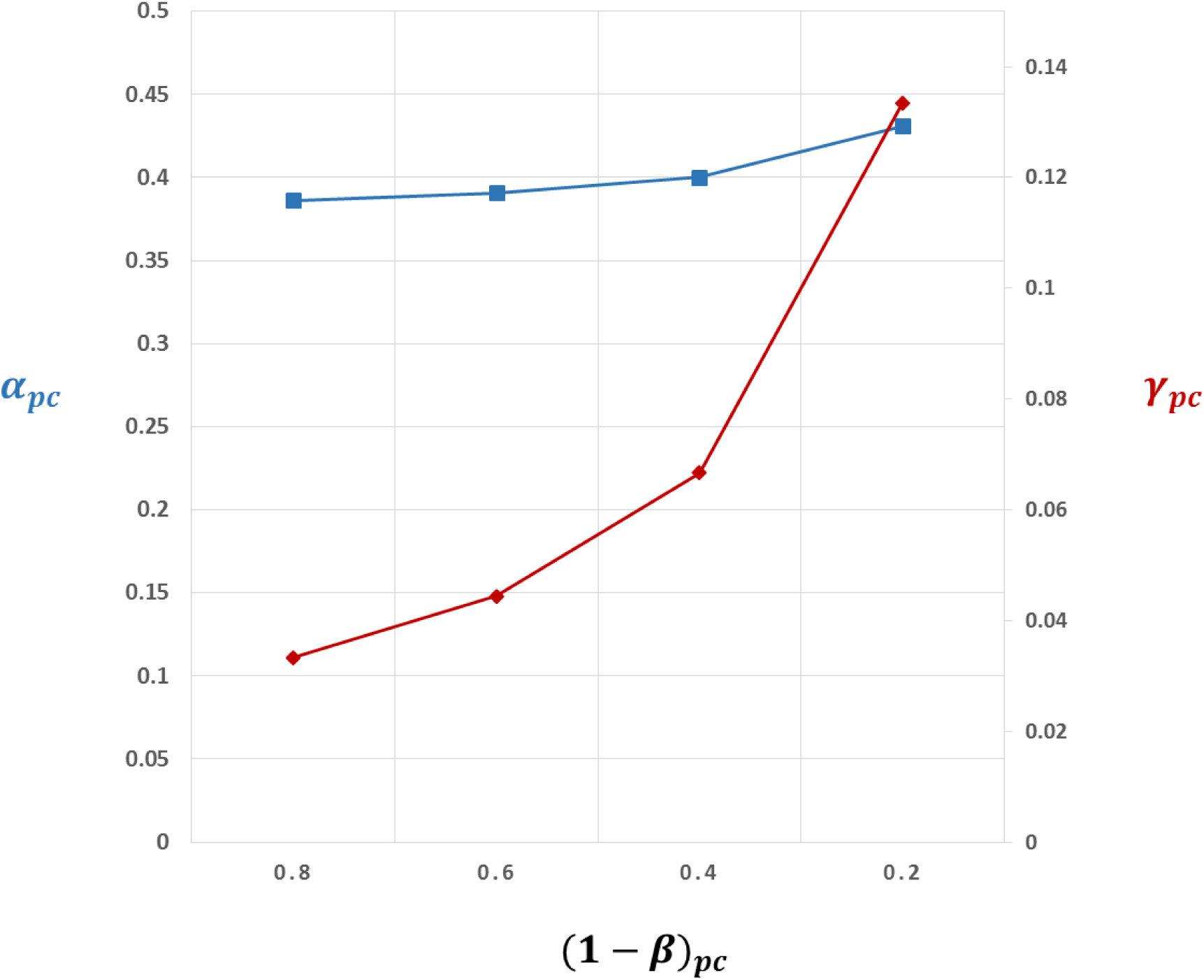
Back calculation of proportion of true target-disease relationships (*γ_pc_*) studied in preclinical development, inferred from observed rates of clinical success (*S_C_* = 0.1) and preclinical success (*S_pc_* = 0.4). Estimates of *γ_pc_* assume power in clinical phase development (1 – *β_C_*) = 0.8 and false positive rate in clinical development, *α_C_* = 0.05, so that the proportion of true target-disease relationships in clinical development, *γ_C_* = 0.0667. The graph shows estimates of *γ_pc_* (red line) for a range of values for power (1 – *β_pc_*) in preclinical development and corresponding estimates of the preclinical false positive rate, *α_pc_* (blue line). (See text for details).

### The impact of the target selection step in orthodox (nongenomic) preclinical development on rug development success

The calculations presented thus far assume that it is possible for orthodox (non-genomic) studies based on cells, tissues and animal models to evaluate every protein in every disease but, in contrast to the genomic approach, this is clearly not feasible. Although numerous orthodox (nongenomic) preclinical programmes, investigating scores of targets at a time, can and do proceed in parallel, the number of such parallel target evaluation programmes is limited by logistics and cost. This imposes the need for a selection step in which a subset of targets must first be prioritized for inclusion in a small number of parallel early phase drug development programmes. By contrast a GWAS for target identification, by definition, interrogates every target in parallel.

This selection step in standard (non-genomic) preclinical drug development therefore introduces a further probability consideration.

The probability that 0,1,2,…*A* causal targets is present in a sample of size *N* (where each member of the sample corresponds to an independent development programme based on a different drug target and encoding gene), drawn without replacement from the pool of 4000 druggable genes (proteins), of which *C* are causal for the disease of interest, is given by the hypergeometric distribution where:

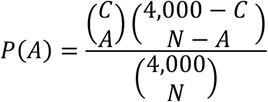

The expected number of causal, druggable targets *E*(*A*) in the sample is given by:

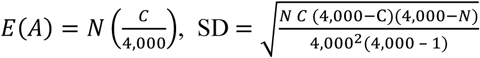

Expected values for *A* based on a range of values of *N* and *C* are shown in **Table S3.** Unless *N* is very large (e.g. 200 independent preclinical programmes proceeding in parallel, each evaluating a different target), there is a very low probability of a causal, druggable target being included in the set selected for preclinical studies, based on a random pick. This emphasises the need for very strong priors before embarking on a drug development programme.

However, there are yet further considerations. Let us assume that a company pursues all *N* targets in parallel preclinical programmes. A true causal target in the sample will have a probability of being correctly detected (true positive rate) corresponding to the power of the relevant experiments (1 – *β*). The probability of a non-causal target being erroneously inferred as causal is given by the experimental Type 1 error rate (*α*). The probability of missing a causal, druggable target is the false negative rate (*β*), while the probability of correctly excluding a noncausal, druggable target (true negative rate) is (1 – *α*). As previously shown in **Figure 3**, for any given disease, the druggable genome can be resolved into components comprising genes that encode causal, druggable targets (previously estimated as around 20 per disease), and druggable but non-causal targets for that particular disease (estimated as 3980). If all *N* parallel preclinical programmes in the sample progress to completion, four outcomes are possible: a) one or more true positives is correctly identified with no false positives; b) a mixture of one or more true and false positives emerge; c) there are no positive findings; or, d) in a worst-case scenario, one or more false positive results emerge with no true positives.

Let us imagine that one nominally positive target is pursued for clinical development under the three scenarios that generate positive findings from preclinical studies (regardless of whether they are true or false positives), and that correct target selection is the only barrier to eventual drug development success (**Assumption 9**). Under the first scenario, clinical development will always be successful, under the second it will sometimes be successful and under the fourth never successful. Consider a thought experiment in which a large number of companies repeat the same process so as to generate a frequency distribution of eventual company successes. The probabilities of eventual development success in this hypothetical drug development world are given by equations in the **Appendix 2** and the results are shown in **Table S5** and **Figure 6.** Assuming there are 20 causal, druggable targets to find, increasing the number of parallel preclinical programmes from 20 to 50 to 200 has a modest impact on drug development success if these are picked from the full set of 4000 druggable proteins. However, if it were possible to obtain *reliable* biological information on the relevance (or not) of selected targets, such that the sampling frame could be reduced in size to 2000, 1000, or perhaps even 200 targets, while retaining all 20 causal targets in the sample, success rates would improve.

**Figure 6** shows the relationship between the expected number of true and false positive findings, the number of causal, druggable targets in the original sampling frame, and the total number of trials. It is relevant that no matter how many parallel drug development programmes are undertaken, the expected number of true positives will only be greater than the number of false positives if the set of targets in the sampling frame is relatively low (< 400 targets) and all causal, druggable targets are retained in the sample. Clinical phase development programmes therefore need to be supported by extremely strong priors. As we argue here, genomic evidence provides compelling biological priors for the full set of 4000 drugggable targets each time a GWAS is done in a particular disease.

**Figure 6.**
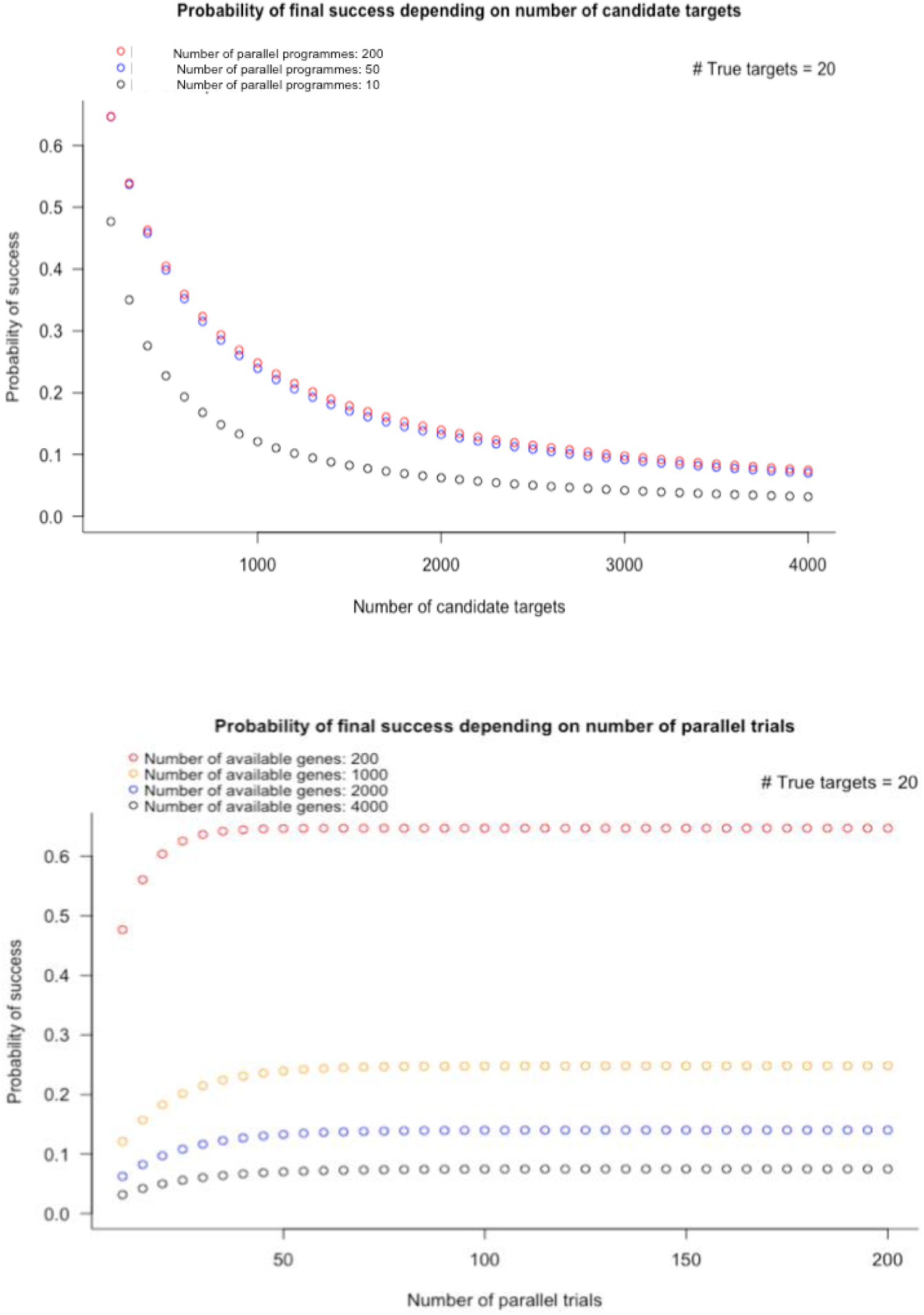
Probability of orthodox drug development success according to the number of candidate targets in the initial sampling frame (upper panel) and the number of parallel preclinical development programmes pursued (lower panel). The calculations assume there are 4000 druggable genes and 20 causal, druggable targets per disease.

Therefore, on the assumption that incorrect target specification is the overarching reason for drug development failure, these considerations go a long way towards explaining the currently low rates of drug development success. They also indicate that the genomic approach to drug target identification should outperform the orthodox non-genomic approach to preclinical drug development at least by several orders of magnitude, even providing the potential to reverse the odds of drug development success.

## Part 3: Assumptions, parameters and limitations

> ‘***Seek simplicity and mistrust it’***
>
> — - **Alfred North Whitehead, In *The Concept of Nature* (1919), Chapter VII, p.143**

> ‘***Your assumptions are your windows on the world. Scrub them off every once in a while, or the light won’t come in’***
>
> — - **Attributed to Isaac Asimov and Alan Alda**

The inferences we have drawn depend on the validity of our assumptions, and on the parameters we used to calculate the various probabilities. We now explore these in more detail before addressing some important limitations.

### Assumptions

**Assumption 1**: *Each gene encodes a unique protein with a single function*

We assumed a 1:1 relationship between genes and proteins, implicitly arguing that any protein has a single function, echoing the historic one-gene one-protein hypothesis of Beadle and Tatum^76^. However, genes can encode alternative mRNA transcripts, some of which may be translated into different proteins^77^. Ensembl (v.87) contains 22,264 protein coding genes encoding 87,662 transcripts. Post-translational modifications increase the complexity of the proteome while some proteins may also contain domains that serve distinct functions^78^. Other proteins, referred to as ‘moonlighting proteins’ appear to have the ability to undertake alternative functions depending on the cellular context, even in the absence of splice variants or distinct functional domains^79^. Moreover, some drugs may interact with a protein-binding pocket composed of elements of two or more protein subunits, each encoded by a different gene. (An example is the benzodiazepine class of drugs that bind to GABA-A receptors at the interface of two of its subunits). Thus, the assumed 1:1 relationship between genes, proteins, protein functions and drug targets, is an undoubted simplification, posing an additional challenge for drug development to not only target the right protein, but also the correct subtype and isoform, sometimes in the right cellular context.

**Assumption 2**: *A given protein can influence the risk of more than one disease*

It has been estimated that nearly 20% of the genes and 5% of the SNPs currently curated by the GWAS catalogue exhibit (pleiotropic) associations with more than one trait^80^ and that many human traits share common genetic influences. ^81 82^ For example, variants in *GCKR* (type 2 diabetes, non-alcoholic steatotic hepatitis, uric acid, glucose, triglycerides), *IL6R* (coronary heart disease, asthma, abdominal aortic aneurysm) and *SH2B3 (*haemopoetic traits, low-density lipoprotein (LDL)-cholesterol concentration, blood pressure, autoimmune conditions, and coronary heart disease) have been associated with diverse diseases and traits. Although the potential mechanisms underlying pleiotropic associations are numerous^83^, one explanation is that a single protein might play a controlling role in several pathophysiological processes. Since a proportion of such genes could encode druggable targets, the corollary is that treatments proven to be effective in one disease have the potential to be successfully repurposed for another. Prior examples of repurposing successes and broadening of treatment indications also support this assumption (**Table 5**). A further consequence is that drugs used to treat one disease could have adverse effects on other conditions, depending on the direction of effect. For example, it is known now that statins, which inhibit HMG-coA reductase reduce the risk of coronary heart disease by lowering LDL-cholesterol. However, they also modestly increase risk of type 2 diabetes, an effect shown by Mendelian randomisation to be mechanism-based^84^. By implication, study designs that interrogate the association of variants in genes encoding a druggable target with a broad range of disease biomarkers and clinical diagnoses in parallel (sometimes called phenome wide association analysis – PheWAS^85^) should offer a systematic and comprehensive means to identify repurposing and indication expansion
opportunities, as well mechanism-based adverse effects. We return to this point in a later section.

**Table 5.** Examples of drug repurposing

| Compound | Target or mechanism | Original indication | Alternative indication(s) |
| --- | --- | --- | --- |
| Thalidomide | Inhibition of vascular endothelial growth factor induced angiogenesis<br>Anti-TNF | Sedative<br>Anti-emetic | Erythema nodosum<br>leprosum<br>Multiple Myeloma |
| Sildenafil | PDE5 inhibition | Angina | Erectile dysfunction<br>Pulmonary hypertension |
| Minoxidil | K-channel opening | Ulcers | Hypertension<br>Hair loss |
| Aspirin | Cyclooxygenase inhibition inhibition | Anti-inflammatory | Antiplatelet |
| Rituximab | Anti-CD20 | Anti-cancer agent for lymphoma | Immunosuppressant for rheumatoid arthritis and SLE |
| Fingolimod | Sphingosine-1-phosphate modulator | Immuno-suppression for transplantation | Multiple sclerosis |
| Abatacept | B7 protein on APCs | Rheumatoid arthritis | Multiple sclerosis |
| Duloxetine | 5-hydroxytryptamine/noradrenaline reuptake inhibitor | Depression | Stress urinary incontinence |
| Imatinib | Tyrosine kinase inhibition | Chronic myeloid leukaemia | Gastrointestinal stromal tumours |
| Beta-blockers | β-adrenoceptor | Angina | Hypertension, heart failure, portal hypertension<br>Infantile haemangiomas |
| Finasteride | 5-α reductase inhibition | Benign prostatic hyperplasia | Male pattern hair loss in males |

**Assumption 3**: *The probability of a protein influencing the pathogenesis of one disease is independent of the probability that it influences any other*

We have shown that even in the presence of this ‘independence’ assumption, it is highly likely that diseases share causal proteins, as supported by evidence from GWAS^82^, providing one explanation for the observation of genetic pleiotropy.

In reality, the independence assumption is very likely to breakdown for certain groups of diseases, with one consequence being that certain disease groups are even more likely to share common targets, offering increased opportunity for therapeutic repurposing. Autoimmune diseases provide some of the clearest examples. As an illustration, monoclonal antibody therapeutics that target tumour necrosis factor-a for treatment of rheumatoid arthritis, also show efficacy in inflammatory bowel diseases^86^. Ustekinumab, a monoclonal antibody that targets interleukin-12/23 receptor developed for psoriasis also shows efficacy in inflammatory bowel disease^87^. Other examples are provided by conditions that might, at first sight, appear to be less likely to share a therapeutic target. For example, monoclonal antibodies targeting vascular endothelial growth factor have found use in the treatment of age-related macular degeneration as well as certain cancers, and it is now known that the pathogenesis of both diseases involves angiogenesis^88^. However, such agents also raise blood pressure and increase risk of thrombotic vascular events as a consequence of their mechanism of action^89^.

If diseases related by common mechanism were to be grouped as adjacent columns in the sample space (**Figure 1**), and the genes encoding functionally related proteins as adjacent rows, with the sample space being marked using contours corresponding to probabilities of any target-disease paring being disease-causing, then ridges and troughs of higher and lower probability would be observed to emerge from an otherwise flat, homogenous probability space that corresponds to the independence assumption. In due course, we believe the genetic approach we describe will uncover more diseases with common underpinning, that this will enable reconfiguration of gene and disease relationships in the sample space, and will support more rational medication repurposing and indication expansion programmes^90^.

Nevertheless, at present, given the very broad spectrum of human diseases, we consider our simplifying assumption to serve as a useful start point for the concepts we develop and calculations we make.

**Assumption 4**: *Drug treatments for human disease target proteins encoded in the germ line.*

We excluded from consideration the treatment of many infectious diseases, where proteins in the pathogen rather than the host are the therapeutic targets, as well as cancer, where treatment targets are mutated or aberrantly expressed proteins encoded by the abnormal genome of the cancer cell. However, with these restrictions, proteins encoded by the germ line serve as the therapeutic targets of >80% of licensed drugs^91 92^ This simplifying assumption is therefore robust for the sample space as we define it.

**Assumption 5**: *DNA sequence variants in and around a gene encoding a drug target, that alter expression or activity of the encoded protein (cis*-*acting variants) are ubiquitous in the genome*

GWAS of mRNA expression and protein concentration provide hundreds of empirical examples of SNPs influencing the expression of nearby genes (acting in *cis)* leading to the concept of expression (e) and protein (p) quantitative trait loci (QTL)^93 94 95 96 97 98^. Recently, the ENCODE, ROADMAP and GTEX projects have catalogued variants with functional effects on both local (*cis*) and distant (*trans*) gene expression in a variety of cell types and tissues^99 100 101^. As datasets enlarge and improved proteomics platforms encompass a broader set of human proteins, we anticipate the catalogue of *cis* pQTLs will expand, providing a larger armamentarium of such variants
in genes encoding druggable targets that serve as important tools for drug target identification and validation.

**Assumption 6**: *The association of cis*-*acting variants with biomarkers and disease end-points in a population genetic study accurately predict the effects of pharmacological modification of the encoded target in a clinical trial*

The reliability of this assumption has been demonstrated by comparisons of the associations of *cis*-acting variants in genes encoding the targets of licensed drugs in population studies, and the effect of treatments targeting the same protein in clinical trials, using a common set of biomarkers and disease outcomes as the readout. Applied examples of this paradigm have now been used to predict the eventual failure in clinical trials of first-in-class drugs for prevention and treatment of cardiovascular disease^102 103^, to separate on- from off-target effects of drugs^84 104^, and to identify indication expansion opportunities for established drugs ^105^. This concordance may seem surprising given that drugs typically target the *action* of proteins while variants identified by GWAS are typically non-coding, probably influencing mRNA and thence protein *expression.* Nevertheless, the empirical findings are compelling, with recent studies indicating that the concordance between the effects of genetic variants and drugs targeting corresponding proteins can extend across scores of biomarkers and disease end-points^106^. These proof-of-concept examples (**Appendix 1**) now provide strong motivation for scaling the approach to interrogate the association of *cis*-acting variants in all druggable genes against the full spectrum of diseases and biomarkers in parallel.

Coding region (loss- and gain-of-function) variants have also been shown to be useful tools for drug target selection and validation^107 108^. As falling costs lead to an expansion in sequencing studies, including in populations with a high rates of consanguinity, thereby enriched for homozygous loss of function variants^109 110^, we also anticipate that a broader spectrum of druggable genome variation will be discovered encompassing rare, low frequency and common variants in both coding regions (influencing function) and non-coding regions (influencing expression) that, when linked to phenotype and disease outcome, will provide invaluable information for target identification and validation.

**Assumption 7**: *Genotyping arrays used in GWAS provide comprehensive, appropriately powered coverage of the genome, and associations discovered at any one gene are independent of those detected at any other*

We have made the assumption that the genotyping arrays used in GWAS provide comprehensive coverage of all genes (including all druggable genes), that all such studies are conducted such that power is 0.8 at all loci, with *α* = 5 × 10^−8^, and that the discovery of any one genetic locus is independent of any other. We recognise that in reality, power in many GWAS is likely to be much lower than 0.8 suggesting that additional loci are likely to be identified by increased sample size. We also recognise that the local correlation between SNPs (linkage disequilibrium; LD) can lead to ambiguity on the source of the association signal(s) at any locus identified by a GWAS (placing uncertainty on the role of any implicated drug target). We showed previously that GWAS to date have identied LD regions containing a single druggable gene in around 10.5% of cases^67^, and 31.9% of such LD regions contain 2 or more genes, at least one of which encodes a druggable target. However, to begin to address the issue of verifying the causal gene(s) in an associated region, sequencing projects have led to haplotype reference panels that enable dense imputation and fine mapping of association signals^111^. *In silico* approaches based on functional annotation of the genome have been developed, as have statistical-, pathway-, and eQTL-co-localisation methods, to address this problem, together with scoring systems that assimilate results from multiple methods with various degrees of weighting^112^. An alternative approach to elucidation of causal signals with translational potential is to flip the problem by focusing genetic association studies exclusively on *cis*-acting variants within the druggable genome – ‘druggable genome wide association studies’. To that end, we recently designed the content of a new genotyping array, with dense marker coverage of genes encoding druggable targets^67^, facilitating a gene-centric approach to disease association studies for drug development. The array design also enables gene-based, not just SNP-based, association tests. The inclusion of common, non-coding as well as less frequent coding variation, should also enable the construction of allelic series^113^ (the genetic counterpart of a pharmacological dose response relationship).

**Assumption 8**: *The probability that a protein affects disease pathogenesis and the probability the protein can be targeted by a drug is independent*

This assumption is more speculative. An argument could be made that genes included in our recent update of the druggable genome^67^ that encode the protein targets of small molecule drugs are more likely than other genes to be disease causing. This is because druggability predictions are based, in part, on membership of protein families containing licensed drug targets that, by definition, are both druggable and play a controlling role in disease susceptibility. However, this bias should not apply to the 2000 or so genes that were included in the druggable set because of sequence similarity to drugged proteins, or because they encode extracellular regions that are targetable by monoclonal antibodies^67^. Moreover, the converse argument is equally plausible that druggable genes are less likely than others to be pathogenic, because the druggable set is enriched for proteins with natural ligands that sub serve key cellular functions. Evolutionary forces might therefore exert purifying selection on deleterious variants in such genes, if they were to affect reproductive fitness. Empirical evidence on this issue is limited. In our own recent analysis using findings curated in the GWAS catalogue^67^, we find that the proportion of druggable genes present in regions of LD with disease-associated SNPs is an approximately constant proportion of all genes present in such regions, that this is consistent across disease categories, and close to the proportion of druggable genes in the genome overall (i.e. ~4000/20,000 = 0.2). This would be expected if disease association and druggability were independent. However, others have found an apparent enrichment of druggable genes among disease-associated loci ^73^. We expect this uncertainty will be resolved as more GWAS are undertaken in a wider range of diseases with the purpose of drug target identification and validation.

**Assumption 9**: *Inaccurate target selection is the exclusive reason for clinical phase (stage 2) drug development failure*

Drug development can fail for numerous reasons including idiosyncratic compound toxicity, incorrect dosing, unfavourable pharmacokinetics, incorrect end-point selection, mechanism-based adverse effects and commercial considerations. Nevertheless, recent reviews have documented lack of efficacy (despite adequate target engagement) as the reason for clinical phase drug development failures in around two-thirds of cases ^24 25 28^. With this asssumption, we will have over attributed failure due to inaccurate drug target selection. However, adjustment of the relevant estimates by the multiplication factor of 2/3 (to account for other reasons for failure) would not overturn our broad conclusions, given the orders of magnitude improvement in developmental success rates predicted from the genomic approach.

### Parameters

We estimated several key parameters when making our calculations. Here we review their likely accuracy.

*Number of human protein-coding genes:* As summarized in **Box 4**, recent estimates of the number of protein coding genes, derived from diverse sources of evidence, have settled to a figure of close to 20,000.

*Number of complex diseases:* We recognize that it is problematic to define diseases based on the use of coding schemes such as ICD-10^114^, utilized primarily for billing and record keeping, which offer a finite list of possible disease options, and which classify disease mainly according to appearance rather than cause. We also recognize that an ultimate outcome of research on the genetic basis of human disease may be the reclassification of disease according to molecular mechanism rather than appearance. As diseases often lie on a spectrum, with overlaps in both disease phenotypes and genetic causation, defining discrete disease entities often involves a degree of subjectivity. In the post-genomic era, biomedical ontologies have been created to provide controlled terms for biological attributes. The emphasis of coverage in the Human Phenotype Ontology (HPO) is on phenotypic abnormalities and clinical observations rather than diseases, while the Experimental Factor Ontology (EFO) describes experimental variables from the cellular to disease level in the European Bioinformatics Institute (EBI) databases. The Human Disease Ontology (DO) is a biomedical resource of standardised disease concepts organised by disease aetiology. It addresses the complexity of disease nomenclature through extensive cross mapping and integration of ICD, Online Mendelian Inheritance in Man (OMIM), Orphanet, EFO, National Cancer Institute (NCI) Thesaurus, SNOMED CT and MeSH concepts^115 116^. As of 20 January 2016, the DO had 9,196 terms. The number of terms in the DO is regularly updated with technical and conceptual advances in disease phenotyping and will increase with improved understanding of molecular pathways. Therefore, given the current state of knowledge, we propose that a figure of 10,000 is a reasonable estimate of the number of common human diseases with genetic susceptibility. However, we explain in earlier sections why the various probabilities we have estimated do not depend on the absolute number of disease entities under consideration.

*Number of susceptibility genes for common diseases:* Estimating a reasonable figure for the number of susceptibility genes for common diseases is a critical parameter when estimating probabilities of drug development success and requires consideration of the genetic architecture of these conditions ^54 55 117 118 119^. This area is controversial, as reviewed by Gibson^120^, and recently by Pritchard^54^. The approach we took in this article implicitly accepts the front-running, common-variant, common-disease hypothesis, which states that complex diseases and associated biological traits are determined by the additive (perhaps occasionally synergistic) action of common, small effect variants in a large number of human genes. Under this model, every individual carries a different repertoire of largely independently inherited variants. (This model also has implications for the success or otherwise of precision medicine therapies).

The diametrically opposed hypothesis is that the association of multiple SNPs at any locus with a disease or trait seen in GWAS occurs exclusively because common SNPs mark the presence of unobserved, rare (large effect) variants present in subsets of the population (a phenomenon referred to as ‘synthetic association’)^121^. Rare variants of this type are under-represented in the commonly used genotyping arrays used in GWAS, may be difficult to impute from haplotype reference panels, and should be better captured by exome or whole genome sequencing.

However, evidence from post GWAS fine mapping studies, and a recent report on the genetic architecture of type 2 diabetes, in which whole genome sequencing allowed an unbiased survey of both common and rare variant effects in tandem, continues to provide evidence for the common variant common disease hypothesis^122 123 124^. However, it is also clear that rare, or infrequent, large effect, coding variants can also coexist in any given gene. Evidence from GWAS and emerging sequencing studies also suggest that a very large number of loci contribute to susceptibility to most common diseases and biomedical traits, but that the sometimes hundreds of loci exerting the largest effect, detected most readily by GWAS, explain only a small fraction of the heritability, with the remainder perhaps being distributed across the many thousands of remaining genes throughout the large expanse of the genome. This ‘omnigenic’ could be inaccurately interpreted as ‘all genes contribute (equally) to all diseases’. However, effect sizes at loci beyond ‘core’ (or ‘critical’) genes may be beyond detection even by massive expansion in sample sizes^120^. Moreover, even allowing for development of highly potent compounds against ‘peripheral’ targets, the biological effect may still be too small to be of therapeutic interest and might necessitate unfeasibly large clinical trials for any effect to be reliably detected. For this reason, we believe the concept of scores or hundreds of causal (‘critical’, ‘core’) genes for any disease, i.e. those with the main effect, is still valid.

We estimated the number of such genes for a given disease using information from published GWAS of common diseases with the largest available datasets. These have typically identified hundreds of genetic susceptibility loci. As it is conceivable that even more loci will be uncovered by further increases in sample size^125^, we also estimated relevant probabilities for 1000 ‘causal’ genes per disease (corresponding to around 200 druggable genes per disease). We consider a further 10-fold increase in the number of causal genes (to 10,000 genes per disease in total) is unlikely, if only because the observed rates of drug development failure from lack of efficacy would be difficult to explain if half of all genes in the genome (corresponding to 2000 of the 4000 druggable genes under **Assumption 8**) critically affected risk of any given disease.

*Size of the druggable genome:* A historical perspective of the druggable genome was provided in **Box 6**. We recently re-estimated the extent of the druggable genome based on up to date annotations of protein coding genes, information on protein motifs targeted by drugs that have been licensed since prior estimates of the druggable genome were made, and by incorporating predicted targets of monoclonal antibody therapeutics which are either membrane-bound or secreted proteins identifiable by specific motifs in their primary structure. This estimate of approximately 4479 druggable, protein-coding genes was used to inform the content of a new genotyping array developed specifically to facilitate genetic studies for drug target identification^67^.

This figure was rounded (conservatively) to 4000 genes for the illustrative calculations used in the current paper. We recognize this estimate is not fixed but likely to be revised with time as new therapeutic modalities are developed^126^, evidenced by recent clinical successes of RNA therapeutics^127^, of gene therapy^128^, and of gene editing technologies that may play a therapeutic role in certain rare disorders^129^. However, we believe it is a reasonable first approximation that drugs that act by interfering with the action of proteins readily target only a subset of human gene products, and that the factors that determine whether a protein is druggable and whether it plays a controlling role in a disease are somewhat distinct. This echoes the arguments made by others^65^, that the challenge in drug development is to identify the proteins that lie at the intersection of druggability and disease regulation, and that human genomics is in a unique position to delineate this set of proteins for each disease of interest.

### Limitations

There are a number of limitations to our analysis.

We have argued that *cis*-acting variation is widespread in the human genome, but it may not be universal. In the absence of natural variation in a gene encoding a drug target of interest, influencing its expression or activity, it would be impossible to use the approach described to anticipate the pharmacological action of a corresponding drug. However, there may be ways of addressing this issue in the infrequent instances where this occurs. For example, in the absence of variants reliably influencing expression of the gene encoding interleukin-6, variants in the gene encoding the interleukin-6 receptor were used to model the effect of interference with interleukin-6 signaling on coronary heart disease risk, through pharmacological blockade of the receptor rather than the ligand^105^.

Theoretically, since genetic influences on protein expression or activity are present from early life, they may entrain developmental adaptation (canalization) through changes in other pathways that mitigate any biologically adverse effect on the system as a whole^66^. Thus, the null association of variants in a gene encoding a drug target of interest in a particular disease need not completely exclude it as a therapeutic target. This is because drugs, particularly for common diseases, are administered late in life, when developmental adaptation is inactive. Yet there are now numerous instances of both common (small effect) and rare (large effect) variants in genes encoding druggable targets that reliably anticipate the effects of drugs for late life diseases (see **Appendix 1**). Thus, it would seem that canalization is a more theoretical than practical consideration for genomic identification and validation of therapeutic targets.

We have observed that *cis*-acting variants in a gene encoding a drug target can anticipate both the pattern and rank order of effects of the corresponding drug on disease biomarkers. However, the effect sizes observed, particularly with common genetic variants, are typically one fifth to tenth that of the cognate drug. Thus, there remains the possibility that if certain biological actions are only observed beyond some threshold, achieved through target perturbation by a potent drug, but not by the weak effect of natural genetic variation, such variants will fail to anticipate the full spectrum of effects of drug treatment. Thus, any discrepancy in the effects of genetic variants and drug action might arise not only from off-target actions of a drug (not shared by natural genetic variation), but also because of on-target threshold effects. The availability of common (weak effect) and rare (large effect) genetic variants in the same gene, that allows the construction of an allelic series (effectively a genetic dose-response curve), may go some way toward mitigating this possibility in specific cases^65 113^.

We noted previously that local correlation between SNPs (LD) might lead to ambiguity on the source of the association signal(s) at any locus. Since LD can extend beyond gene boundaries, this issue can affect gene-centric as well as whole genome association studies, though perhaps less so. In such gene-centric studies, there remains the possibility that disease and biomarker associations attributed to the local gene of interest in fact arise from effects of adjacent genes. Approaches for exploring and accounting for this possibility were discussed earlier. The genomic approach to target identification and validation we describe is also necessarily limited by the range of available phenotypes. Failure to comprehensively capture phenotypes influenced by perturbation of the target of interest, could lead to incomplete anticipation of the effect of drug treatment. Recently, the monoclonal antibody romosozumab targeting sclerostin for the treatment of osteoporosis was developed based on the observation that patients with rare mutations in the encoding gene have increased bone mass. This agent increased bone mineral density and reduced osteoporotic fracture rate in two phase 3 randomised trials but, in one of the trials, the rate of serious adverse cardiovascular events was also increased^130 131^. Since prior genetic studies, which had focused mainly on patients with rare mutations, had not evaluated cardiovascular end-points, it remains uncertain whether the apparent adverse signal of cardiovascular safety is real and if so, whether it is an on- or off-target, or threshold effect.

Finally, most common disease genetic association studies that might inform drug development that have been performed to date have been undertaken in population-based longitudinal cohorts or case-control control datasets, where cases typically represent the first occurrence of a condition (e.g. a coronary heart disease event). However, first-in-class agents for CHD, and for many other common conditions, are tested or used initially patients with established disease, for prevention of disease progression or recurrence^132^. Mendelian randomization studies for target identification and validation in longitudinal clinical cohorts with established disease are few, currently limited by the available datasets, and also perhaps by potential biases arising from survivorship of, or indexing by, an initial event, that may limit inferences that can be drawn^133^. Nevertheless, the rediscovery by GWAS of over 70 drug targets suggests that genes influencing disease onset can, in many (but perhaps not all) cases, provide useful insight on targetable pathways for prevention of progression or recurrence of common conditions.

In our *a priori* and *a posteriori* calculations of *γ_pc_* and other relevant metrics, we artificially reduced drug development to two steps: a preclinical component to predict target-disease pairings destined for clinical phase success (stage 1), and a clinical component (stage 2) to evaluate target-disease pairings brought forward from stage 1. The approach allowed the generation of formulas that highlight the key variables influencing drug development success, and some estimates of their values, based on observed success rates. These calculations should be viewed as no more than an illustration to help inform developers of the key variables influencing success rates.

## Part 4: Summary and implications for drug development

> “***Knowing is not enough; we must apply. Willing is not enough; we must do.***”
>
> — —**Johann Wolfgang von Goethe, Writer and Statesman (1749-1832)**

### Summary

Three crucial factors have conspired to inhibit drug development success:

a. The apparently widespread contamination of the scientific literature by false discoveries, which undermines the validity of the hypotheses used to prioritise the selection of drug targets for different diseases;
b. The poor predictive accuracy of orthodox preclinical studies, arising due to animal-human differences in pathophysiology; and
c. The system flaw in drug development that sees the definitive target validation step (the RCT) deferred to the end of the drug development pipeline.

With reasonable assumptions about the number of protein coding genes, druggable proteins and human diseases, and using probabilistic reasoning, we estimated that the observed success rate in drug development (~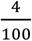 for compounds; ~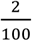 for targets) only marginally exceeds the probability 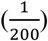 of correctly selecting a causal, druggable protein-disease pair through a random pick from a sample space defined by the 4,000 genes that are predicted to encode druggable targets and 10,000 diseases, assuming an average of 100 causal genes per disease. With a target success rate of 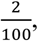 based on the orthodox (non-genomic) approach to target selection and validation, over 100 independent drug development programmes for each disease need to proceed in parallel to have a 90% probability of even one success.

Based on reported clinical and preclinical success rates, and making reasonable assumptions about values of clinical phase type 1 and type 2 error rates (*α_C_* and *β_C_*), we also found evidence that the proportion of true target disease relationships studied in preclinical development is small, that these form only the minor proportion of nominally positive findings that are brought forward into clinical phase studies. This likely contributes to the high preclinical false discovery rate and low clinical phase success rate.

Even applying the assumption that the probability of a protein influencing the pathogenesis of one disease is independent of the probability of it influencing any other, we show that it is highly likely that even small groups of diseases taken at random share at least one common target. This implies numerous opportunities should exist for therapeutic repurposing, but also that even highly specific modification of any target still runs a high risk of mechanism-based adverse effects. However, knowledge of the effect of target-specific perturbation on multiple disease outcomes currently remains incomplete because the orthodox approach to target identification and validation is neither systematic nor comprehensive.

In contrast to established non-genomic, approaches to preclinical drug development, GWAS deliver a methodical and reliable means of specifying the correct drug targets for a disease, provided that the genotyping arrays that are deployed have sufficient coverage of the druggable genome, and that the studies are adequately powered. GWAS differ from established non-genomic preclinical experiments for target identification in that the evidence source is the human not an animal model; the false positive (type 1) error rate is low (typically set at 5 × 10^−8^); every potential drug target is interrogated in parallel (not just a selected subset); and the study design shares features of an RCT, the pivotal step in drug development. For these reasons, we suggest that genetic studies will soon be universally regarded as an indispensable, though not exclusive element of drug development for common diseases. By improving the efficiency and reliability of target identification, GWAS and similar genetic study designs offer the potential to overturn the currently poor odds of success currently beleaguering drug development.

However, GWAS have yet to be optimally designed or sufficiently widely deployed to fully realise their potential to uncover the correct drug targets for many poorly treated diseases. There are several reasons for this that relate to the design of genotyping arrays used in GWAS, the range of diseases studied, and the datasets used.

*Design of genotyping arrays used in GWAS:* Genotyping arrays used in GWAS to date have been designed to provide broad coverage of the human genome, while other widely used genotyping arrays were designed to fine-map disease-associated loci identified by prior GWAS. Neither design focuses explicitly on genes encoding druggable targets. In whole genome arrays, local coverage of variants in genes encoding druggable targets could be sparse, while in fine-mapping arrays such coverage could be incomplete. For this reason, we recently specified the content of the Illumina DrugDev consortium genotyping array that combines the properties of a whole genome array with focal coverage of variants in the druggable genome to support genetic association studies for drug target selection and validation (‘druggable GWAS’) ^67^.

*Diseases represented in GWAS:* The 400 or so unique diseases and biomarkers subjected to GWAS so far represents only a fraction of the thousands of terms listed by disease classification systems or ontologies, or that are observed in electronic health record datasets. Moreover, retrospective power calculations suggest that sample sizes in many GWAS to date may have been insufficient to detect all causal, druggable targets. Despite this, more than 70 of the 680 or so known drug targets have already been ‘rediscovered’ based on therapeutic indications or mechanism-based adverse effects, signposting the future potential of this approach in drug development.

*Datasets used in GWAS:* Datasets subjected to GWAS up to now have typically been conducted one disease at a time. Yet, when information from such studies is collated, it becomes apparent that the same loci, genes or even SNPs can contribute to the susceptibility to more than one disorder, a phenomenon referred to by geneticists as ‘pleiotropy’. Pleiotropy can arise through a number of mechanisms^83^, but an important one for drug development is the involvement of the same encoded protein in the pathogenesis of more than one disease, flagging potential opportunities to repurpose therapies effective in one disease for another. In this paper, we estimated that a single gene (and thereby a single druggable target) could affect the risk of 50 different disease entities. Undertaking GWAS one disease at a time and cross-referencing findings later is a relatively inefficient method for pleiotropy detection. An alternative approach to pleiotropy detection at druggable targets is to undertake phenome wide association studies (PheWAS) using extremely large prospective cohorts, or genomic studies within healthcare systems. Though there is emerging activity in this area, there is much yet untapped potential.

### Implications for future drug development

The concepts and calculations in this paper suggest avenues by which drug target selection and validation, and hence drug development success, might be improved in the future, even if a complete reversal of the odds of drug development success is only a theoretical rather than practically achievable goal.

First, more systematic mining should be undertaken of data emerging from GWAS for the purposes of drug target identification. Several groups, including our own, have initiated such work^67^ and new initiatives such as MR Base^134^ and Open Targets^135^, and commercial spinouts (e.g. Genomics PLC) suggest there is a growing interest in this area.

Second, more systematic and comprehensive genomic studies of high priority targets could be undertaken prospectively against as broad a range of biomarkers and disease end-points as possible (drug-target PheWAS) to facilitate drug target validation.

Third, to realise the full potential of genomics for both drug target identification and validation, genomic studies with comprehensive coverage of variation in the druggable genome need to be conducted at even larger scale, and with attention to multipe (not just single) biomarker and disease outcomes - joint genome- and phenome-wide association analyses (**Figure 7**). This ‘big data’ approach requires resources that couple comprehensive genomics with extensive phenotype and disease capture. One route to achieving this is to pull together analyses across cohorts, consortia and large national biobanks, and there are emerging examples of this approach^136^. Cohort consortia and large national biobanks can also exploit their ability to undertake and evaluate emerging technologies in detail (e.g. transcriptomics, epigenomic, proteomic and metabolomic measures in tissues, blood and urine). Summary level genetic associations with mRNA and protein expression, with metabolite level and with disease risk obtained in different datasets can subsequently be connected by a variety of statistical methods, to elucidate pathways to disease, because natural genetic variation (unaffected by disease and allocated at random) provides a fixed anchor point with which to connect such datasets, exploiting the central dogma^137^ of the unidirectional flow of information from DNA to RNA, to protein and via downstream mechanisms to disease. It should be possible in this way to gain comprehensive insight on mechanism and pathway, as well as the likely downstream consequence of targeting a druggable protein pharmacologically.

**Figure 7.**
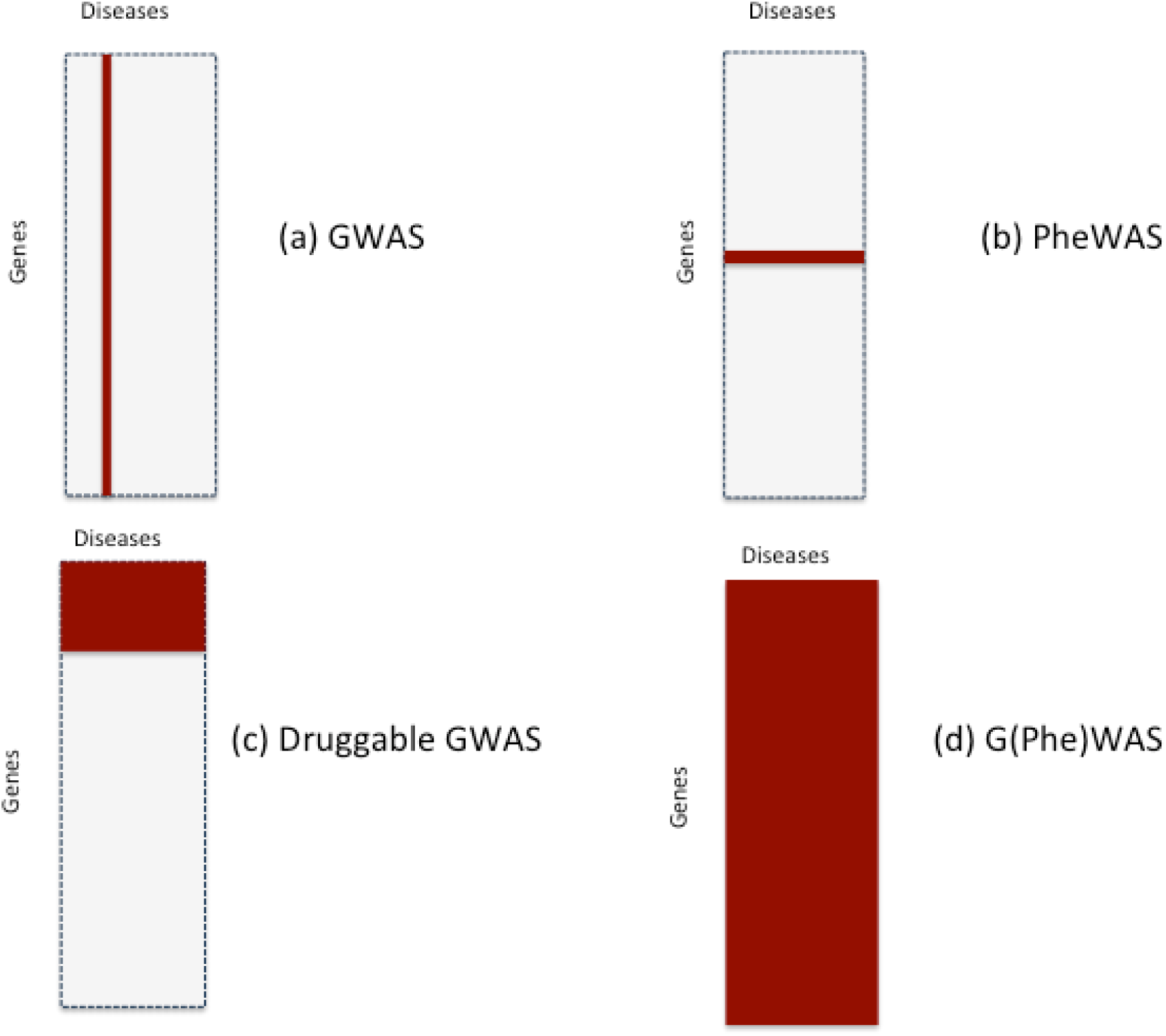
Study designs relevant to drug target identification and validation based on human genomics: (a) conventional genome-wide association analysis in which variation in 20,000 genes is tested against a single disease; (b) phenome wide association analysis of a gene encoding a drug target in which variation in a single druggable gene is evaluated against many (all) diseases; (c) druggable genome and phenome wide association analysis; and (d) whole genome and phenome wide association analysis

However, we believe a further step to increase the scale, breadth and depth of the approach is to embed genomic analysis within the healthcare setting so that information on natural genetic variation could be linked to the multiplicity of clinical and disease outcome data ascertained during routine clinical care^138^.

To achieve a shift in development of this type, the benefits need to be clear to healthcare providers (whether insurers or governments), to academia and industry and, most importantly of all, to patients and society, addressing legitimate concerns that might exist about privacy, security and secondary use of health data.

If such concerns can be addressed, through rigorous governance and data security, a new model of drug development might supervene because healthcare data typically resides outside the domain of the pharmaceutical industry within the healthcare sector, which, in some countries, is wholly or substantially state-run.

In turn, this would dictate that a new funding and delivery structure might need to be established, at least for the component of drug development that relates to target identification and validation.

We explore these issues in greater detail.

*Healthcare genomics as a means to increase the scale and range of gene-disease associations to improve drug target identification:* Most datasets used in prior GWAS have either been investigator-led collections of patients with single diseases or population based cohort studies. Efforts to expand existing studies or to make new disease collections proceed sporadically because they are expensive to undertake and unattractive to research funders given that the initial creation of the dataset, no specific scientific hypothesis is explored. Population cohort studies measure numerous preclinical biomarkers and are increasingly being enriched with new proteomic and metabolomics measures. However, only relatively modest numbers of cases of different disorders accumulate in such datasets over time, determined by their natural incidence rates. Consortia of cohort studies, and large national biobanks^139^ have gone some way towards achieving the necessary scale but we believe a further step-change is now needed to maximise the value of genetic studies for drug target identification.

Based on the arguments developed in this paper, we propose that genotyping or eventually sequencing be embedded in routine healthcare settings to explicitly aid target identification and validation for drug development. This is because routine diagnostic and prognostic tests are undertaken, and clinical diagnoses made in patients (as well as healthy citizens as part of preventative medicine efforts) on a scale and with a range that would be challenging to reproduce using investigator-led case collection or cohort studies in the conventional research setting. Indeed, in countries with healthcare systems that provide universal coverage such as the National Health Service in the UK, the theoretical cohort size extends to the whole population (63 million people in this example), which would encompass disease collections of unsurpassed size and breadth. Were such healthcare datasets to be connected to information on genetic variation, even at summary level, the genotype-disease associations that would be gathered would enable drug targets to be matched accurately, systematically and efficiently to the multiplicity of diseases occurring in such healthcare settings, with the bonus of capturing multiple disease outcomes in the same individual.

There are already precedents to using healthcare data at this scale for research. In the UK, the Clinical Practice Data Link^140^ has provided anonymised primary care records for research since 1987 and, more recently, CALIBER^141^ has created a research cohort of ~10 million individuals by linking health records from primary care, hospital episodes, disease registry and mortality statistics. Mature efforts to utilise routine healthcare data for research have also been established in Scandinavia and elsewhere^142^. In the USA, precedents have already been set for connecting genotyping data to healthcare records to help identify disease-susceptibility and treatment response genes, e.g. in the EMERGE consortium^143^ and the Million Veterans Programme^144^. In the UK, information on genome sequence is being connected to health record data in UK Biobank, in patients with rare diseases through the Genomics England (GEL) project^145^ and in individuals from ethnic minority groups with a high prevalence of certain diseases and a high degree of relatedness through the East London Genes and Health Initiative^146^.

A national healthcare genomics effort would build on and complement these efforts. It would extend research platforms based on electronic health records alone (e.g. CPRD and CALIBER) into the genomic space. It would surpass the scale and representativeness of existing genomics healthcare platforms or initiatives (e.g. EMERGE or Geisinger, which have been in the vanguard of these developments, but which are confined to participating private healthcare systems) as well as the Million Veterans Programme which, through its design, includes almost exclusively male participants (see **Table 6** for other examples of genomics and healthcare initiatives). Moreover, unlike GEL and the East London Genes and Health project, where recruitment is highly targeted, a national genomics effort would receive all comers. Until costs fall further and informatics pipelines are more streamlined, it could also focus on genotyping using fixed content arrays, exploiting increases in the number of genotyping assays per array and improved reference panels for imputation. This approach would be less expensive and less analytically demanding than whole-genome or whole-exome sequencing. As sequencing eventually becomes more cost-efficient this technology would eventually replace genotyping.

**Table 6.** Selected examples of Academia, Pharma, and Pharma-Academia initiatives concerning genomics and drug development

| Initiative | Partners | Drug development model | Aims |
| --- | --- | --- | --- |
| **Accelerating Drug Development and Repurposing Incubator at Vanderbilt University<sup>a</sup>** | Multiple departments at Vanderbilt University Medical Centre | Academic incubator | De-identified genotype data linked to de-identified demographic and health record data to aid precision drug development and drug repurposing |
| **DECODE Genetics<sup>b</sup>** | Decode is a subsidiary of Amgen, a biopharmaceutical company | Within-company | Discover genetic variation underlying human disease in the Icelandic population with the aim of diagnosing, treating and preventing disease |
| **Open Targets<sup>c</sup>** | GSK, Biogen, European Bioinformatics Institute, Wellcome Trust Sanger Institute | Pre-competitive, open access | Public-private initiative based on the use of genomics for drug target validation |
| **Astra Zeneca Centre for Genomics Research** | Human Longevity, Inc<br>Wellcome Trust Sanger Institute<br>Institute for Molecular Medicine, Finland | Within-company | 'Integrated genomics initiative to transform drug discovery and development across (AZ's) entire therapeutic pipeline' |
| **Eisai Andover Innovative Medicines Institute<sup>e</sup>** | Seeking collaborations with external scientific partners | Pre-competitive research consortia | 'Executing novel therapeutic targets validated by human genetics' |
| **Regeneron Genetics Centre<sup>f</sup>** | Geisinger Health System, and other health service and academic partners | Within-company | 'Comparing genetic information against medical histories .to develop new means of diagnosing, preventing and/or treating medical conditions' |
| **GSK-Regeneron UK Biobank Partnership<sup>g</sup>** | GSK, Regeneron and UK Biobank | Industry academia partnership, with 9 month exclusivity period for Pharma partners | Exome sequencing of stored DNA from UK Biobank participants: 50,000 samples in year 1, 500,000 by year 3. |

<sup>a</sup> http://online.liebertpub.com/doi/10.1089/adt.2016.772
<sup>b</sup> http://www.decode.com/
<sup>c</sup> https://www.opentargets.org/
<sup>d</sup> https://www.astrazeneca.com/media-centre/press-releases/2016/AstraZeneca-launches-integrated-genomics-approach-to-transform-drug-discovery-and-development-22042016.html
<sup>e</sup> http://us.eisai.com/research/andover-innovative-medicines-institute
<sup>f</sup> https://www.regeneron.com/genetics-center
<sup>g</sup> http://www.ukbiobank.ac.uk/2017/03/gsk-regeneron-initiative-to-develop-better-treatments-more-quickly

The optimal mechanisms for obtaining consent, for biospecimen collection, and for data management would need to be established, but much could be learnt from preexisting efforts. For example, bio-specimen collection might occur in hospital (at the point of emergency or elective care, during imaging or blood taking), in primary care, and / or by a direct-to-patient approach, using a despatched saliva collection kit, or some combination.

*Justifying a healthcare genomics initiative to healthcare providers and users:* The full engagement, understanding and support of patients and healthcare providers would need to be gathered at scale, with an open dialogue about the potential risks (e.g. of unintended patient data disclosure) balanced against individual and societal benefits. Recent enterprises such as the Transforming Genetic Medicine Initiative (TGMI)^147^, the Personal Genome Project^148^ and Patients Like Me^149^ may have an important role to play, if the ideas are to gain traction.

Healthcare providers and users might at first consider any potential research benefits from the initiative we describe to be too speculative and the benefits too remote. However, we believe the arguments elaborated in this paper and elsewhere make the overall scientific and economic case compelling. Moreover, evidence is already emerging that genomics has whet the public appetite for wider participation in medical research. For example, direct to consumer genotyping has been available for some time through 23andme and other providers^150^, including distribution through high street outlets. Participants submitting samples to 23andMe outside the UK have had the opportunity to participate in medical research by submitting self-reported healthcare data. Such information has already contributed to disease gene discovery in Parkinson’s disease^151^, depression^152^ and a range of other diseases and traits^153^. Similar efforts are being made by the academia led Genes for Good collaboration^154^. It seems not a very great leap of faith to consider that, with appropriate public discourse on potential benefits, and mitigation of any risks, that there could be widespread public enthusiasm for an initiative that explicitly links anonymised personal genomic data to health records for the purpose of accelerating drug development, under a new model, to the benefit of society.

Since healthcare providers and users might still rightly argue for more immediate and individual benefit from a healthcare genomics initiative, the genotyping arrays for this project could be designed with a dual purpose. Genotypic information of immediate value in healthcare decision-making could be made available to patients and their doctors as part of a healthcare episode: *personal healthcare genomics for diagnosis*, *risk assessment and individualised treatment*. This could include information on clinically actionable genetic variants that influence beneficial and adverse drug response, disease risk^155^, compatibility of transfusions and transplants, or risk of recessive genetic diseases that might manifest in future generations, to aid preconception planning, as such variants become sufficiently validated. Validated genotypic information from prior GWAS of general interest to patients could also be returned, e.g. on ancestry; on genes influencing sleep pattern, facial appearance, hair and eye colour, coffee and alcohol metabolism and so on. In parallel, the remaining genomic information from participants, linked *anonymously* to health record phenotypes and disease outcomes, would contribute *in aggregate* (at *summary <u>not</u> individual* level) to large-scale investigations of the causes of human diseases and the identification of disease-specific drug targets: *public health genomics for drug development.*

*Democratising drug development:* If accepted, an effort such as this would be likely to convert drug target identification and validation from an almost exclusively private sector, commercially sensitive enterprise to an open, pre-competitive, societal endeavour, with the joint involvement of academia and industry, healthcare providers and healthcare users, all with the shared goal of developing new medicines more efficiently. In effect, drug development would become democratised; with healthcare users also becoming participants in drug target discovery and validation.

If new medicines are to arise from this endeavour, there would still need to be intellectual property and revenue opportunities for commercial partners. The biotech and e-tech industries could be engaged to develop and deploy the optimal tools for bio-sample collection, genomic analysis, data generation, management and interpretation. The pharmaceutical industry would continue to lead on the numerous, essential tasks of drug development beyond target selection and validation including compound synthesis and screening, detailed mechanistic studies to elucidate mode of action, toxicology, pharmacokinetics, first-in-man studies and clinical trials. The intellectual property and commercial advantage would accrue from the agents developed, and from developing and evaluating the best drugs most efficiently against targets that have already been reliably deduced and validated. Since these activities would be concentrated on the correct therapeutic targets, and less likely misplaced, the risk of development failure should be reduced. This should stimulate a shift in R&D from the derivative to innovative, inspiring drug development for diseases that have previously been considered too risky to tackle. The benefits to society would come from containing drug development costs and expanding the therapeutic armamentarium against a broader range of diseases.

There would be additional benefits from such an effort. We have focused here mainly on genomic studies for matching targets with diseases (target identification). However, in related work (**see Appendix 1**) we (and others) have shown that the principle can also be used to anticipate the spectrum of effects of pharmacological action on biomarkers, surrogate and clinically relevant disease end-points. Mendelian randomisation for drug target validation has been used to accurately predict phase 3 trial outcomes, distinguish on- from off-target effects of drugs, correctly identify detailed biomarker profiles of therapeutic response, and to identify repurposing opportunities for licensed therapies. This underscores the view that such studies are not just useful for target identification but can also for inform drug development programmes from start to finish by indicating biomarkers of therapeutic response to measure in phase 1/2 clinical studies, and the relevant spectrum of clinical outcomes that should be ascertained in clinical trials. The incorporation of outcomes in clinical trials that are anticipated to be affected by pharmacological action on a particular target (*target*-*specific outcomes* of both efficacy and safety) would represent a departure from the current norm where end-points in a particular therapeutic area tend to be uniform regardless of the target being evaluated. Genetic information could also be useful for compound optimisation since the profile of biomarker effects of a SNP in a gene encoding a drug target should be those of a clean drug with no off-target actions ^84 104^. Where compounds are developed that have actions that are distinct from those observed in a genetic study, these may be off-target effects, and suggest that a more specific compound may need to be developed before the programme progresses. By the same principle, PheWAS would inform which clinical efficacy and safety end-points should be specified as outcomes in RCTs of compounds against a specified target. The spectrum of outcomes could differ from target to target, even for two targets being evaluated for the same primary disease indication. RCTs would need to be powered for both safety and efficacy outcomes, so that the balance between the benefits and any risk of target modification can be quantified before licensing. This should reduce the problem of mechanism-based side effects only emerging post marketing. This would also ensure that RCTs do not fail for failure to select the correct end-points, or because of the contamination of composite end-points (and thereby dilution of any treatment effect) by inclusion of outcomes that are unaffected by target modification.

*Delivery vehicle and funding:* The appropriate delivery vehicle for such an initiative requires careful consideration. It could be a form of social enterprise entrusted to create an open innovation platform where individual data is secured and protected, while aggregated data on genetic associations is shared, for the purpose of drug target identification and validation. Investment for the platform could come from a partnership of academic funders, healthcare and industrial sources with the knowledge generated helping all sectors and stakeholders.

Patients and healthcare providers would benefit from more efficient drug development, cost containment and, as a wider range of diseases is tackled, from access to a wider range of therapies. This could encourage government investment from healthcare, research, and business and innovation funding streams.

The biotech and digital technology sectors would benefit from a growing market for their technologies, while the pharmaceutical sector would benefit from what we believe will be greatly reduced failure rates in drug development.

The societal benefits that we believe will accrue may also be attractive to entrepreneurs looking to invest in a transformative social enterprise.

The leadership and oversight of such an endeavour would need to be trustworthy and accountable. It could come as a natural progression for academic medical centres that have established strong translational research programmes. In England, for example, these are funded by the National Institute of Health Research through Biomedical Research Centres (BRCs) formed of University / NHS partnerships, with the deep involvement of patients in their research activities. Increasingly, such centres are also establishing collaborative research activities and partnerships with industry, based on projects that are most likely to have patient benefit. Mature patient and public involvement activities, which underpin the work of all BRCs, could help identify and address patient and societal concerns, gauge enthusiasm for the proposal and, if accepted, help enrol patient champions for the project. Law Faculties in the academic sector, working with their counterparts in the healthcare systems and industry would also be well placed to develop solutions for legal, ethical and data protection issues that would undoubtedly arise.

Whatever the organisational structure, the outputs of the project – information on the correct drug targets for human diseases, and the outcomes relevant to perturbation of individual targets – would be made available without restriction, using an open access model. This would ensure target identification is pre-competitive, with any commercial advantage and intellectual property coming from other aspects of drug development.

### Conclusions

The fundamental problem in contemporary drug development has been the unreliability of target identification leading to low development success rates, inefficiency and escalating cost to healthcare users. Genomics now provides a tool to address the problem directly by accurate identification of proteins that both play a controlling role in a disease and which are amenable to targeting by drugs. Maximising the opportunities arising from this paradigm requires the wider use of genomics in the healthcare setting and with this, the active participation of healthcare users in drug development. The democratisation of drug development could have the consequence of reducing wasted investment, increasing value for investors and, eventually, reducing drug price inflation for healthcare providers. It might also provide the sorely needed stimulus for true drug development innovation, to the benefit of patients, health systems, business and society.

## Acknowledgements

ADH and HH are NIHR Senior Investigators and receive funding from the UCL Hospitals NIHR Biomedical Research Centre, the British Heart Foundation, Rosetrees Trust and Stoneygate Trust. JPO is an employee of Medicines Discovery Catapult, a UK non-profit aimed at supporting the discovery of novel medicines. Work at the Farr Institute of Health Informatics Research is funded by The Medical Research Council (K006584/1), in partnership with Arthritis Research UK, the British Heart Foundation, Cancer Research UK, the Economic and Social Research Council, the Engineering and Physical Sciences Research Council, the National Institute of Health Research, the National Institute for Social Care and Health Research (Welsh Assembly Government), the Chief Scientist Office (Scottish Government Health Directorates) and the Wellcome Trust. Work at the European Bioinformatics Institute is funded by Member States of the European Molecular Biology Laboratory.

---

| Orthodox drug development |  |  |  |  |  | Mendelian randomisation trials (MRT) |  |  |  |
| --- | --- | --- | --- | --- | --- | --- | --- | --- | --- |
| Drug target | Compound(s) evaluated | Developmental stage | Therapeutic area | Outcomes assessed in preclinical studies or RCTs of selective drug interventions | Findings from preclinical studies or RCTs of selective drug interventions | Encoding gene | Outcomes evaluated in MRTs | Findings from MRTs | Inferences drawn from comparison of the findings from preclinical studies or RCTs and MRT |
| **Cholesteryl ester transfer protein**[1] | Torcetrapib | Phase III | Cardiovascular disease | Blood lipids (total-, LDL-, and HDL cholesterol, triglycerides); blood pressure; CVD events | HDL-elevation, triglyceride and LDL-reduction. Unintended BP elevation. Unintended increase in CVD events | *CETP*[2] | Blood lipids (total-, LDL-, and HDL cholesterol, triglycerides); blood pressure | Associations with blood lipids consistent with effects in RCTs. No genetic association with BP. | Blood pressure elevating effect of torcetrapib is offtarget |
| **Hydroxy methyl (HMG)-coA reductase**[3] | Statins | Phase IV (post-marketing) | Cardiovascular disease | Blood lipid fractions, weight, type 2 diabetes risk | Statin treatment in RCTs linked to increased weight and risk of type 2 diabetes. | *HMGCR*[3] | Blood lipid fractions, anthropometric measures, glucose and insulin, type 2 diabetes risk | *HMGCR* SNPs associated with lower LDL-C, higher weight, fasting glucose and insulin, and type 2 diabetes risk | Increased risk of type 2 diabetes is an unintended on-target effect of statins mediated in part through weight gain |
| **Niemann-Pick C1-like 1** [4] | Ezetimibe | Phase III | Cardiovascular disease | LDL-cholesterol, cardiovascular death, non-fatal myocardial infarction, unstable angina requiring hospitalisation and revascularisation | Ezetimibe added to statins produces modest additional benefit in cardiovascular outcomes in patients following an acute coronary syndrome | *NPC1L1* [5] | Plasma lipid levels and risk of coronary heart disease. | Inactivating mutations in *NPC1L1* are associated with lower LDL-cholesterol and protection from myocardial infarction risk. | Niemann-Pick C1-like 1 is a validated target for LDL-cholesterol lowering and coronary heart disease prevention. |
| **Proprotein convertase subtilisin/kexin type 9 serine protease** [6] | Alirocumab, evolocumab | Phase II | Lipid lowering and cardiovascular disease | LDL-cholesterol | Alirocumab and evolocumab reduce LDL-cholesterol among patients with heterozygous familial or polygenic hypercholesterolaemia and reduce cardiovascular events in patients with or at high risk of cardiovascular disease | *PCSK9* [7] | LDL-cholesterol and risk of coronary heart disease | Inactivating mutations in *PCSK9* associated with reduced LDL-cholesterol and CHD risk | Proprotein convertase subtilisin/kexin type 9 serine protease is a validated target for LDL-cholesterol lowering and reduction in cardiovascular risk |
| **Glucagon-like peptide-1 receptor** [8] | Liraglutide | Phase III | Diabetes and cardiovascular disease | Death from cardiovascular causes, non-fatal myocardial infarction, or non-fatal stroke. | Liraglutide reduced risk of death from cardiovascular causes, nonfatal myocardial infarction, or nonfatal stroke among patients with type 2 diabetes mellitus | *GLP1R* [9] | Body weight, glycaemic traits, lipids, blood pressure, risk of type 2 diabetes and coronary heart disease | A low frequency, coding region missense variant in *GLP1R* is associated with lower fasting glucose, diabetes risk and risk of coronary heart disease. | *GLP1R* is a validated target for treatment of diabetes and reducing coronary heart disease risk |

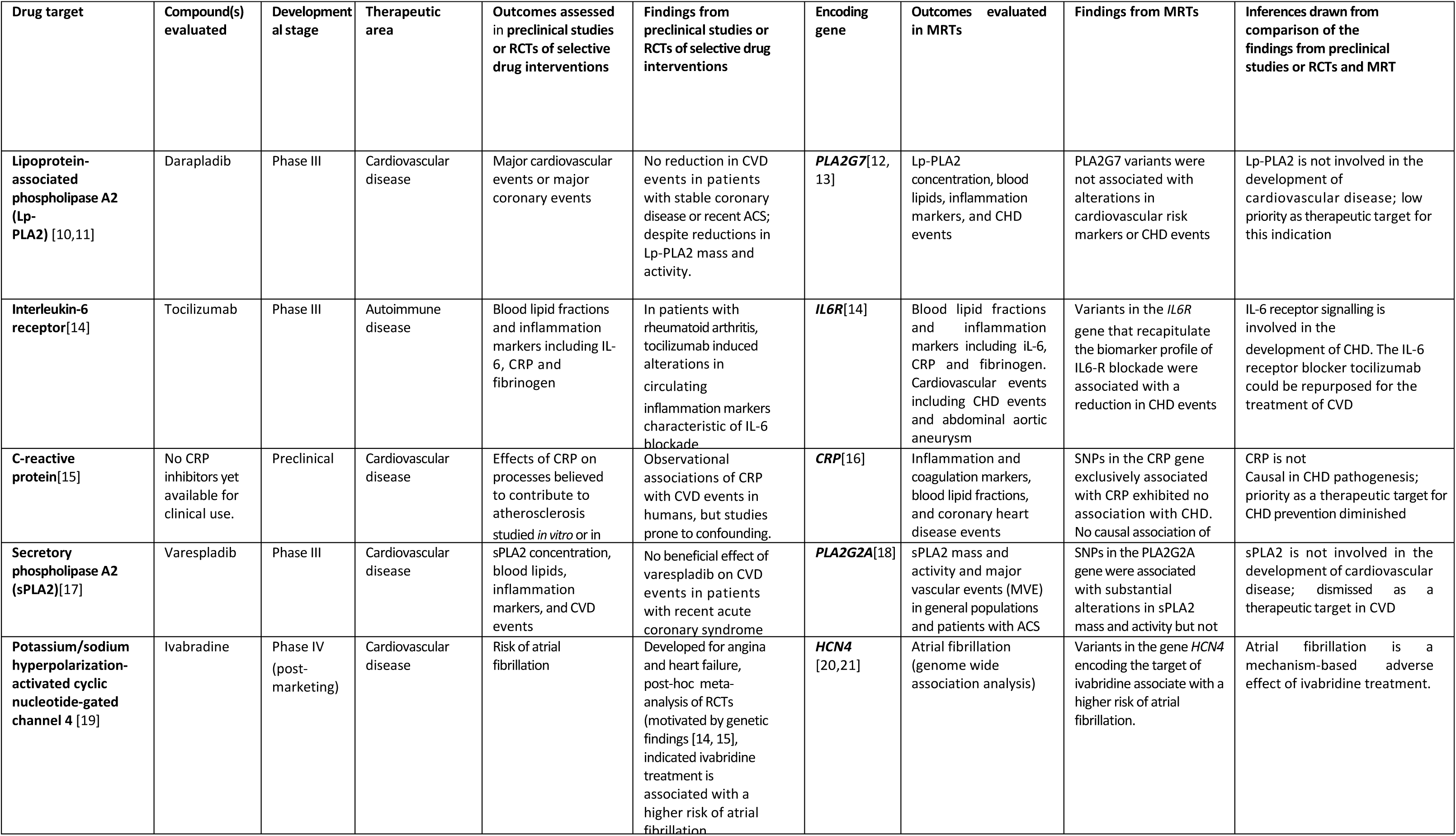

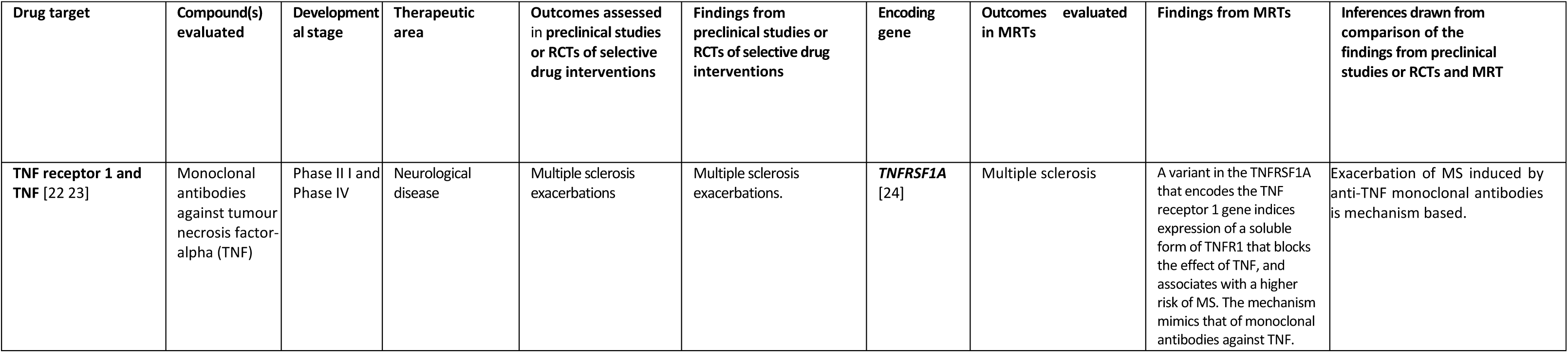

## Appendix 1. Comparison of the findings from orthodox randomised controlled trials or meta-analyses, and Mendelian randomisation trials of the corresponding therapeutic target

## Appendix 2

### Calculation of the probability of success for a company that initiates *N* parallel pre-clinical trials but will only pursue one of the signals to a further clinical trial

Suppose industry selects *N* targets at random_from a pool of *t* targets where only *c* targets are causal to the disease of interest. The *N* pre-clinical programmes will generate a number of positive signals of which the company will select **only** to progress to clinical phase following which there will be a licensing success (if the signal comes from a true target) or failure if the preclinical signal is a false positive. To calculate the probability of eventual licensing success we consider a situation where many companies repeat an experiment involving *N* preclinical programmes only pursuing only one of the positive signals to a phase 3 clinical trial, and then calculating what proportion of such trials will result in a licensing success.

1. We first calculate the probability of having *A* causal targets among the *N* targets selected at random from the pool of *t* possible targets. Each company will select a different number by chance (*A = 0,1,2,3*…) with the probabilities of each following the hypergeometric distribution:

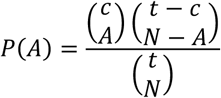 So, if *t* = 4000 with *c* = 20, and we run *N* = 20 preclinical trials then:

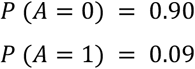
2. We next calculate the probability of generating true signals (*St*) and false signals (Sf): The *A* causal targets in the *N* programmes can generate from 0 to *A* signals (*St* = 0,1…*A*), while the non-causal target can generate from 0 to *N* – *A* signals (*Sf* = 0,1,2…*N* – *A*). Each of these probabilities follow a binomial distribution independent from each other:

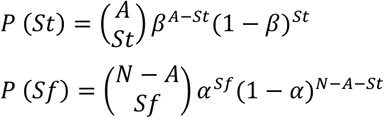 Where (1 – *β*) and *α* are the probabilities that a causal and non-causal target will produce a signal respectively. The two probabilities being independent, the probability of a particular combination of signals from causal and non-causal targets is the product of the separate probabilities: *P*(*St*, *Sf*) = *P*(*St*) *x P*(*Sf*). For example, the probability that, in a given repetition the causal targets produce *2* signals and the non-causal targets produce three signals is *P*(*St* = 2, *Sf* = 3) = *P*(*St* = 2) × *P*(*Sf* = 3)
3. The probability of selecting a real target among a combination of true and false signals (*St*,*Sf*) is given by the proportion of true signals: *St* / (*St* + *Sf*)

Thus, for a given *N*, *c* and *t*, the final probability of licensing success across all possible values of *A*, *St* and *Sf* is:

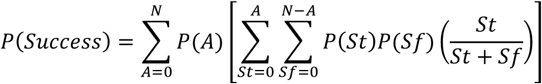

## Supplementary tables

**Table S1.** Expected number of licensed drug targets rediscovered (***E_T_***) by 200 hypothetical GWAS of diseases with at least one licensed drug based on a range of plausible values of the power (1 – *β*) to detect each genetic locus encoding a licensed drug target, and a range of plausible values for the average number of licensed drug targets per disease. (See text for further details)

| Number of licensed drug targets per disease | Power ($1 - \beta$) | $E_T$ (SD) |
| --- | --- | --- |
| 1 | 0.6 | 120 (7) |
| 1 | 0.8 | 160 (6) |
| 1 | 0.9 | 180 (4) |
| 3 | 0.6 | 360 (12) |
| 3 | 0.8 | 480 (10) |
| 3 | 0.9 | 540 (7) |
| 5 | 0.6 | 600 (15) |
| 5 | 0.8 | 800 (13) |
| 5 | 0.9 | 900 (9) |
| 10 | 0.6 | 1200 (22) |
| 10 | 0.8 | 1600 (18) |
| 10 | 0.9 | 1800 (13) |

**Table S2.** Effect of varying estimates of the number of causative genes per disease (*C*), and the number of diseases (***N_D_***) on the probability of selecting a causal gene-disease pair (***γ_C_***); the probability of selecting a causal, druggable, gene-disease pair (***γ_CT_***); and the number diseases influenced by any one gene (or encoded protein) (***E_D_***.). Estimates assume 20,000 protein-coding genes.

| $C$ | $N_D$ | $\gamma_C$ | $\gamma_{CT}$ | $E_D$ |
| --- | --- | --- | --- | --- |
| 10 | 2500 | 0.0005 | 0.0001 | 1.25 |
| 10 | 5000 | 0.0005 | 0.0001 | 2.5 |
| 10 | 10000 | 0.0005 | 0.0001 | 5 |
| 100 | 2500 | 0.005 | 0.001 | 12.5 |
| 100 | 5000 | 0.005 | 0.001 | 25 |
| 100 | 10000 | 0.005 | 0.001 | 50 |
| 1000 | 2500 | 0.05 | 0.01 | 125 |
| 1000 | 5000 | 0.05 | 0.01 | 250 |
| 1000 | 10000 | 0.05 | 0.01 | 500 |

**Table S3:** Expected yield of causal druggable targets from orthodox (non-genomic) preclinical programmes according to the number of causal targets for each disease and whether the sampling frame is the whole genome or the druggable genome.

| Number of programmes | Number of causal, druggable targets per disease | Number of targets in sampling frame | Expected number (*SD*) of causal, druggable targets among all programmes | Number causal druggable targets detected ($1 - \beta = 0.8$) | Number of non-relevant targets declared positive ($\alpha = 0.05$) |
| --- | --- | --- | --- | --- | --- |
| 10 | 20 | 20,000 | 0.01 (0.07) | 0.008 | 0.49 |
| 20 | 20 | 20,000 | 0.02 (0.1) | 0.016 | 1.0 |
| 50 | 20 | 20,000 | 0.05 (0.2) | 0.04 | 2.5 |
| 100 | 20 | 20,000 | 0.1 (0.2) | 0.08 | 5.0 |
| 200 | 20 | 20,000 | 0.2 (0.3) | 0.16 | 10.0 |
| 10 | 20 | 4,000 | 0.05 (0.2) | 0.04 | 5.0 |
| 20 | 20 | 4,000 | 0.1 (0.2) | 0.08 | 1.0 |
| 50 | 20 | 4,000 | 0.25 (0.4) | 0.2 | 2.5 |
| 100 | 20 | 4,000 | 0.5 (0.5) | 0.4 | 5.0 |
| 200 | 20 | 4,000 | 1 (0.7) | 0.8 | 10.0 |
| 10 | 200 | 20,000 | 0.1 (0.2) | 0.08 | 0.5 |
| 20 | 200 | 20,000 | 0.2 (0.3) | 0.16 | 1.0 |
| 50 | 200 | 20,000 | 0.5 (0.5) | 0.4 | 2.5 |
| 100 | 200 | 20,000 | 1 (0.7) | 0.8 | 5.0 |
| 200 | 200 | 20,000 | 2 (1) | 1.6 | 10.0 |
| 10 | 200 | 4,000 | 0.5 (0.5) | 0.4 | 0.5 |
| 20 | 200 | 4,000 | 1 (1) | 0.8 | 1.0 |
| 50 | 200 | 4,000 | 2.5 (1) | 2 | 2.4 |
| 100 | 200 | 4,000 | 5 (1) | 4 | 4.8 |
| 200 | 200 | 4,000 | 10 (2) | 8 | 9.5 |

**Table S4.** Number of drug development programmes (*N*) that to be pursued in parallel to have a probability (*P*) of at least one development success. Analyses are based on either 90% or 50% (evens) probability of at least one developmental success, and a range of development success rates (*p*) starting with the currently observed industry wide average success rate of 0.01 (See text for details)

| $P$ ($\geq 1$ success) in $N$ programmes | Within-programme developmental success rate ($P_S$) | Number of parallel programmes ($N$) |
| --- | --- | --- |
| 0.9 | 0.01 | 229 |
| 0.9 | 0.02 | 114 |
| 0.9 | 0.1 | 22 |
| 0.9 | 0.2 | 10 |
| 0.9 | 0.5 | 3 |
| 0.5 | 0.01 | 69 |
| 0.5 | 0.02 | 34 |
| 0.5 | 0.1 | 7 |
| 0.5 | 0.2 | 3 |
| 0.5 | 0.5 | 1 |

**Table S5.** Expected number of true and false positives in parallel drug development programmes based on a sample of targets drawn from all or part of the druggable genome based on orthodox preclinical experiments designed with (1 – *β*) = 0.8 and *α* = 0.05 (left hand panel). Probability of eventual drug development success taking forward one positive preclinical programme to clinical phase (right hand panel). (See text for further details)

| Targets in sampling frame | True causal genes | Number of parallel development programmes pursued | Expected true positives in sample | Expected false positives in sample | Positive programmes are exclusively true positives | Positive programmes are a mixture of true and false positives | No positive programmes | Positive programmes are exclusively false positives | Overall probability of a development success |
| --- | --- | --- | --- | --- | --- | --- | --- | --- | --- |
| 4000 | 20 | 20 | 0.08 | 1.00 | 2.9% | 4.8% | 33.1% | 59.2% | 5.0% |
| 2000 | 20 | 20 | 0.16 | 0.99 | 5.7% | 9.2% | 30.6% | 54.5% | 9.7% |
| 1000 | 20 | 20 | 0.32 | 0.98 | 10.6% | 17.2% | 26.0% | 46.2% | 18.3% |
| 200 | 20 | 20 | 1.60 | 0.90 | 33.3% | 49.0% | 6.5% | 11.1% | 60.4% |
| 4000 | 20 | 50 | 0.20 | 2.49 | 1.5% | 16.8% | 6.3% | 75.5% | 7.0% |
| 2000 | 20 | 50 | 0.40 | 2.48 | 2.7% | 30.6% | 5.2% | 61.5% | 13.3% |
| 1000 | 20 | 50 | 0.80 | 2.45 | 4.7% | 51.4% | 3.4% | 40.5% | 24.0% |
| 200 | 20 | 50 | 4.00 | 2.25 | 9.9% | 89.1% | 0.1% | 0.9% | 64.6% |
| 4000 | 20 | 200 | 0.80 | 9.95 | 0.0% | 55.9% | 0.0% | 44.1% | 7.5% |
| 2000 | 20 | 200 | 1.60 | 9.90 | 0.0% | 81.2% | 0.0% | 18.7% | 14.0% |
| 1000 | 20 | 200 | 3.20 | 9.80 | 0.0% | 97.0% | 0.0% | 3.0% | 24.8% |
| 200 | 20 | 200 | 16.00 | 9.00 | 0.0% | 100.0% | 0.0% | 0.0% | 64.7% |

## Supplementary figures

**Figure S1a.**
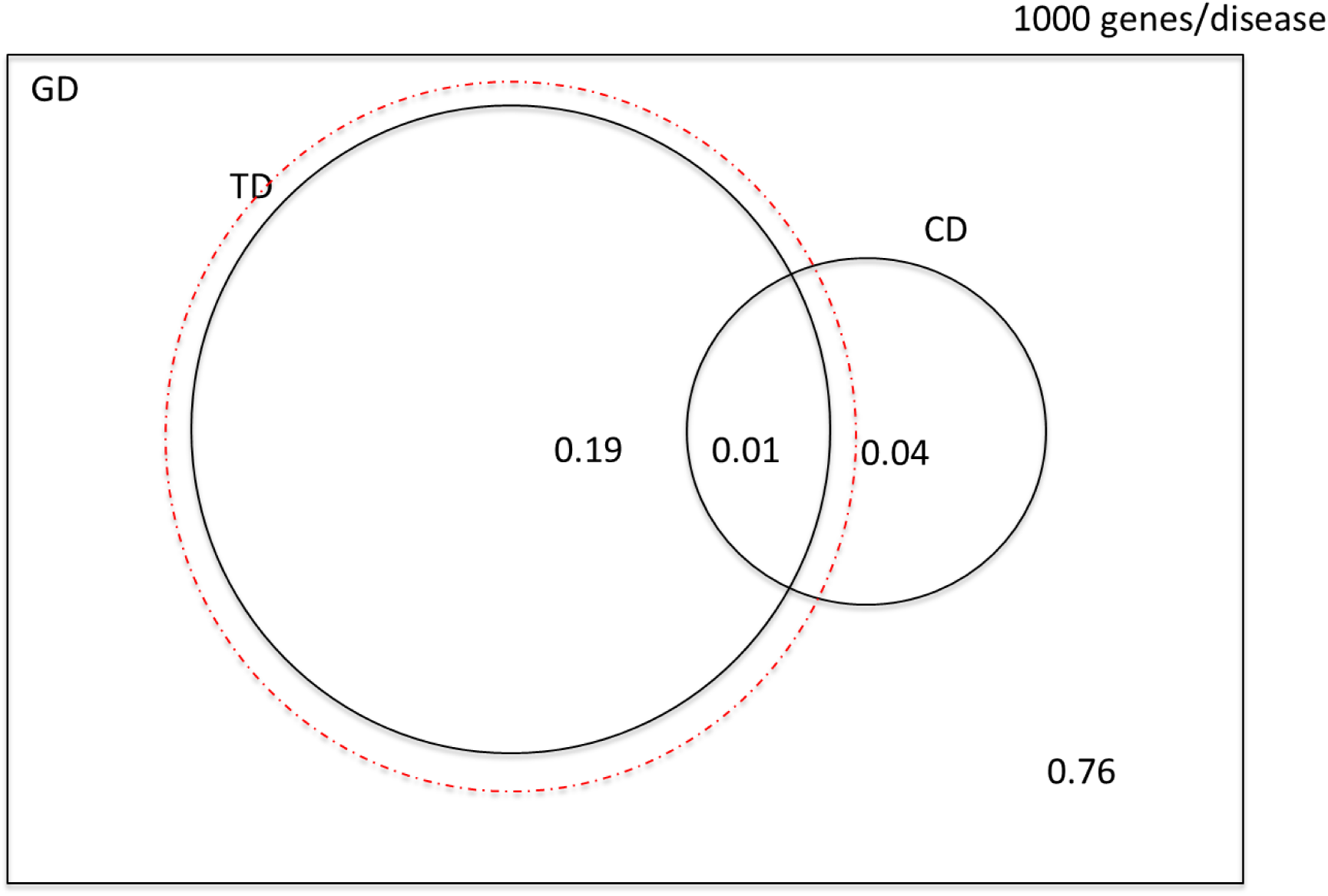
Venn diagram illustrating the probabilities of selecting a causal, druggable gene-disease pair (*CD* ∩ *TD*), a druggable gene disease pair (*TD*) and a causal, gene disease pair (*CD*) from a sample space of 200 *x* 10^6^ gene disease pairings, 1000 causal genes per disease and 4000 druggable genes from the 20,000 in the genome. The dashed red circle encloses a probability space restricted to druggable genes. (Not to scale).

**Figure S1b.**
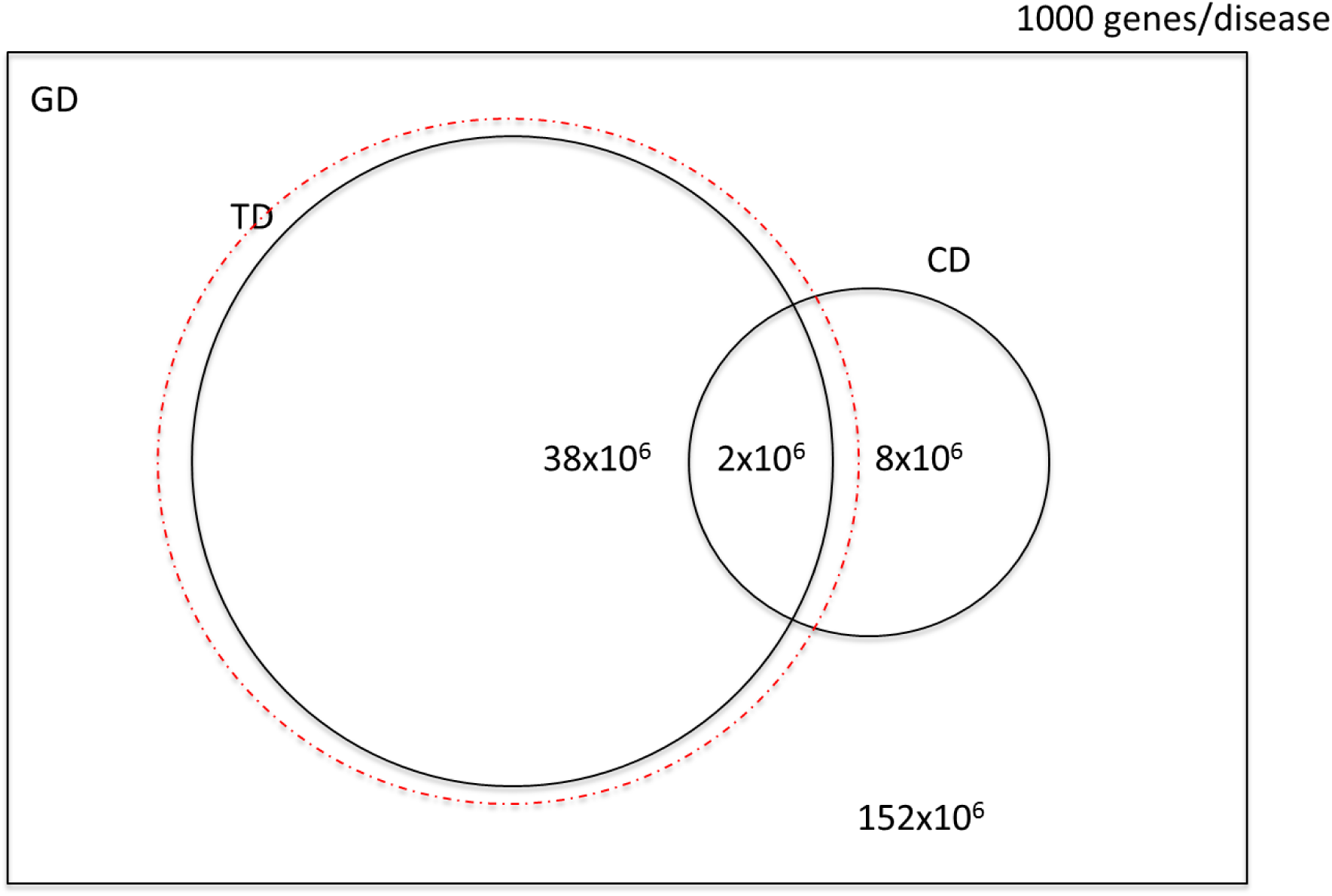
Venn diagram illustrating the number of causal, druggable gene-disease pairs (*CD* ∩ *TD*), druggable gene disease pairs (*TD*) and causal gene disease pairs (*CD*) from a sample space of 200 *x* 10^6^ gene disease pairings, 1000 causal genes per disease and 4000 druggable genes from the 20,000 in the genome. The dashed red circle encloses a probability space restricted to druggable genes. (Not to scale).

**Figure S2a.**
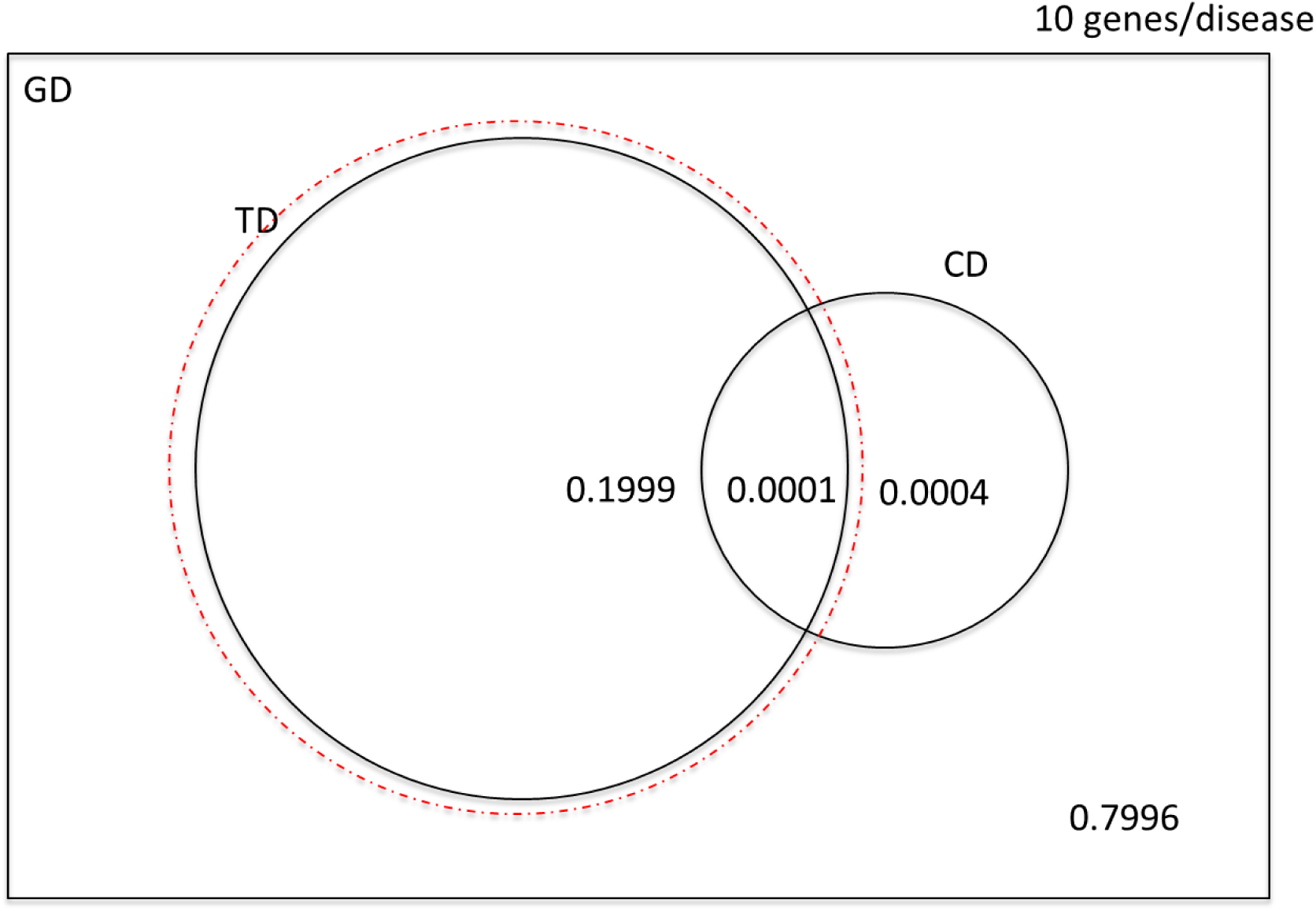
Venn diagram illustrating the probabilities of selecting a causal, druggable gene-disease pair (*CD* ∩ *TD*), a druggable gene disease pair (*TD*) and a causal, gene disease pair (*CD*) from a sample space of 200 *x* 10^6^ gene disease pairings, 10 causal genes per disease and 4000 druggable genes from the 20,000 in the genome. The dashed red circle encloses a probability space restricted to druggable genes. (Not to scale).

**Figure S2b.**
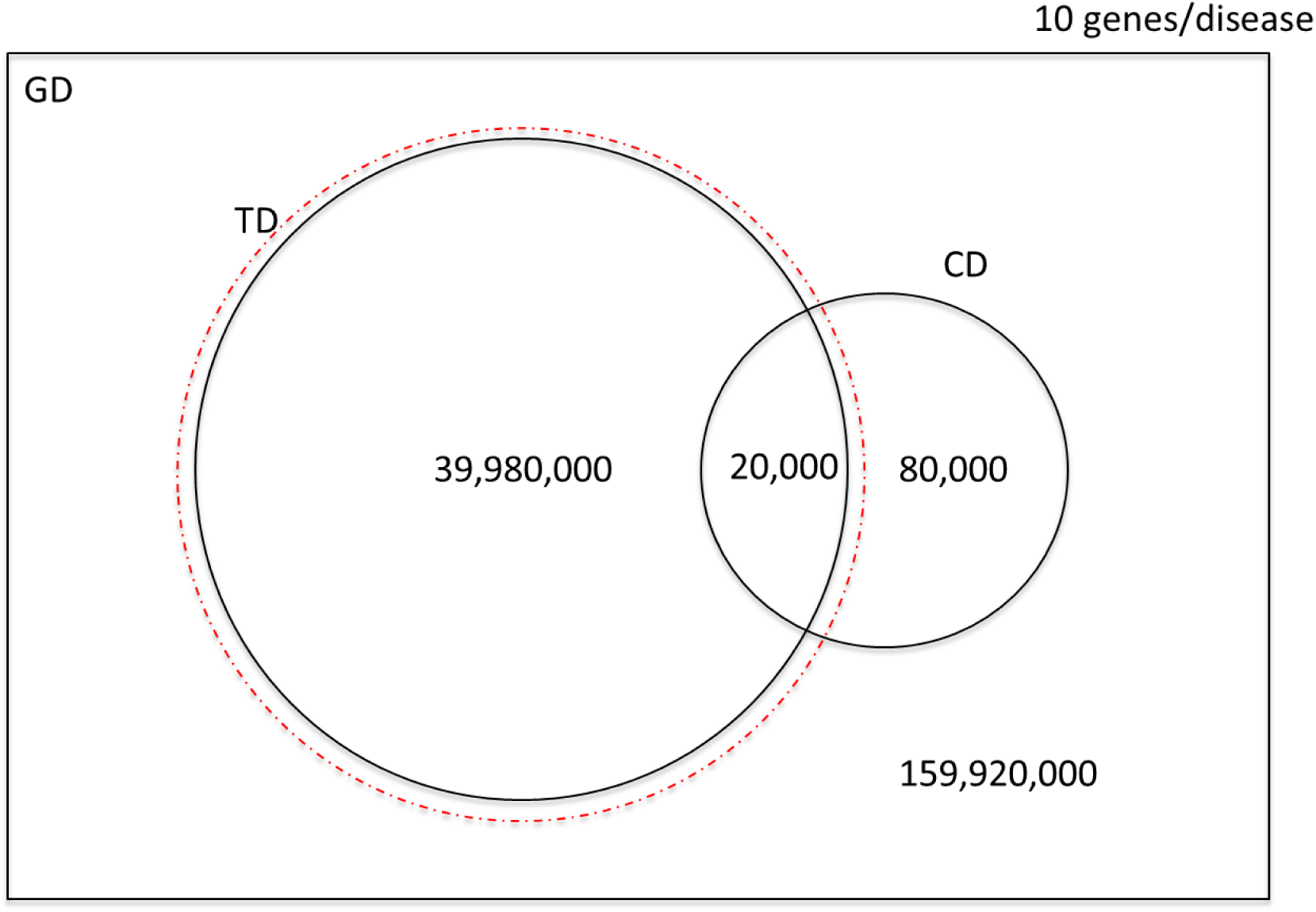
Venn diagram illustrating the number of causal, druggable gene-disease pairs (*CD* ∩ *TD*), druggable gene disease pairs (*TD*) and causal gene disease pairs (*CD*) from a sample space of 200 *x* 10^6^ gene disease pairings, 10 causal genes per disease and 4000 druggable genes from the 20,000 in the genome. The dashed red circle encloses a probability space restricted to druggable genes. (Not to scale).

## Footnotes

a We exclude drug targets encoded by the abnormal genome of cancer cells as well as antimicrobials, which typically target proteins encoded in the genomes of pathogens. For further discussion, see Part 4

